# Functional profiling of *KCNE1* variants informs population carrier frequency of Jervell and Lange-Nielsen syndrome type 2

**DOI:** 10.1101/2025.03.28.646046

**Authors:** Carlos G. Vanoye, Reshma R. Desai, Jordan D. John, Steven C. Hoffman, Nicolas Fink, Yue Zhang, Omkar G. Venkatesh, Jonathan Roe, Sneha Adusumilli, Nirvani P. Jairam, Charles R. Sanders, Adam S. Gordon, Alfred L. George

## Abstract

Congenital long-QT syndrome (LQTS) is most often associated with pathogenic variants in *KCNQ1* encoding the pore-forming voltage-gated potassium channel subunit of the slow delayed rectifier current (*I*_Ks_). Generation of *I*_Ks_ requires assembly of KCNQ1 with an auxiliary subunit encoded by *KCNE1*, which is also associated with LQTS but causality of autosomal dominant disease is disputed. By contrast, *KCNE1* is an accepted cause of recessive type 2 Jervell and Lange-Nielson syndrome (JLN2). The functional consequences of most *KCNE1* variants have not been determined and the population prevalence of JLN2 is unknown. *Methods*: We determined the functional properties of 95 *KCNE1* variants co-expressed with KCNQ1 in heterologous cells using high-throughput voltage-clamp recording. Experiments were conducted with each KCNE1 variant expressed in the homozygous state and then a subset was studied in the heterozygous state. The carrier frequency of JLN2 was estimated by considering the population prevalence of dysfunctional variants. *Results*: There is substantial overlap between disease-associated and population KCNE1 variants. When examined in the homozygous state, 68 KCNE1 variants exhibited significant differences in at least one functional property compared to WT KCNE1, whereas 27 variants did not significantly affect function. Most dysfunctional variants exhibited loss-of-function properties. We observed no evidence of dominant-negative effects. Most variants were scored as variants of uncertain significance (VUS) and inclusion of functional data resulted in revised classifications for only 14 variants. The population carrier frequency of JLN2 was calculated as 1 in 1034. Peak current density and activation voltage-dependence but no other biophysical properties were correlated with findings from a mutational scan of KCNE1. *Conclusions*: Among 95 disease-associated or population KCNE1 variants, many exhibit abnormal functional properties but there was no evidence of dominant-negative behaviors. Using functional data, we inferred a population carrier frequency for recessive JLN2. This work helps clarify the pathogenicity of KCNE1 variants.

## Introduction

Congenital cardiac arrhythmia susceptibility including monogenic etiologies of the long-QT syndrome (LQTS) are important and recognizable causes of sudden death in the young.^1,2^ Autosomal dominant and recessive LQTS are most often associated with pathogenic variants in genes encoding voltage-gated potassium, sodium and calcium channels or channel auxiliary subunits or specific modulatory proteins.^3^ Determining the functional consequence of LQTS-associated variants helps define disease mechanisms at the molecular level and can assist in discriminating pathogenic from benign protein changes.

Variants in *KCNQ1* are the most frequently discovered cause of autosomal dominant LQTS (designated as the LQT1 subtype).^4^ This gene encodes the KCNQ1 (K_V_7.1) voltage-gated potassium channel, which is the pore-forming subunit responsible for generating the slow delayed rectifier current (*I*_Ks_), a major repolarizing current in cardiac myocytes. The generation of *I*_Ks_ depends on a molecular partnership between KCNQ1 and an auxiliary subunit encoded by *KCNE1*.^5,6^ Heterozygous *KCNE1* variants are associated with genetic LQTS in sporadic cases and small families (designated LQT5), but an international LQTS gene curation working group of the Clinical Genome Resource (ClinGen) found limited evidence supporting disease causality.^7^ However, *KCNE1* is a recognized cause of the autosomal recessive Jervell and Lange-Nielson Syndrome type 2 (JLN2) featuring LQTS combined with congenital sensorineural hearing impairment.^8,9^ JLN2 is a rare disorder with an uncertain population prevalence.

An international multicenter study evaluated the clinical features and electrocardiographic penetrance of *KCNE1* variants in several genotype-positive individuals including 229 who were heterozygous and 19 who were diagnosed with JLN2.^10^ This study also noted that many of the 32 unique disease-associated *KCNE1* variants among the study population were also found in individuals from the Genome Aggregation Database (gnomAD).^11^ Penetrance and arrhythmia event frequencies were low in both groups but experimental evidence of deleterious consequences was lacking for the majority of the variants raising the possibility that some non-pathogenic missense variants were falsely associated with LQTS in some heterozygous probands.^12^ Additional experimental evidence may assist in determining pathogenicity for *KCNE1* variants.

The most widely used approach for determining the functional consequences of an ion channel variant involves heterologous expression of a recombinant channel coupled with voltage-clamp electrophysiological recording. The typical experimental approach (e.g., manual patch clamp recording) is slow, tedious and has limited throughput. Automated patch clamp recording enables much higher throughput and enables study of large numbers of channel variants including those in LQTS genes.^13-17^ We previously optimized automated patch clamp recording methods to study dozens of *KCNQ1* variants, which are also suitable for examining the functional effects of *KCNE1* variants.^13,18^ Beyond electrophysiological studies, other approaches (e.g., deep mutational scanning) for screening large numbers of engineered variants have been applied to specific ion channels or regulatory proteins.^17,19-21^ Muhammad and colleagues reported use of high-throughput variant effect mapping of KCNE1 combining a cell viability assay with a flow-cytometry enabled cell surface expression analysis to assign functional classes to the majority of all possible missense KCNE1 variants.^21^ How the findings from such high-throughput screening assays correlate with detailed electrophysiological analyses is not well studied.

In this study, we determined the functional consequences of a large number of *KCNE1* variants using high-throughput voltage-clamp recording. These data provide detailed biophysical properties of disease-associated and population variants in this gene. We assessed the impact of incorporating variant function on assignment of pathogenicity. We also used these data to estimate the population carrier frequency of JLN2, and to benchmark results from a deep mutational scan of KCNE1.

## Methods

### Plasmids and mutagenesis

Full length cDNA encoding WT KCNQ1 (GenBank accession number AF000571) was engineered in the mammalian expression vector pIRES-CyOFP1 (pIRES-CyOFP1-KCNQ1, AddGene #173162). Human KCNE1 cDNA (GenBank accession number EF514883) was engineered in the mammalian expression vector pIRES2-EGFP and included a 53 base pair cassette (GGATCCACCGGTCATCCACGCCTGAGATCTAGACAAACTTCCTAGACTGCATG) inserted immediately after the stop codon at a *Bam*HI site to facilitate cloning (pIRES-EGFP-GA-KCNE1, AddGene #173160). Both vectors enabled co-expression of untagged channel subunits with either orange or green fluorescent proteins as a means for tracking successful cell transfection. KCNE1 variants were introduced into the WT coding sequence using one of two methods. For some variants, the complete KCNE1 open reading frame with individual variants was synthesized using a Synthetic Genomics BioXp 3200 workstation (SGI-DNA, La Jolla, CA) and cloned into pIRES-EGFP-GA-KCNE1 using Gibson assembly. Other KCNE1 variants were generated by site-directed PCR mutagenesis using Q5 Hot Start High Fidelity DNA polymerase (New England Biolabs, Ipswich, MA) as previously described.^13^ Mutagenic primers for PCR mutagenesis were designed using custom software (Q5 primer designer; available at https://prism.northwestern.edu/records/tcsgw-z4295), and are presented in **Supplemental Table S1**. Complete sequence of each variant plasmid was performed with nanopore sequencing (Primordium Labs, Arcadia, CA) and analyzed using a custom multiple sequence alignment tool (MuSIC; available at https://prism.northwestern.edu/records/h6hc6-n0j20).

### Cell culture and electroporation

CHO-K1 cells (CCL-61, American Type Culture Collection, Manassas, VA) or CHO-K1 cells constitutively expressing human KCNE1 (designated CHO-KCNE1 cells)^13^ were used. Cells were grown in F-12 nutrient medium (Thermo Fisher Scientific, Waltham, MA) supplemented with 10% fetal bovine serum (Atlanta Biologicals, Flowery Branch, GA), penicillin (50 units/mL), streptomycin (50 μg/mL) at 37°C in 5% CO_2_ and maintained under selection with hygromycin B (600 μg/mL). Plasmids encoding WT KCNQ1 and WT or mutant KCNE1 were transiently expressed by electroporation using the MaxCyte STX system (MaxCyte Inc., Rockville, MD).^14^ Cells were grown to 70-80% confluence then harvested using 0.25% trypsin (Thermo Fisher Scientific, Waltham, MA). A 500 µl aliquot of cell suspension was used to determine cell number and viability using an automated cell counter (ViCell, Beckman Coulter, Brea, CA). Remaining cells were collected by centrifugation at 160 x g for 4 minutes, washed with electroporation buffer (EBR100, MaxCyte Inc.), and re-suspended in electroporation buffer at a density of 100 x 10^6^ viable cells/ml. Each electroporation was performed using 100 µl of cell suspension. CHO-K1 or CHO-KCNE1 cells were electroporated with 10 µg of WT KCNQ1 cDNA and 20 µg of WT or mutant KCNE1. The DNA-cell suspension mix was transferred to an OC-100×2 processing assembly (MaxCyte Inc.) and electroporated using the preset CHO-PE (CHO Protein Expression) protocol. Immediately after electroporation, 10 µl of recombinant human deoxyribonuclease I (dornase alpha, Pulmozyme^®^, Genentech, Inc., San Francisco, CA) was added to the DNA-cell suspension and the entire mixture was transferred to a 35 mm tissue culture dish for a 30 min incubation at 37°C in 5% CO_2_. Following incubation, cells were gently suspended in culture media, transferred to a T75 tissue culture flask and grown for 24 hours at 37°C in 5% CO_2_. Following incubation, cells were harvested, counted, transfection efficiency determined by flow cytometry (see below), and then frozen in 1 ml aliquots at 1.8 x 10^6^ viable cells/ml in liquid nitrogen until they were used in experiments. Transfection efficiency was evaluated before freezing using a benchtop flow cytometer (CytoFLEX, Beckman Coulter). Forward scatter (FSC), side scatter (SSC), and orange and green fluorescence were recorded. FSC and SSC were used to gate single viable cells and to eliminate doublets, dead cells and debris. Ten thousand events were recorded for each sample. Non-electroporated cells were assayed as a control for all parameters and used to set the gates for each experiment. The percentage of fluorescent cells was determined from the gated population.

### Electrophysiology

The day before automated patch clamp recording, electroporated cells were thawed, plated and incubated for 10 hours at 37°C in 5% CO_2_. The cells were then grown overnight at 28°C in 5% CO_2_. Prior to experiments, cells were passaged using 0.25% trypsin in cell culture media. Cell aliquots (500 µl) were used to determine cell number and viability by automated cell counting and transfection efficiency by flow cytometry. Cells were then diluted to 200,000 cells/ml with external bath solution (see below) and allowed to recover 60 minutes at 15°C while shaking on a rotating platform at 200 rpm. Automated patch clamp experiments were performed using the Syncropatch 768 PE platform (Nanion Technologies, Munich, Germany). Single-hole, 384-well medium resistance (4-5 MΩ) or S-Type (2.5-3.5 MΩ) recording chips were used. Pulse generation and data collection were carried out with PatchController384 V.1.3.0 and DataController384 V1.2.1 software (Nanion Technologies). Whole-cell currents were recorded at room temperature in the whole-cell configuration, filtered at 3 kHz and acquired at 10 kHz. The access resistance and apparent membrane capacitance were estimated using built-in protocols. The external bath solution contained: 140 mM NaCl, 4 mM KCl, 2 mM CaCl_2_, 1 mM MgCl_2_, 10 mM HEPES, 5 mM glucose, pH 7.4. The internal solution contained: 60 mM KF, 50 mM KCl, 10 mM NaCl, 10 mM HEPES, 10 mM EGTA, 2 mM ATP-Mg, pH 7.2. Whole-cell currents were elicited from a holding potential of -80 mV using 2000 ms depolarizing pulses (from -80 mV to +60 mV in +10mV steps, every 10 secs) followed by a 2000 ms step to - 30 mV to analyze tail currents and channel deactivation rate. Non-specific currents were eliminated by recording whole-cell currents before and after addition of the *I*_Ks_ selective blocker JNJ-303 (2 μM; Tocris Bioscience, Minneapolis, MN).^22^ Only JNJ-303 sensitive currents and recordings meeting the following criteria were used in data analysis: seal resistance ≥ 0.5 GΩ, series resistance ≤ 20 MΩ, capacitance ≥ 1 pF, voltage-clamp stability (defined as the standard error for the baseline current measured at the holding potential for all test pulses being <10% of the mean baseline current).

Data were analyzed and plotted using DataController384 V1.8 (Nanion Technologies), Excel (Microsoft Office 2013), SigmaPlot 2000 (Systat Software, Inc.) and Prism 8 (GraphPad Software) software packages. Additional custom semi-automated data handling routines were used for rapid analysis of current density and voltage-dependence of activation. Whole-cell currents were normalized for membrane capacitance and results expressed as mean ± 95% confidence interval (95% CI). Peak currents were recorded at 1990 ms after the start of the voltage pulse, while tail currents were measured 10 ms after changing the membrane potential to -30 mV. The voltage-dependence of activation was calculated by fitting the normalized G-V curves with a Boltzmann function (tail currents measured at -30 mV). The voltage-dependence of activation was determined only for cells with mean current density greater than the background current amplitude. The rate of channel deactivation was determined by fitting tail currents pre-activated with the +60 mV pulse to a single exponential function. Typical experiments compared five variants to the WT channels assayed on the same plate with up to 64 replicate recordings. Properties of each variant are presented relative to the WT channel assayed in parallel as percent current density measured at +60 mV, difference (Δ) in voltage-dependence of activation V_½_, and the ratio of activation or deactivation time-constants (variant / WT τ_act,_ or τ_deact_). The number of cells used for each experimental condition is given in supplemental datasets.

Statistical analysis was performed using one-way analysis of variance (3 or more groups) and *P* ≤ 0.02 was considered significant based on a Bonferroni correction for typical experiments in which 5 variants were measured in parallel with WT.

### Classification of *KCNE1* variants

*KCNE1* variants were classified as benign (B), likely benign (LB), variant of uncertain significance (VUS), likely pathogenic (LP), or pathogenic (P) in accordance with the American College of Medical Genetics and Genomics (ACMG) guidelines.^23^ Six reviewers (J.J., S.H., N.F., Y.Z., O.V., J.R.) independently classified each variant based on existing evidence and then arrived at a consensus. ACMG criteria that could be applied for one or more variants included PVS1, PM2, PP1, PP3 and BP4. In addition to these criteria, we also used results from our functional assay to incorporate the relevant PS3 criterion under three scenarios: 1) classification of variants without applying PS3; 2) classification of variants with PS3 determined from our experiments demonstrating loss-of-function (see below); and 3) classification of variants using PS3 at varying strengths to reflect the full spectrum of electrophysiological phenotypes (see below).

Evidence pertaining to *KCNE1* missense variants was gathered using multiple sources including published literature, ClinVar (https://www.ncbi.nlm.nih.gov/clinvar),^24^ InterVar (http://wintervar.wglab.org),^25^ gnomAD v.2.1.1 (https://gnomad.broadinstitute.org/),^11^ and ENSEMBL (http://www.ensembl.org).^26^ The PVS1 criterion, reflecting loss-of-function, was applied for any KCNE1 variant with deletion of more than the last 10% of the protein product consistent with Clinical Genome Resource (ClinGen) Sequence Variant Information Workgroup.^27^ For deletions within the final 13 residues of the 129 amino acid protein (i.e., those that may escape nonsense mediated mRNA decay), PVS1 was applied only if loss-of-function was demonstrated by our functional study. The PM2 allele frequency criteria was applied if the highest single population allele frequency in gnomAD (Grpmax) did not exceed 0.001. The PP1 co-segregation criterion was applied at different strengths according to the number of informative meiosis among published family data as proposed by Jarvik and Browning.^28^ Published pedigrees were only included in the PP1 calculation if they confirmed co-segregation of disease phenotype with a heterozygous KCNE1 variant, if genotypes were confirmed in all family members, if no more than one KCNE1 variant was present in affected family members, and if no KCNQ1 variants were present. The PP3 and BP4 criteria, reflecting *in silico* evidence, were derived from prediction scores generated by SIFT (https://sift.bii.a-star.edu.sg/)^29^ and PolyPhen-2 (http://genetics.bwh.harvard.edu/pph2/).^30^ PP3 or BP4 was assigned to a variant if SIFT and PolyPhen scores were concordant (deleterious or probably damaging for PP3; tolerated or benign for BP4), and neither were assigned if SIFT and PolyPhen2 were discordant.

The PS3 criterion was applied based on current density measured in our study relative to WT KCNE1. Functional data from previously published literature were not used. Variants exhibiting mild (<25%), moderate (25-74%) or severe (≥ 75%) reductions in current density in the homozygous state relative to KCNE1 WT were assigned PS3 with evidence strengths of supporting (PS3[P]), moderate (PS3[M]), or strong (PS3[S]), respectively. In an expanded analysis, PS3(M) was applied for variants exhibiting >150% current density relative to KCNE1 WT consistent with gain-of-function, and BS3[P] was applied if a variant exhibited 100 ± 10% current density relative to KCNE1 WT consistent with normal to near-normal activity.

### Estimation of JLN2 carrier frequency

To estimate the population carrier frequency for JLN2, we applied principles of Hardy-Weinberg equilibrium to publicly available, population-scale allele frequency data, adapting a previously described method.^31,32^ For this analysis, we used functional criteria to define a carrier as any individual who is heterozygous for a variant generating current density less than 50% of WT (**Supplemental Table S2**). We extracted the global counts of homozygous and heterozygous individuals for these specific variants from gnomAD, v2.1.1 (original source of population variants we studied), along with the total count of chromosomes assessed for each of these variants. Based on the rarity of these variants, we assumed linkage equilibrium among them – that is, we assumed that individuals are unlikely to carry a haplotype with more than one of these variants in *cis*. We therefore calculated carrier rate (2pq in the Hardy-Weinberg equilibrium) according to the following equation:

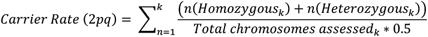

in which *k* is the number of variants with current density less than 50% of WT, *p* represents the frequency of the most common allele, and *q* represents the frequency of the variant allele.

## Results

The primary objective of this study was to determine detailed functional properties of a large number of *KCNE1* variants co-expressed with KCNQ1 in a heterologous cell line with rigorous and standardized voltage-clamp protocols performed using automated patch clamp recording.^13, 14^ This approach provided a uniform experimental platform with a high degree of replication and lack of operator bias, which overcomes the heterogeneity of conventional voltage-clamp recording performed in diverse laboratory settings. Secondary goals were to determine if functional data can contribute to classifying the pathogenicity of *KCNE1* variants, and to test if findings from a deep mutation scan correlate with direct electrophysiological assessments. Finally, our knowledge of dysfunctional *KCNE1* variants enabled us to estimate the population carrier frequency of *KCNE1*-associated JLN2.

### Functional properties of KCNE1 variants

We determined the functional properties of all known disease-associated *KCNE1* variants reported prior to 2022 along with an approximately equal number of rare and ultra-rare *KCNE1* missense variants. Reported disease-associated variants were acquired from the professional version of the Human Gene Mutation Database (HGMD).^33^ Additional disease-associated variants were obtained from the literature and the ClinVar database entered prior to 2022. Population variants with minor allele frequencies less than 0.001 were ascertained from gnomAD (version 2.1.1). There was substantial overlap between these two groups of variants. In gnomAD, 22 variants were absent whereas 23 were heterozygous in a single individual (**Supplemental Table S2**). All other *KCNE1* variants we studied were heterozygous in 2 to 73 individuals in gnomAD.

Each variant was engineered in a recombinant human KCNE1 cDNA and co-expressed with WT human KCNQ1 in CHO-K1 cells to generate the slow delayed rectifying current (*I*_Ks_), which we pharmacologically isolated using a selective inhibitor (JNJ-303). In separate experiments, we mimicked the heterozygous state by expressing KCNE1 variants with WT KCNQ1 in a CHO-K1 cell line stably expressing WT KCNE1. Two prominent functional properties were measured including peak current density and voltage-dependence of activation quantified as the voltage-midpoint (V_½_) of conductance-voltage relationships. We also quantified activation and deactivation kinetics as time constants derived from exponential fits to the time course of these gating events. Functional results are presented separately for the three main KCNE1 structural domains (N-terminus, transmembrane, C-terminus). We first determined functional properties of variants expressed in the absence of WT KCNE1 (homozygous state), and then for those variants exhibiting abnormal function we separately measured properties in the presence of WT KCNE1 (mimicking the heterozygous state).

### Functional properties of KCNE1 variants in the homozygous state

KCNE1 variants analyzed in the homozygous state were distributed approximately equally among the three major protein domains: 36 in the N-terminus, 29 in the transmembrane domain and 30 in the C-terminus (**Supplemental Figure S1)**. Averaged whole-cell currents (normalized to WT channel peak amplitude) recorded from CHO-K1 cells co-expressing WT KCNQ1 with each of the 95 KCNE1 variants are presented in **Supplemental Figure S2**. We designated four groups based on current density: severe loss-of-function (peak current density ≤0.25 fold of WT *I*_Ks_), partial loss-of-function (peak current density 0.25-0.75 fold of WT *I*_Ks_), WT-like, and gain-of-function (peak current density >1.5 fold of WT *I*_Ks_). Complete quantitative data are presented in **Supplemental Dataset 1**.

Figure 1. illustrates peak current density relative to WT *I*_Ks_ for each of the 95 KCNE1 variants grouped by their location in the KCNE1 protein and presented as scatter plots consisting of all individual measurements, mean values and 95% confidence intervals. **Figure 2** summarizes mean peak current density data in volcano plots illustrating variants with statistically significant differences from WT *I*_Ks_ (plotted above the horizontal dashed line representing *P* = 0.02) either to the left or right of a vertical horizontal line indicating loss- or gain-of-function, respectively. Approximately half (47 of 95) of the KCNE1 variants tested in the homozygous state exhibited peak current densities significantly different from WT *I*_Ks_. Severe and partial loss-of-function were most common for variants in the transmembrane domain. Six variants (R32C, S34P, E83K, Q88K, Q96R, V99L) exhibited peak current density significantly greater than WT *I*_Ks_ and none were in the transmembrane domain.

**Figure 1.**
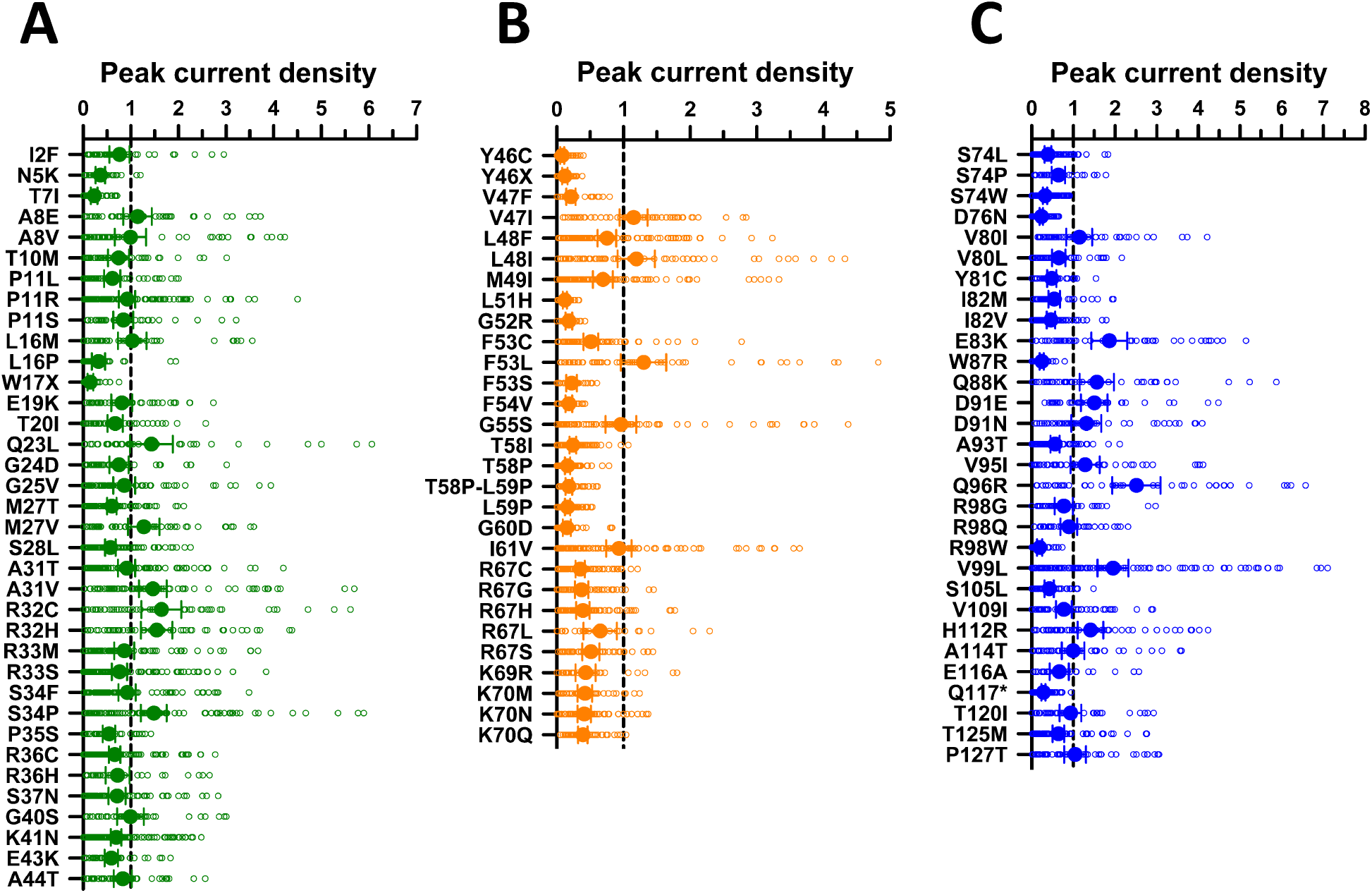
Peak current density for *KCNE1* variants in the homozygous state. JNJ-303 sensitive peak whole-cell currents measured at +60 mV from CHO-K1 cells co-expressing WT KCNQ1 with KCNE1 variants plotted as fold difference from WT channels recorded in parallel. (**A**) Variants in the N-terminus (green symbols). (**B**) Variants in the transmembrane domain (orange symbols). (**C**) Variants in the C-terminus (blue symbols). All individual data points are plotted as open symbols and mean values are shown as filled symbols with error bars representing the 95% CI. Values to the right or left of the vertical dashed line (normalized WT value) represent current density larger (gain-of-function) or smaller (loss-of-function) than WT, respectively. The number of recorded cells for each variant and individual *P*-values are presented in **Supplemental Dataset 1**.

**Figure 2.**
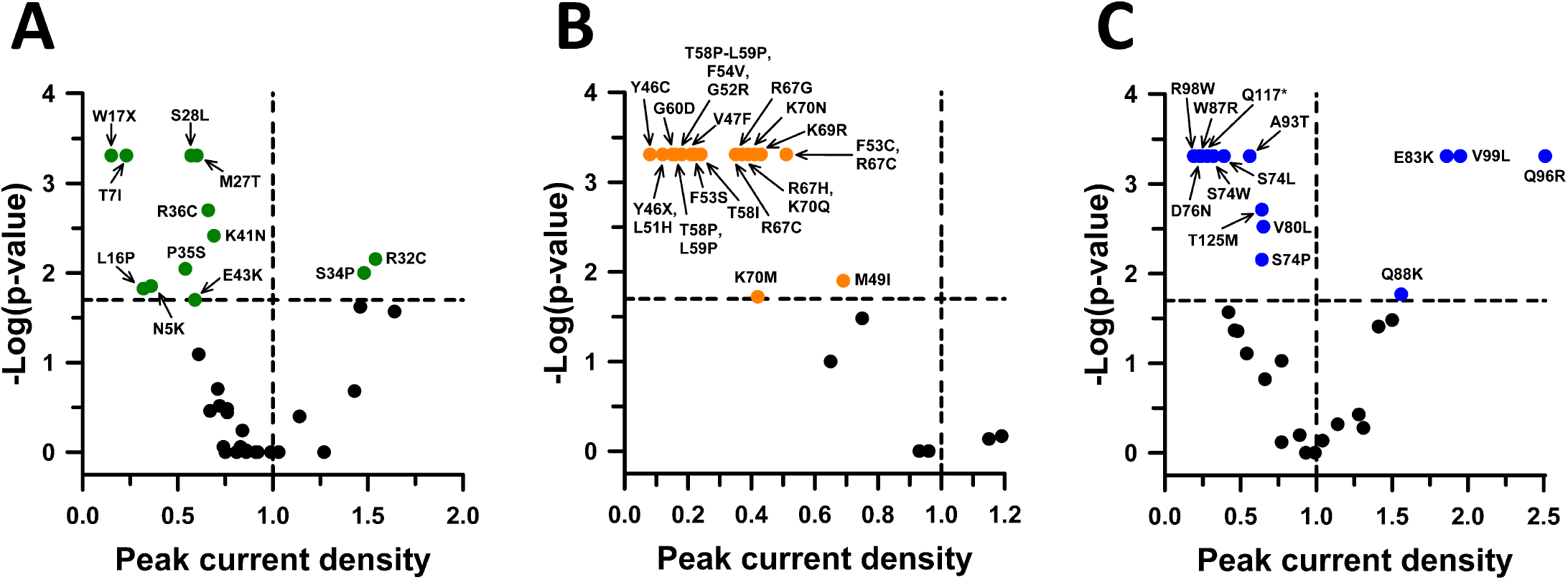
Volcano plots of current density for KCNE1 variants in the homozygous state. Volcano plots of mean JNJ-303 sensitive peak whole-cell currents from Figure 1. (**A**) Variants in the N-terminus (green symbols). (**B**) Variants in the transmembrane domain (orange symbols). (**C**) Variants in the C-terminus (blue symbols). Only variants with peak current density significantly different from WT (*P* < 0.02, horizontal dashed line) are labeled. Black symbols represent variants with no significant difference from WT. Symbols to the left of the vertical dashed line denote smaller (loss-of-function) and symbols to the right indicate larger (gain-of-function) current density.

Tail current amplitudes were sufficiently large to calculate voltage-dependence of activation parameters reliably for 78 variants. About one third (34%, 27/78) of variants predominantly in the transmembrane domain or C-terminus induced significantly hyperpolarizing (10 variants) or depolarizing (17 variants) shifts of activation V_1/2_ (**Figure 3; Supplemental Figure S3**). For these 78 variants, we also determined the voltage sensitivity of channel activation (*k*, the slope of the Boltzmann fit curve, **Supplemental Figure S4**). Interestingly, for all variants with significant shifts in activation V_½_, hyperpolarizing shifts were accompanied by larger slope values (10 variants), while depolarizing shifts had smaller slope values (17 variants). This association was independent of variant location.

**Figure 3.**
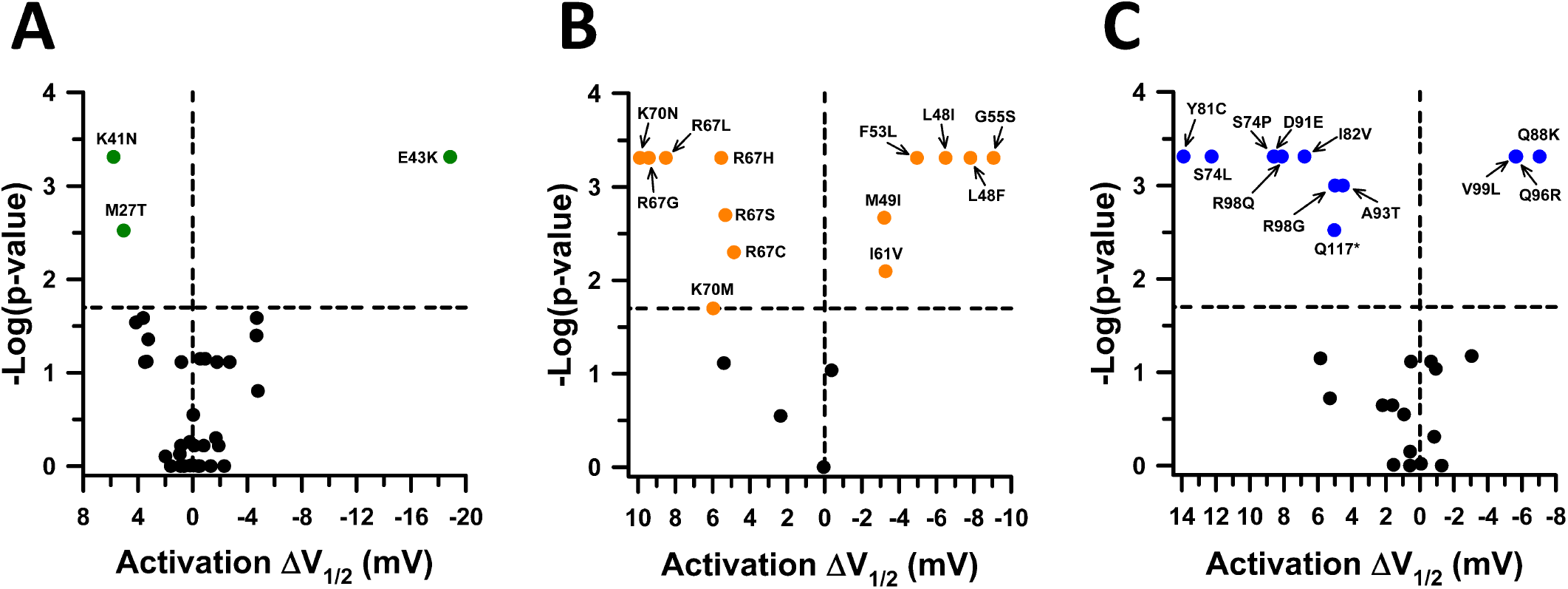
Activation voltage-dependence for KCNE1 variants in the homozygous state. Voltage-dependence of activation measured in KCNE1 variant-expressing cells plotted as the difference (ΔV½ in mV) from the averaged V½ for WT KCNE1 channels recorded in parallel and presented as volcano plots. (**A**) Variants in the N-terminus (green symbols). (**B**) Variants in the transmembrane domain (orange symbols). (**C**) Variants in the C-terminus (blue symbols). Only variants with V½ significantly different from WT (*P* < 0.02, horizontal dashed line) are labeled. Black symbols represent variants with no significant difference from WT. Symbols to the left of the vertical dashed line denote depolarized V½ values (loss-of-function), while symbols to the right indicate hyperpolarized V½ values (gain-of-function). Black symbols represent variants with no significant difference from WT. The number of recorded cells for each variant and individual *P*-values are presented in **Supplemental Dataset 1** and scatter plots of individual data are presented in **Supplemental Figure S3**.

We also determined the effect of homozygous KCNE1 variants on activation and deactivation kinetics. Sixteen variants caused significant changes in the rate of channel activation (**Figure 4; Supplemental Figure S5**). Five variants located in the N-terminus (T10M, R32C, P35S, A43K, A44T) and 7 in the transmembrane domain (M49I, V47F, L48F, G55S, T58I, I61V, R67G) caused faster activation relative to WT channel, while C-terminal variants caused either faster (S74W, Q88K, Q96R) or slower (D91E, R98Q, A114T) activation. Twenty-four KCNE1 variants evoked altered deactivation kinetics (**Figure 5; Supplemental Figure S6**). Variants in the transmembrane domain and C-terminus mostly induced faster deactivation relative to the WT channel. Four transmembrane variants (L48F, M49I, G55S, T58I) accelerated both activation and deactivation. Two variants in either the N-terminus (E43K) or C-terminus (D91E) affected both activation and deactivation in opposite directions, faster activation with slower deactivation for E43K, and slower activation with faster deactivation for D91E.

**Figure 4.**
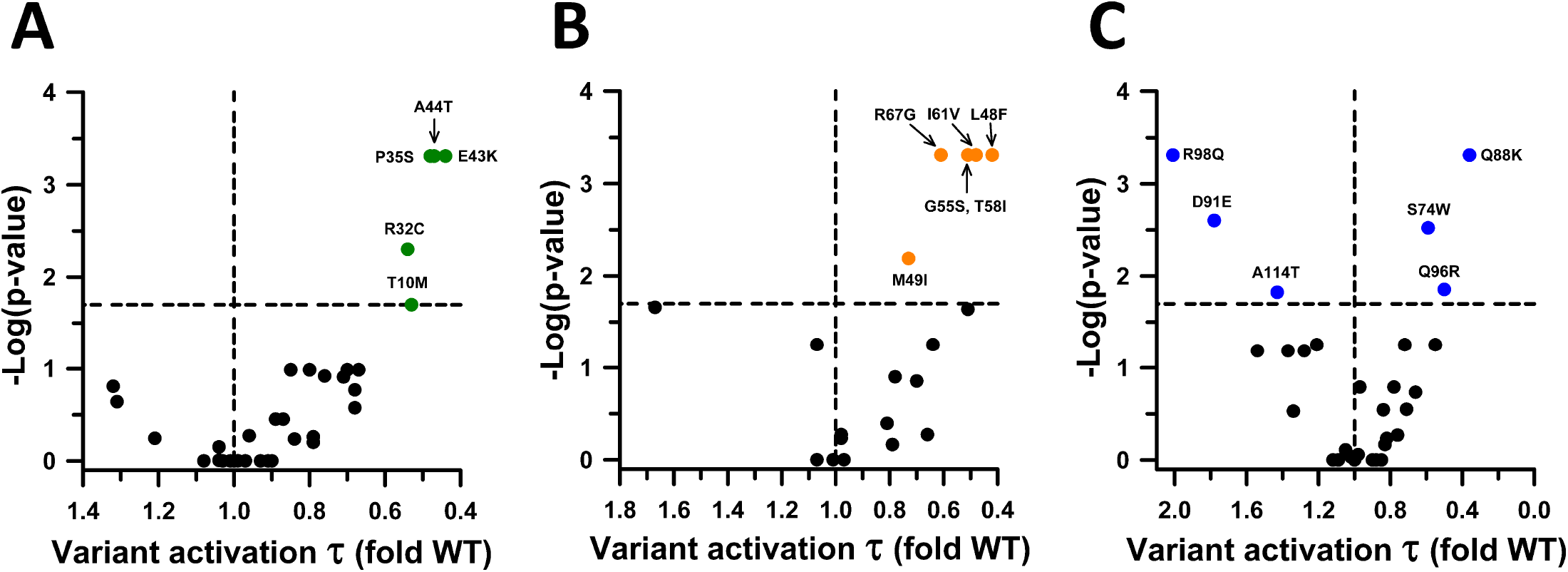
Activation kinetics for KCNE1 variants in the homozygous state. Activation time constants determined for each KCNE1 variant expressed as a ratio to the averaged value of WT channels recorded in parallel and presented as volcano plots. (**A**) Variants in the N-terminus (green symbols). (**B**) Variants in the transmembrane domain (orange symbols). (**C**) Variants in the C-terminus (blue symbols). Only variants with time constants significantly different from WT (*P* < 0.02, horizontal dashed line) are labeled. Black symbols represent variants with no significant difference from WT. Values to the right or left of the vertical dashed line (indicating no difference from WT) represent smaller (faster activation, gain-of-function) or larger (slower activation, loss-of-function) activation time constants, respectively. The number of recorded cells for each variant and individual *P*-values are presented in **Supplemental Dataset 1**. Scatter plots of individual data are presented in **Supplemental Figure S5**.

**Figure 5.**
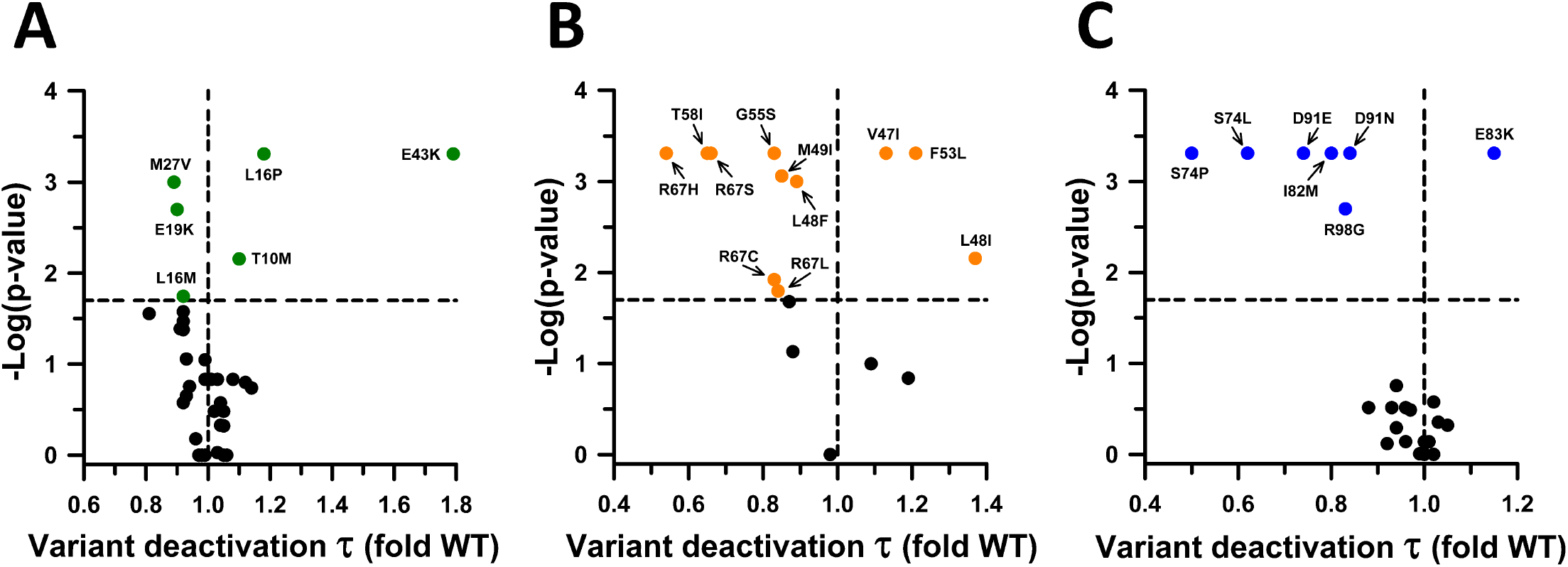
Deactivation kinetics for KCNE1 variants in the homozygous state. Deactivation time constants determined for each KCNE1 variant expressed as a ratio to the averaged value of WT channels recorded in parallel and presented as volcano plots. (**A**) Variants in the N-terminus (green symbols). (**B**) Variants in the transmembrane domain (orange symbols). (**C**) Variants in the C-terminus (blue symbols). Only variants with time constants significantly different from WT (*P* < 0.02, horizontal dashed line) are labeled. Black symbols represent variants with no significant difference from WT. Values to the right or left of the vertical dashed line (indicating no difference from WT) represent larger (slower deactivation, gain-of-function) or smaller (faster deactivation, loss-of-function) deactivation time constants, respectively. The number of recorded cells for each variant and individual *P*-values are presented in **Supplemental Dataset 1**. Scatter plots of individual data are presented in **Supplemental Figure S6**.

In summary, among the 95 KCNE1 variants tested in the homozygous state, 68 induced significant differences in the measured functional properties, while 27 variants did not significantly affect *I*_Ks_ function. Transmembrane domain variants exerted the largest number of significant effects on functional properties with 100% of these variants causing at least one significantly different functional property compared to WT. In contrast, variants in the N-terminus induced the least number of significant functional effects with approximately half (53%) exhibiting WT-like function. In the C-terminus, 22 variants (73%) affected at least one functional property.

### Functional properties of KCNE1 variants in the heterozygous state

To determine if any variants with significant dysfunction exerted dominant effects, we repeated voltage clamp experiments in the presence of WT KCNE1. The KCNE1 variants analyzed in the heterozygous state included 18 severe loss-of-function, 8 with partial loss-of-function, and 2 gain-of-function variants. Averaged normalized whole-cell currents recorded from CHO-E1 cells co-expressed with WT KCNQ1 and each of the 28 KCNE1 variants are presented in **Supplemental Figure S7**.

Figure 6. and **Supplemental Figure S8** illustrate the functional properties of KCNE1 variants expressed in the heterozygous state and complete quantitative data are presented in **Supplemental Dataset 2**. Only one variant (Q96R) (gain-of-function in the homozygous state) exhibited significantly different peak current density relative to the WT channel. None of the other 27 variants were significantly different from WT. In the heterozygous state, only 2 variants, both located in the C-terminus, caused significant differences in activation V_1/2_; D76N (severe loss-of-function in homozygous state) caused a depolarizing shift, while Q96R generated a hyperpolarizing shift, similar to when the variant was expressed in the homozygous state. Unlike in the homozygous state, no significant differences were observed for activation curve slope (**Supplemental Dataset 2**).

Analysis of the whole-cell current kinetics revealed that three transmembrane variants (G52R, F53S, T58P) in the heterozygous state induced significantly slower activation. All three variants exhibited severe loss-of-function in the homozygous state based on current density (**Figures 1 and 2**). Three variants in the transmembrane domain (G52R, F53S, L59P; all severe loss-of-function in the homozygous state) and three in the C-terminus (D76N, Y81C, W87R; severe or partial loss-of-function in the homozygous state) accelerated deactivation. Two transmembrane domain variants (F53S and T58P) slowed activation while also causing faster deactivation in the heterozygous state.

**Figure 6.**
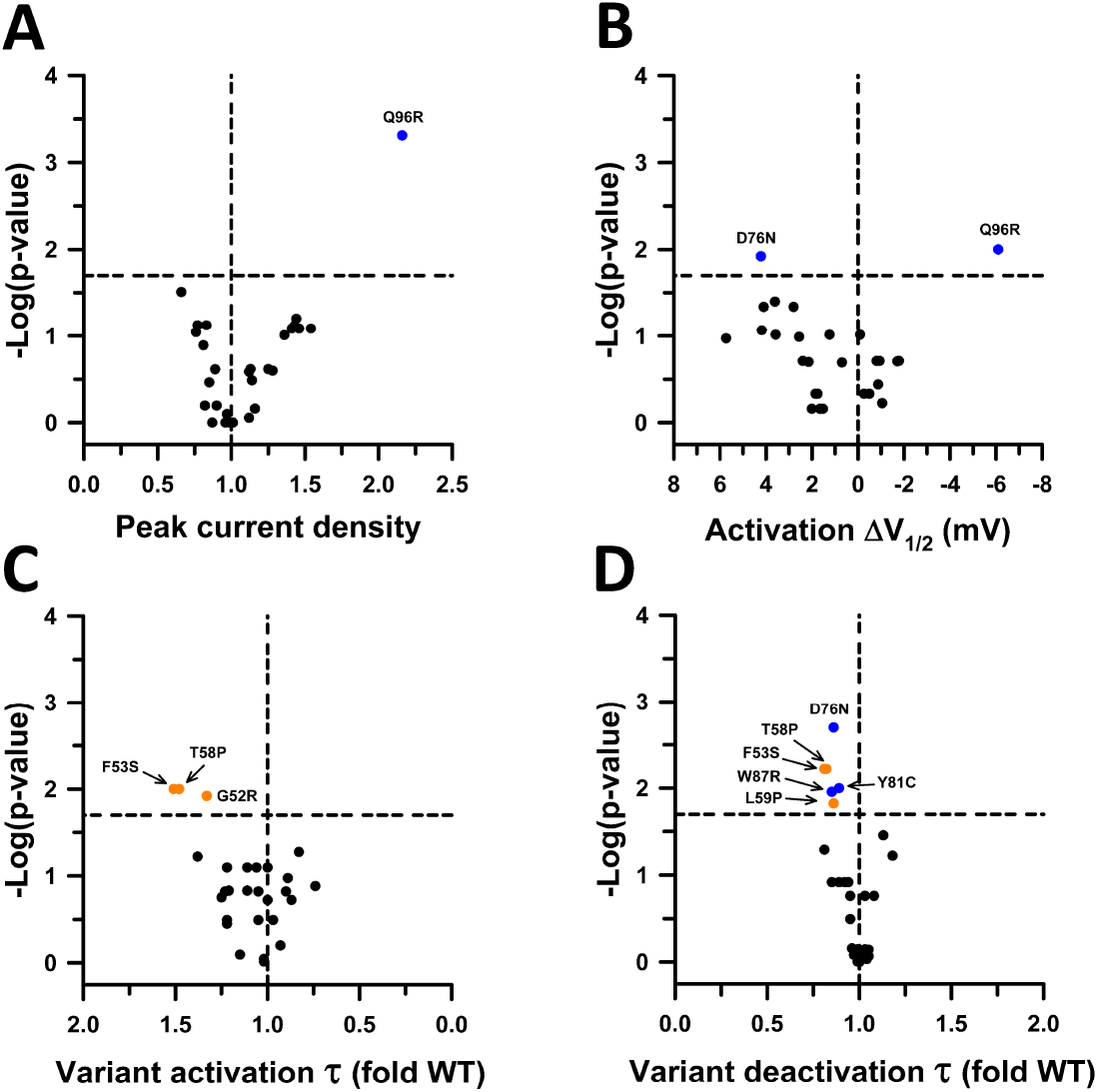
Functional properties of KCNE1 variants in the heterozygous state. (**A**) Volcano plots of mean JNJ-303 sensitive peak whole-cell current density measured at +60 mV from CHO-E1 cells co-expressing WT KCNQ1 with each KCNE1 variant. Data are displayed as fold divergence from WT channels recorded in parallel. Only variants with peak current density significantly different from WT (*P* < 0.02, horizontal dashed line) are labeled. Values to the right or left of the vertical dashed line (normalized WT value) represent current density larger or smaller than WT, respectively. (**B**) Volcano plots of mean difference (ΔV½ in mV) between the averaged V½ for variant and WT channels recorded in parallel. Only variants with peak current density significantly different from WT (P<0.02, horizontal dashed line) are labeled. Values to the right or left of the vertical dashed line indicate hyperpolarized or depolarized activation V½, respectively. (**C**) Volcano plots of mean activation time constants (τ) determined for each heterozygous KCNE1 variant plotted as the ratio to the averaged activation time constant for WT channels recorded in parallel. Values to the right or left of the vertical dashed line (no difference from WT) indicate faster or slower activation, respectively. (**D**) Volcano plots of mean deactivation time constants (τ) determined for each heterozygous KCNE1 variant plotted as the ratio to the averaged deactivation time constant for WT channels recorded in parallel. Values to the right or left of the vertical dashed line indicate slower or faster deactivation, respectively. Symbols colors are defined in previous figures. The number of recorded cells for each variant and individual *P*-values are presented in **Supplemental Dataset 2**. Scatter plots of individual data are presented in **Supplemental Figure S8**.

In summary of our functional study, all *KCNE1* variants with severe loss-of-function in the homozygous state exhibited peak current densities there were not significantly different from WT channels when tested in the heterozygous state, indicating that none of these variants exert overt dominant-negative effects. However, some of those variants induced slower activation and faster deactivation rates suggesting that co-expression of WT KCNE1 does not fully rescue some variants. In the C-terminus, the variant Q96R exerts a gain-of-function phenotype. This variant causes significant increase in peak current density and hyperpolarizing shift in V_½_ of activation of similar magnitude in both the homozygous and heterozygous state (**Supplemental Datasets 1 and 2)**.

### Impact of functional results on variant classification

To ascertain genotype-phenotype correlations and evaluate evidence of disease co-segregation, we performed an extensive review of published reports in which *KCNE1* was reported to be associated with long-QT syndrome or other clinical phenotypes (**Supplemental Table S3**). We found limited evidence supporting disease co-segregation with most reports describing *KCNE1* variants in isolated cases lacking a detailed family history. This is consistent with the findings of the ClinGen expert panel, which classified the association between *KCNE1* and autosomal dominant LQTS as ‘limited’.^7^

Application of ACMG criteria prior to consideration of functional data classified 91 as VUS, 2 as likely pathogenic (LP; A8V, G52R) and 2 as pathogenic (P; W17X, Y46X) (**Supplemental Table S2**). The inclusion of functional data resulted in revised ACMG classifications with 77 VUS, 15 LP, and 3 P (**Supplemental Table S2**). The 13 variants reclassified from VUS to LP exhibited severe loss-of-function in the homozygous state. Among 33 variants with partial loss-of-function, 32 remained classified as VUS and one was reclassified as LP (S74L). We initially applied functional data without considering gain-of-function as pathogenic or WT-like current density as benign criteria. In an expanded analysis we scored gain-of-function as PS3[M] and WT-like current density as BS3[P]. This resulted in additional reclassification of 5 VUS to LB (P11S, A31T, G40S, V80I, T120I) and 1 LP to VUS (A8V; **Supplemental Table S2**). There were no additional reclassifications from VUS to LP in this expanded analysis. Among the 22 variants absent in gnomAD, 21 were initially classified as VUS and six could be reclassified from VUS to LP (V47F, L51H, F53S, F54V, T58P-L59P, W87R) and one from LP to P (G52R) after consideration of expanded functional data.

### Estimation of JLN2 carrier frequency

*KCNE1* is associated with autosomal recessive LQTS with deafness (type 2 Jervell and Lange-Nielsen syndrome; JLN2). Co-inheritance of dysfunctional *KCNE1* variants as either homozygous or compound heterozygous genotypes would be compatible with this condition. Based on our functional study, we posited that variants exhibiting peak current densities less than 50% of WT *I*_Ks_ would be consistent with a pathogenic carrier allele. Therefore, we used functional data from our study combined with allele counts in gnomAD to estimate a population JLN2 carrier frequency (see Methods). Using this approach, we calculated the prevalence of JLN2 carriers in the population to be 0.000967 (1 in 1034).

### Correlation of voltage clamp recordings with KCNE1 deep mutational scan

Voltage clamp recording is widely considered a ‘gold’ standard experimental approach for determining the functional consequences of ion channel variants, but there are limitations in scale even with automation. A recent study using high throughput variant effect mapping of an epitope-tagged KCNE1 reported both cell surface expression and cell fitness in cells co-expressing WT KCNQ1 or a gain-of-function KCNQ1 variant (S140G) as proxies for function for a large number of single amino acid variants.^21^ We compared our findings to results from this deep mutational scan to determine the extent of correlation between the two approaches.

Figure 7 illustrates the correlation between peak current density measured in cells expressing WT KCNQ1 and different KCNE1 variants with the functional and trafficking scores deduced from the mutational scanning study. Both scores showed statistically significant linear correlations with measured current density although many variants were well outside 95% confidence intervals. The correlation was strongest with functional score (r^2^ = 0.304, *P* < 0.0001) but was also significantly correlated with trafficking scores (r^2^ = 0.092, *P* = 0.003) albeit less strong. The weaker correlation with trafficking scores suggests that current density can be influenced by factors other than cell surface expression (e.g., channel open probability, activation or deactivation kinetics). We also examined correlation with biophysical properties of the expressed channels (**Figure 8**). Activation voltage dependence quantified as the absolute difference between WT and variant activation V_½_ (ΔV_½_ in mV) was significantly correlated with functional scores (r^2^ = 0307, *P* < 0.0001), but there was no correlation with activation or deactivation kinetics.

## Discussion

*KCNE1* encodes a single transmembrane domain protein identified originally by expression cloning using *Xenopus laevis* oocytes.^34^ and initially thought to represent a novel minimal potassium channel sequence (dubbed minK).^35^ However, further investigations revealed that it was a modulator of KCNQ1 (originally named K_V_LQT1),^36^ the gene for type 1 LQTS implicating KCNE1 as an essential component of the *I*_Ks_ channel complex.^5,6^ The recognition of this physiologically important molecular partnership identified KCNE1 as a candidate gene for genetic arrhythmia susceptibility and lead to discovery of homozygous and compound heterozygous KCNE1 variants in a few families with JLN.^8^ The addition of KCNE1 to genetic testing panels for cardiac arrhythmia led to discoveries of heterozygous variants in a small subset of congenital LQTS cases designated as LQT5,^9^ which accounts for 1% or less of all genotype-positive LQTS cases.^37^

**Figure 7.**
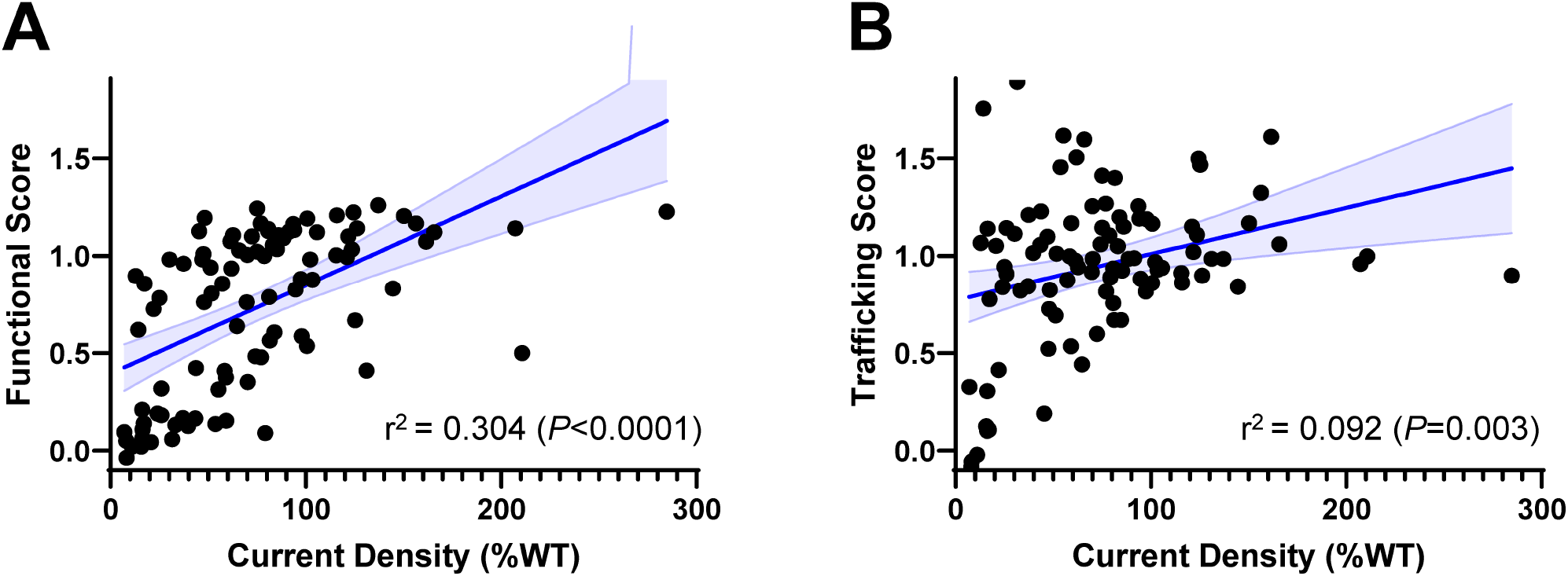
Correlation of peak current density with functional and trafficking scores from a KCNE1 deep mutation scan. (**A**) Plot of peak current density measured in cells co-expressing WT KCNQ1 with KCNE1 variants (homozygous state) compared with the Functional Score reported for each variant determined by a cell fitness assay.^21^ (**B**) Plot of peak current density measured in cells co-expressing WT KCNQ1 with KCNE1 variants (homozygous state) compared with the reported Trafficking Score for each variant.^21^ In both plots, solid blue lines represent a linear regression fit to the data with 95% CI shown as light blue shadow.

**Figure 8.**
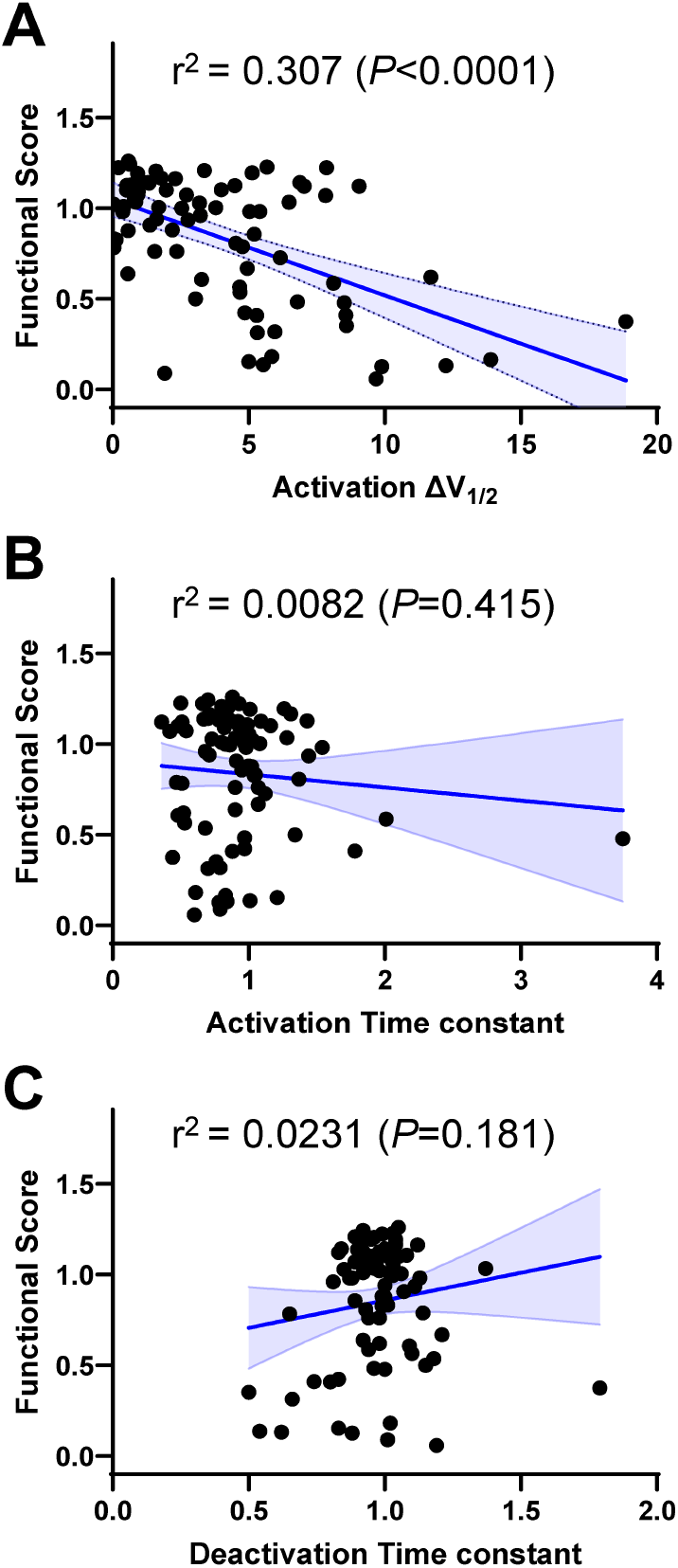
Correlation of biophysical properties with functional scores from a KCNE1 deep mutation scan. (**A**) Plot of differences in activation V½ (ΔV½ in mV) measured in cells co-expressing WT KCNQ1 with KCNE1 variants (homozygous state) compared with the Functional Score reported for each variant determined by a cell fitness assay.^21^ (**B**) Plot of differences in activation time constants measured in cells co-expressing WT KCNQ1 with KCNE1 variants (homozygous state) compared with the reported Functional Score. (**C**) Plot of differences in deactivation time constants measured in cells co-expressing WT KCNQ1 with KCNE1 variants (homozygous state) compared with the reported Functional Score.

While genetic evidence for association between *KCNE1* and JLN, a recessive disease, appears strong, the evidence supporting the association of *KCNE1* with autosomal dominant LQTS only reached the strength of limited when reviewed by a ClinGen expert panel.^7^ Additionally, an international multicenter study concluded that heterozygous loss-of-function *KCNE1* variants were low penetrance contributors to arrhythmia susceptibility with most carriers exhibiting mild to no clinical phenotypes.^10^ Moreover, the emergence of population exome and genome sequencing data revealed the presence of many rare *KCNE1* variants that had been implicated in LQT5, further supporting the assertion of low penetrance. However, a stated limitation of this conclusion was the absence of more complete functional assessments of putative disease-causing variants.^10^ In this study, we sought to evaluate the functional consequences of many human *KCNE1* variants associated with either cardiac arrhythmia or discovered in a population sample (gnomAD). We used a scalable approach based on automated patch clamp recording that was validated previously for study of *KCNQ1* variants.^13,18^ We found extensive overlap in functional behavior between disease-associated and population KCNE1 missense variants. Furthermore, none of the studied variants exhibited dominant-negative effects that would be consistent with highly penetrant disease-causing variants in a heterozygous state. We did observe that several *KCNE1* variants exhibited substantial loss-of-function effects in the homozygous state that were able to drive reclassification of some VUS to pathogenic categories. We also used these data to infer a population carrier frequency of JLN2. Overall, our study contributes to clarifying the genetic contributions of *KCNE1* to congenital arrhythmia susceptibility.

Our findings coupled with a careful review of the literature are consistent with the notion that *KCNE1* contributes to clinically significant arrhythmia susceptibility mainly as a recessive gene. Our estimate of the JLN2 carrier frequency (1 in 1034) indicates that *KCNE1* is not among the most frequent recessive genes in the population with a carrier frequency similar to *MYH7*, a rare inherited cause of cardiomyopathy.^32^ Previous reports of severe arrhythmia phenotypes associated with heterozygous KCNE1 variants may be explained by a reporting bias that highlights the most severe cases recognized before availability of population variant frequencies. An alternative viewpoint as suggested by Roberts et al.,^10^ is that *KCNE1* variants act as modifiers with low penetrance individually, requiring other concurrent genetic or acquired factors to manifest as LQTS. The idea of concurrent factors is supported by evidence that a common *KCNE1* variant (D85N) is a risk factor for drug-induced LQTS.^38^ Other unrecognized factors including undetected pathogenic variants in other known or unknown arrhythmia susceptibility genes may exist in LQT5 cases, and arrhythmia susceptibility in these cases may be oligogenic rather than monogenic. Inclusion of *KCNE1* as a potential monogenic cause of autosomal dominant LQTS on clinical genetic testing panels may warrant reconsideration as suggested by the ClinGen panel.^7^

We also examined the correlation between our direct electrophysiological assessments of 95 *KCNE1* variants and data from a deep mutational scan of this gene that used indirect proxies of function.^21^ As illustrated in **Figure 7**, there was significant correlation between direct measurement of peak current density and a functional score derived from a cell viability assay. This result is encouraging and supports the validity of the mutational scanning data for general functional effects. We also highlight a significant correlation between the functional score and activation voltage-dependence (**Figure 8A**) that further underscores the potential value of the approach. By contrast, the cell viability assay did not correlate with more nuanced functional consequences of KCNE1 variants (activation and deactivation kinetics; **Figure 8B**,**C**) that may contribute to arrhythmia susceptibility. Importantly, the reported deep mutational scan did not assess KCNE1 variants in the heterozygous state, but refinement of the assay (e.g., constitutive expression of WT KCNE1 in the cellular platform) might be possible to address this point.

## Study Limitations

We did not consider other potential interacting partners of KCNE1 such as the human ether-á-go-go related gene (hERG), which might be relevant.^39^ We did not consider performing such studies given the uncertain physiological relevance of hERG interactions with KCNE1 that were demonstrated only in overexpression studies *in vitro*. We also did not use the most relevant cell type (e.g., cardiac myocytes) to conduct our study because of the challenges inherent in culturing, transfecting and recording from such cells that would have severely limited throughput. Finally, in our heterozygous assay, we cannot control stoichiometry of WT and variant KCNE1 in transiently transfected cells. However, using automated patch clamp, we were able to record from a large number of replicates that should sample the range of subunit ratios that arise from random assortment analogous to expression in native cells.

## Conclusions

In this study, we determined the functional consequences of 95 *KCNE1* variants that enabled us to demonstrate absence of a dominant-negative molecular mechanism, infer a population carrier frequency of recessive JLN2, and benchmark findings from an orthogonal deep mutational scan. This work helps clarify the pathogenicity of *KCNE1*, contributes to emerging doubt regarding this gene as a monogenic cause of dominant cardiac arrhythmia, and validates deep mutational scanning for assessing some functional properties of variants in this gene.

## Acknowledgements

This work was supported by NIH grant HL122010.

---

**Supplemental Fig. S1.**
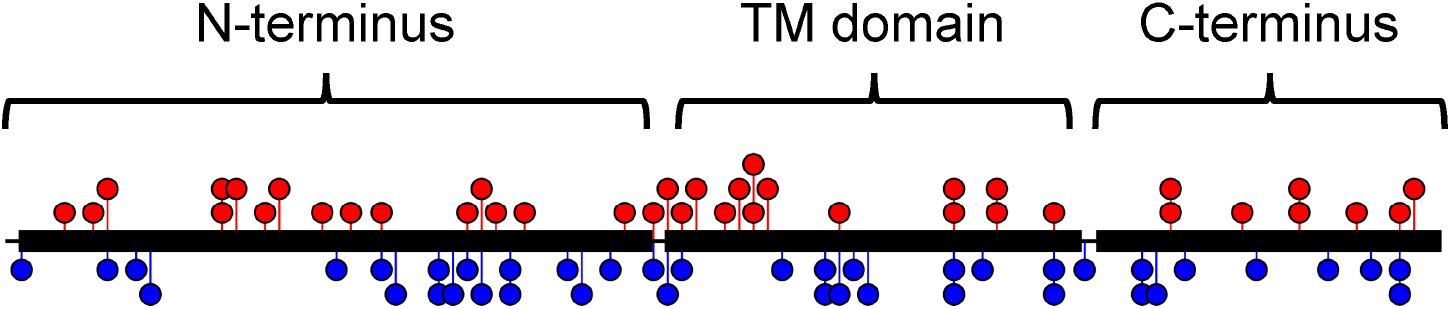
Distribution of variants in the KCNE1 protein. Approximate location of KCNE1 variants within the N-terminus, transmembrane (TM) domain, and C-terminus. Red ‘lollipops’ projecting above the horizontal line represent variants found 0 or 1 times in gnomAD. Blue ‘lollipops’ projecting below the horizontal line represent variants found 2 or more times in gnomAD. The height of each lollipop corresponds to the number of variants at that location. Details about specific variants is presented in Supplemental Table S2.

**Supplemental Fig. S2.**
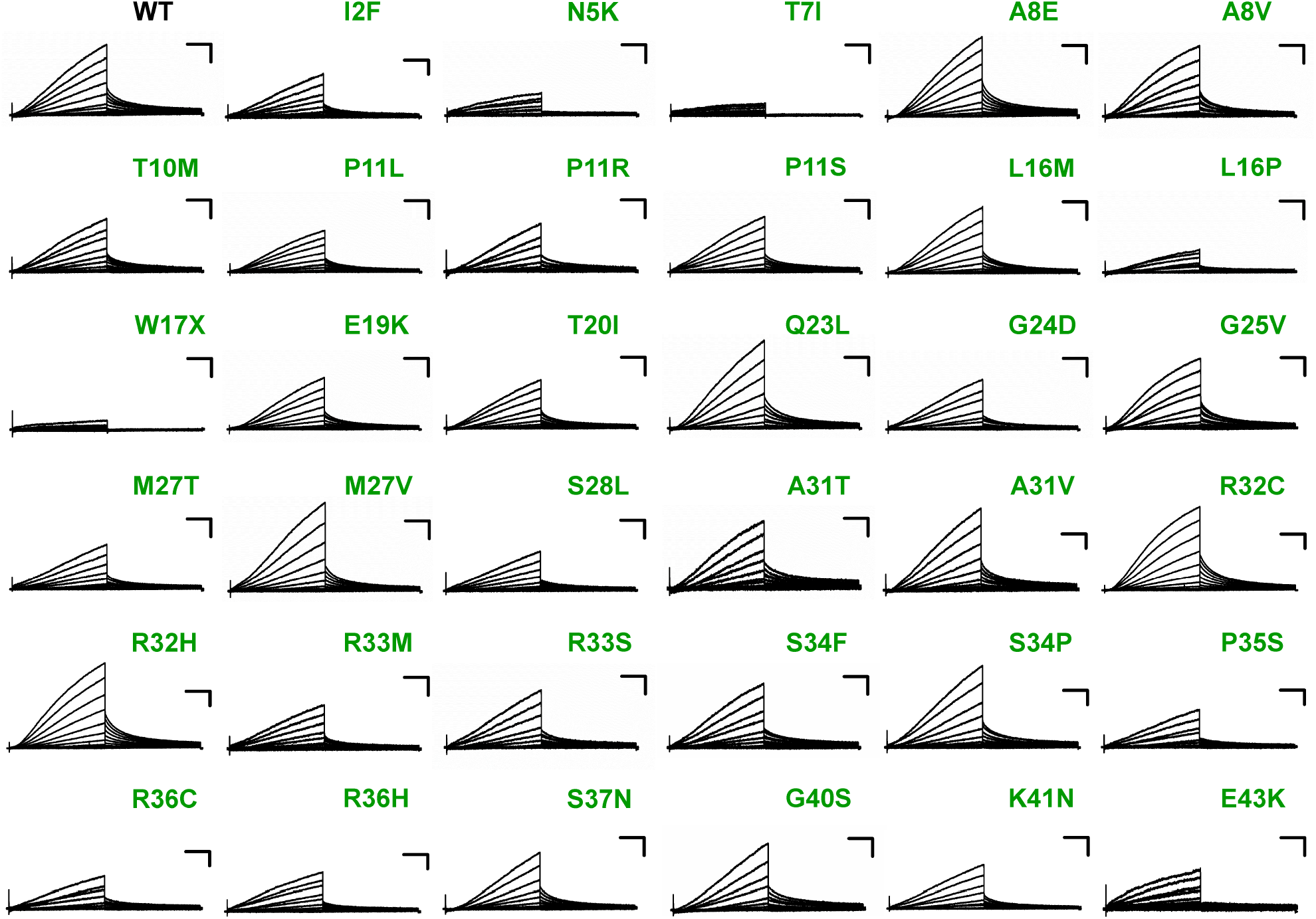
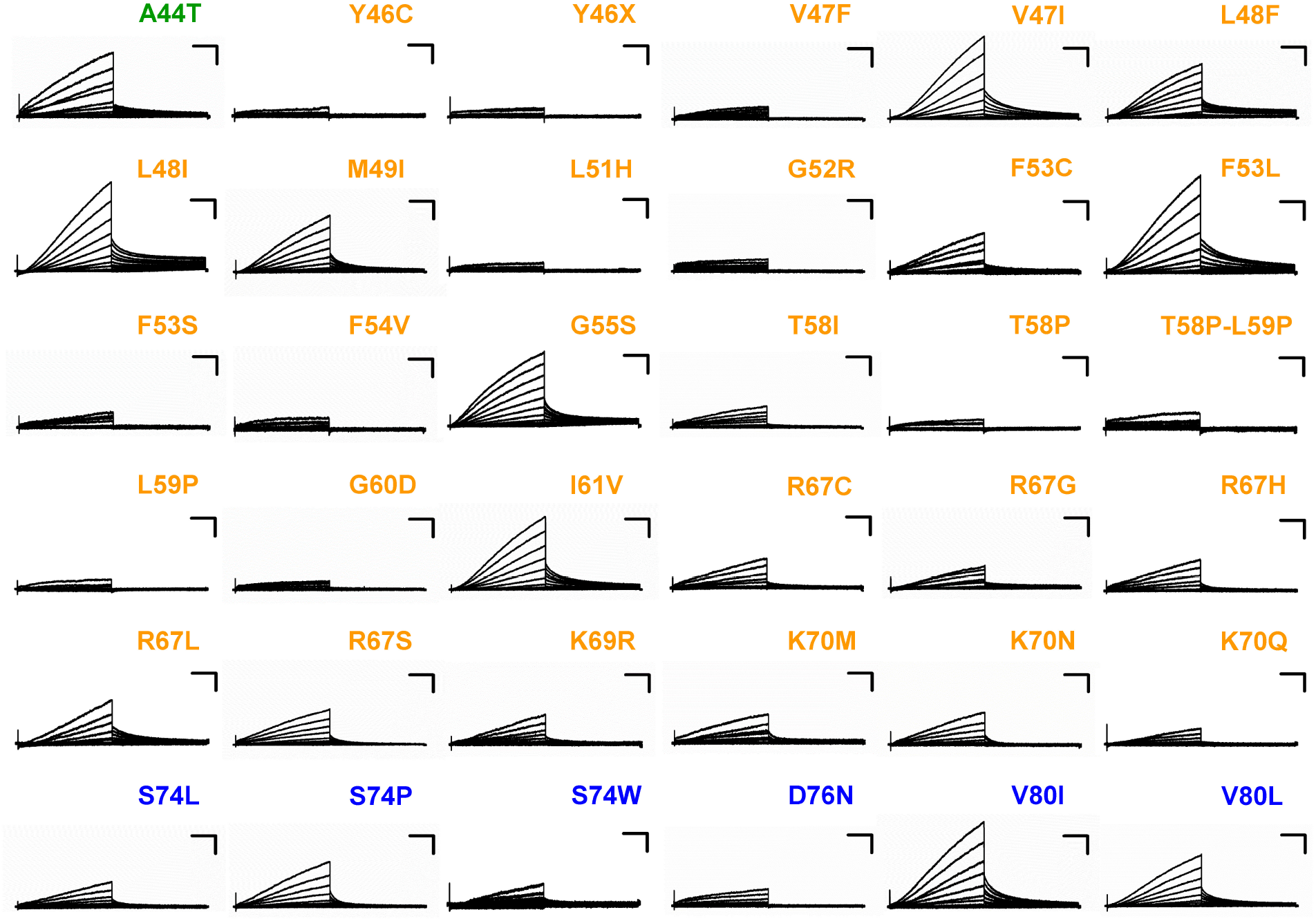
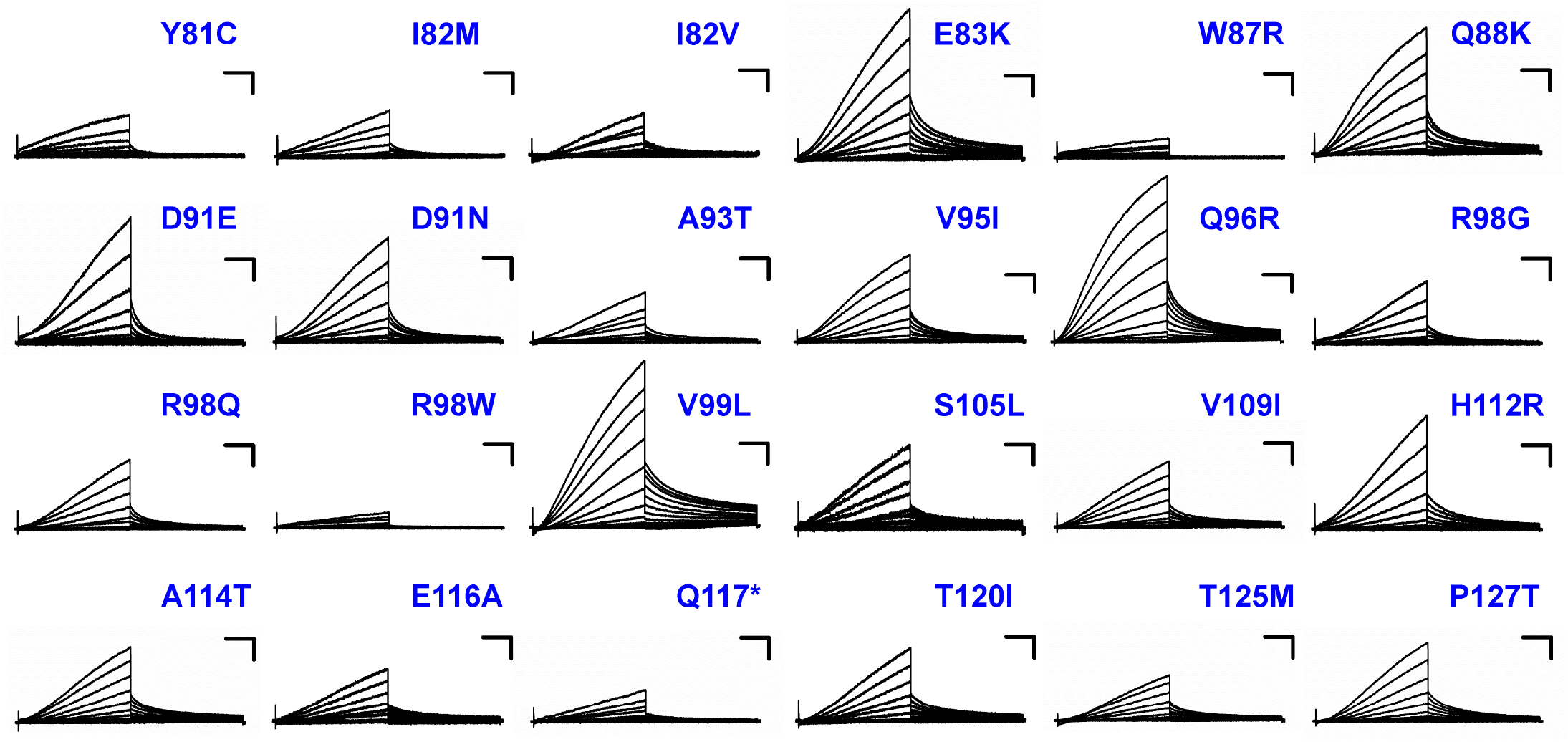
Averaged whole cell current traces from cells expressing KCNE1 variants in the homozygous state. Each trace represents JNJ-303 sensitive currents from an average of 21-126 cells co-expressing KCNQ1 with the indicated KCNE1 variant. Currents were normalized to peak current in cells expressing WT KCNE1 and WT KCNQ1 recorded in parallel. Scale bars represent 25% of WT (vertical) and 500 ms (horizontal). Location of variants is indicated by the colored labels (green = N-terminus; orange = TM domain; blue = C-terminus).

**Supplemental Fig. S3.**
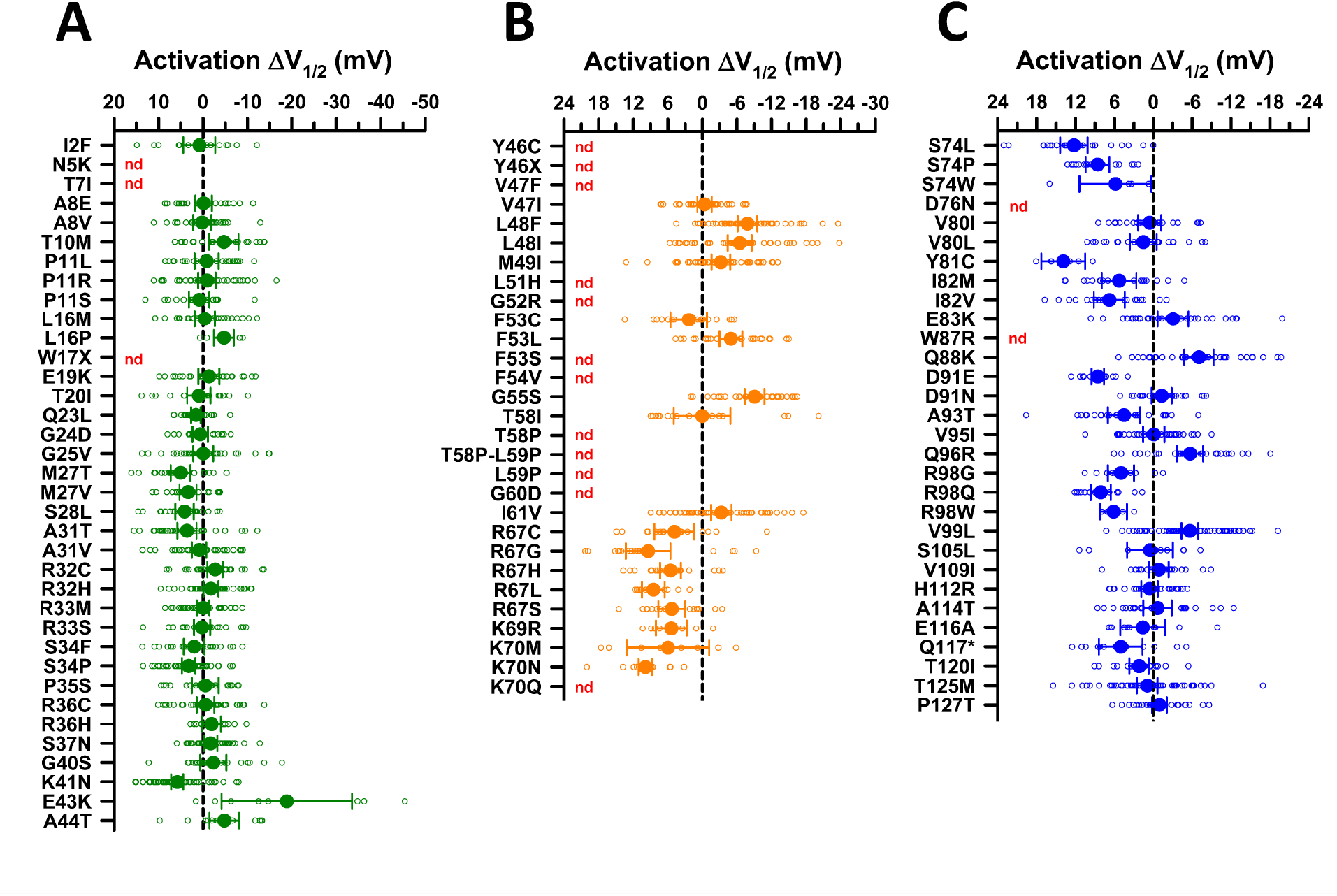
Activation voltage-dependence of KCNE1 variants (homozygous state). Averaged voltage-dependence of activation measured in KCNE1 variant-expressing cells plotted as the difference (ΔV½ in mV) from the averaged V½ for WT KCNE1 channels recorded in parallel. (**A**) Variants in the N-terminus (green symbols). (**B**) Variants in the transmembrane domain (orange symbols). (**C**) Variants in the C-terminus (blue symbols). All individual data points are plotted as open symbols and mean values are shown as filled symbols with error bars representing the 95% CI. Values to the right or left of the vertical dashed line (indicating no difference from WT) represent hyperpolarized (gain-of-function) or depolarized (loss-of-function) activation V½, respectively. For some variants, activation V½ could not be determined (nd). The number of recorded cells for each variant and individual *P*-values are presented in **Dataset 1**.

**Supplemental Fig. S4.**
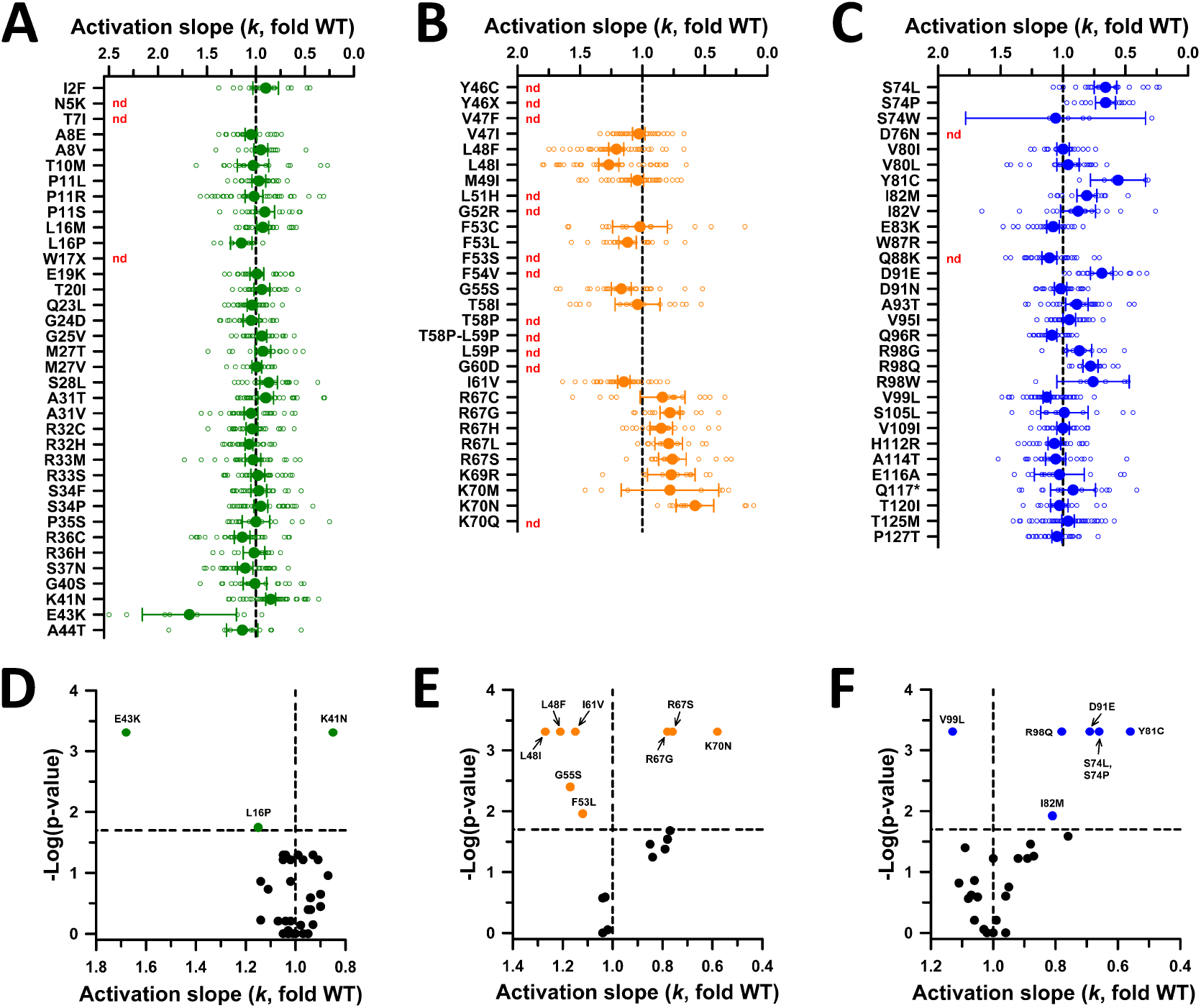
Activation slope factors of KCNE1 variants (homozygous state). Average slope for voltage-dependence of activation curves (*k*) determined from fitting the data for each KCNE1 variant-expressing cell to a Boltzmann function and plotted as ratio from the averaged *k* for WT KCNE1 channels recorded in parallel. (**A**) Variants in the N-terminus (green symbols). (**B**) Variants in the transmembrane domain (orange symbols). (**C**) Variants in the C-terminus (blue symbols). All individual data points are plotted in panels **A**-**C** as open symbols and mean values are shown as filled symbols with error bars representing the 95% CI. Values to the right or left of the vertical dashed line (indicating no difference from WT) represent steeper (i.e., smaller voltage change for increase in function) or shallower (i.e., larger voltage change for increase in function) activation curve slope, respectively. (**D-F**) Volcano plots of mean values for KCNE1 variants with *k* values significantly different from WT (*P* < 0.02, horizontal dashed line). Symbols to the left of the vertical dashed line denote larger *k* values (loss-of-function), while symbols to the right indicate smaller *k* values (gain-of-function). Symbol color denotes location with KCNE1 domains as described for panels **A**-**C**. Unlabeled black symbols represent variants with no significant difference from WT. The number of recorded cells for each variant and individual *P*-values are presented in **Dataset 1**.

**Supplemental Fig. S5.**
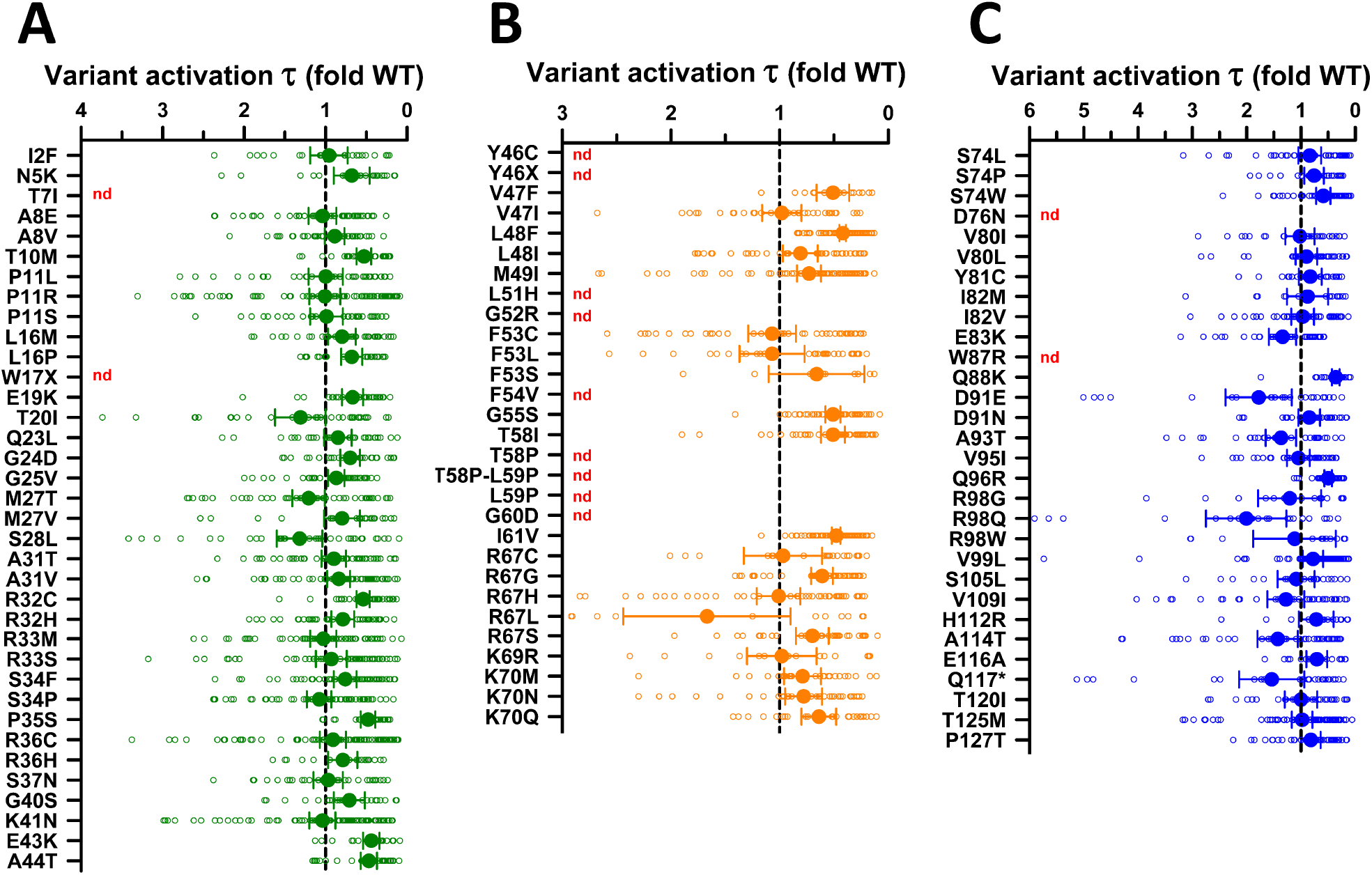
Activation time constants of KCNE1 variants (homozygous state). Activation time constants determined for each KCNE1 variant expressed as a ratio to the averaged activation time constant of WT channels recorded in parallel. (**A**) Variants in the N-terminus (green symbols). (**B**) Variants in the transmembrane domain (orange symbols). (**C**) Variants in the C-terminus (blue symbols). All individual data points are plotted as open symbols and mean values are shown as filled symbols with error bars representing the 95% CI. Values to the right or left of the vertical dashed line (indicating no difference from WT) represent smaller (faster activation, gain-of-function) or larger (slower activation, loss-of-function) activation time constants, respectively. For some variants, activation time constants could not be determined (nd). The number of recorded cells for each variant and individual *P*-values are presented in **Dataset 1**.

**Supplemental Fig. S6.**
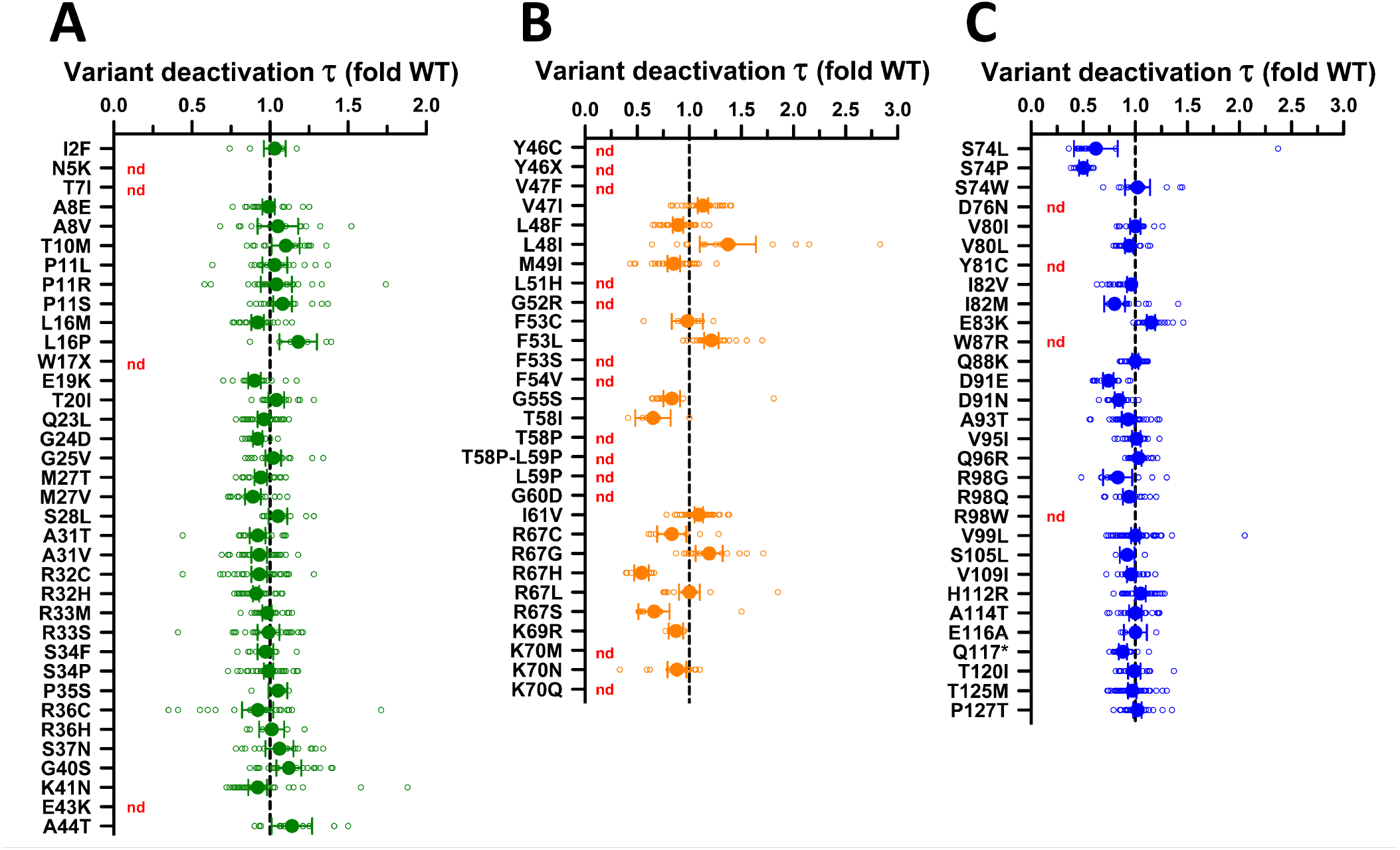
Deactivation time constants of KCNE1 variants (homozygous state). Deactivation time constants determined for each KCNE1 variant expressed as a ratio to the averaged deactivation time constant of WT channels recorded in parallel. (**A**) Variants in the N-terminus (green symbols). (**B**) Variants in the transmembrane domain (orange symbols). (**C**) Variants in the C-terminus (blue symbols). All individual data points are plotted as open symbols and mean values are shown as filled symbols with error bars representing the 95% CI. Values to the right or left of the vertical dashed line (indicating no difference from WT) represent larger (slower deactivation, gain-of-function) or smaller (faster deactivation, loss-of-function) deactivation time constants, respectively. For some variants, deactivation time constants could not be determined (nd). The number of recorded cells for each variant and individual *P*-values are presented in **Dataset 1**.

**Supplemental Fig. S7.**
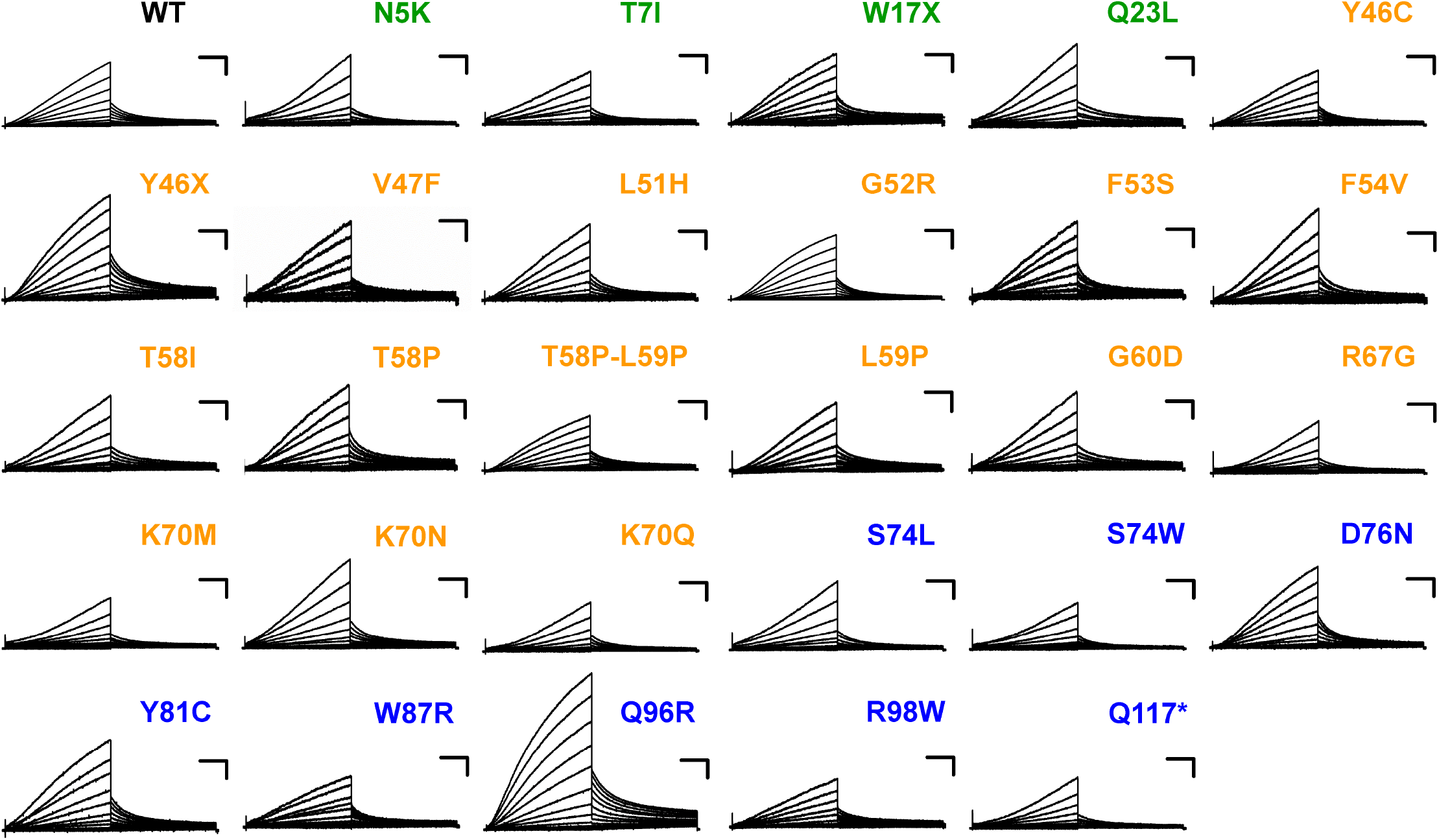
Averaged whole cell current traces from cells expressing KCNE1 variants in the heterozygous state. Each trace represents JNJ-303 sensitive currents from an average of 38-131 CHO-E1 cells coexpressing KCNQ1 with the indicated KCNE1 variant. Currents were normalized to peak current in CHO-E1 cells co-expressing WT KCNE1 and WT KCNQ1 recorded in parallel. Scale bars represent 25% of WT (vertical) and 500 ms (horizontal). Location of variants is indicated by the colored labels (green = N-terminus; orange = TM domain; blue = C-terminus).

**Supplemental Fig. S8.**
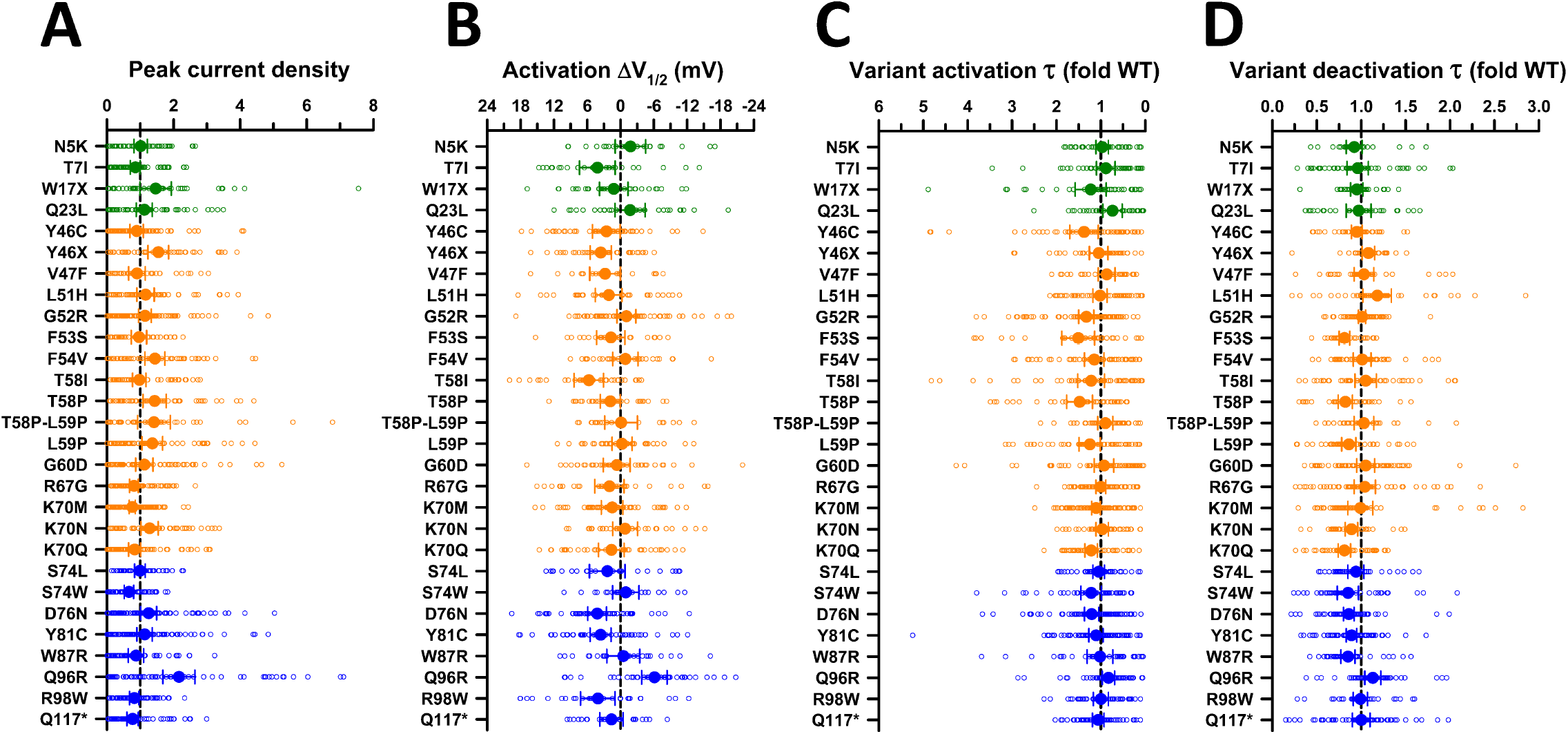
Functional properties of KCNE1 variants in the heterozygous state. (**A**) Average JNJ-303-sensitive peak whole-cell current density measured at +60 mV from CHO-E1 cells co-expressing WT KCNQ1 with each KCNE1 variant. Data are displayed as fold divergence from WT channels recorded in parallel. Values to the right or left of the vertical dashed line (normalized WT value) represent current density larger or smaller than WT, respectively. (**B**) Activation voltage-dependence for each heterozygous KCNE1 variant plotted as the difference (ΔV½ in mV) between the averaged V½ for WT channels recorded in parallel. Values to the right or left of the vertical dashed line indicate hyperpolarized or depolarized activation V½, respectively. (**C**) Activation time constants (τ) determined for each heterozygous KCNE1 variant plotted as the ratio to the averaged activation time constant for WT channels recorded in parallel. Values to the right or left of the vertical dashed line (no difference from WT) indicate faster or slower activation, respectively. (**D**) Deactivation time constants (τ) determined for each heterozygous KCNE1 variant plotted as the ratio to the averaged deactivation time constant for WT channels recorded in parallel. Values to the right or left of the vertical dashed line indicate slower or faster deactivation, respectively. Location of variants is indicated by the colored symbols (green = N-terminus; orange = TM domain; blue = C-terminus). The number of recorded cells for each variant and individual *P*-values are presented in **Dataset 2**.

**Supplemental Table S1.**
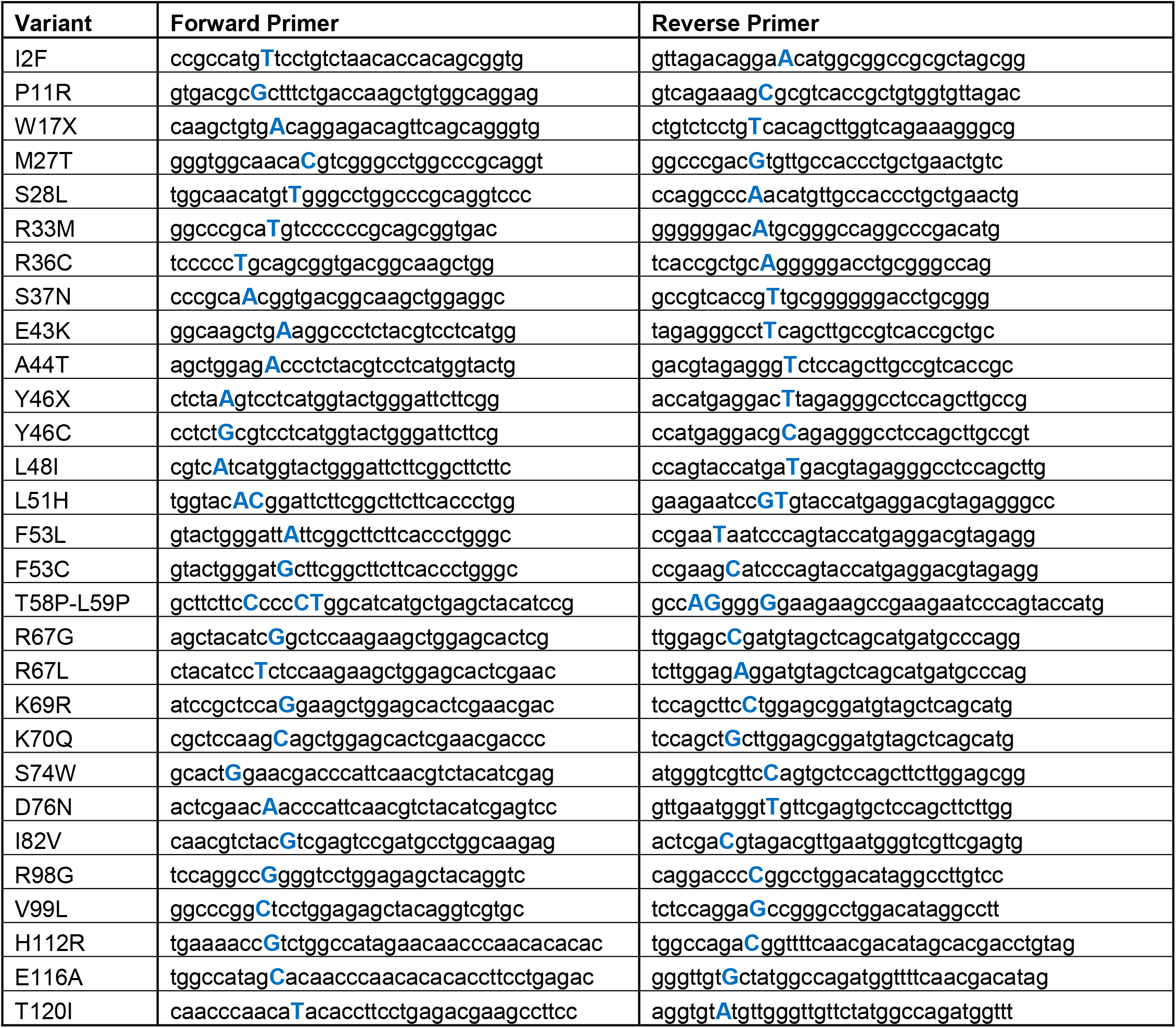
Primer sequences for site-directed mutagenesis of KCNE1 (29 variants).

**Supplemental Table S2.**
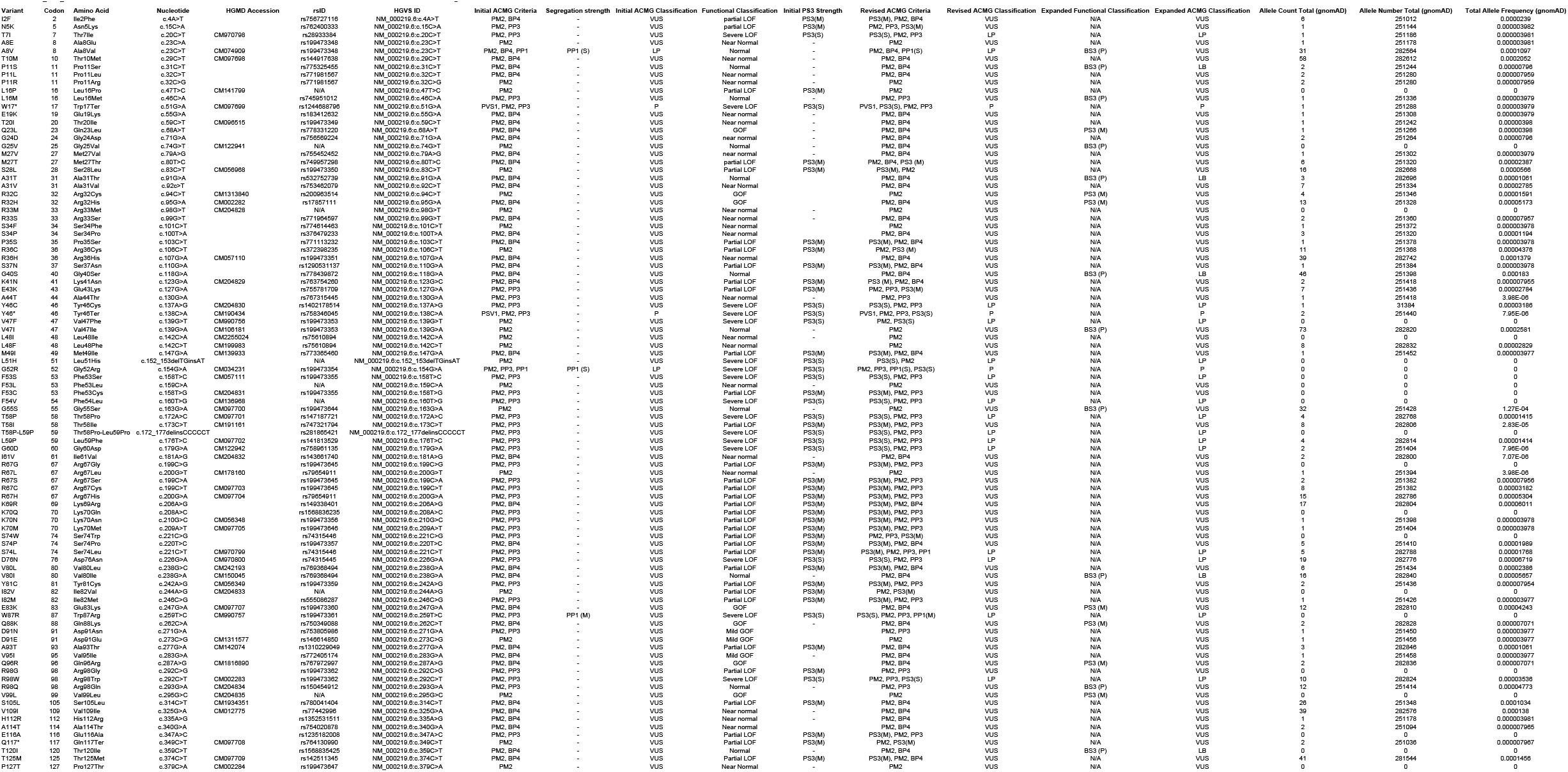
Summary of KCNE1 variant information and classifications.

**Supplemental Table S3.** Literature review of KCNE1 genotype-phenotype correlations. References to original research concerning each variant investigated in our study. Literature was obtained from *KCNE1* variant entries in ClinVar including their corresponding LitVar findings, Bibliome.ai browser, and Mastermind Genomic Search Engine by Genomenon. Additional literature was obtained via Google Search using keywords in the following search string: (KCNE1 OR ISK OR JLNS OR JLNS2 OR LQT2/5 OR LQT5 OR MinK OR “potassium voltage-gated channel subfamily E regulatory subunit 1”) AND ([single letter code amino acid variant] OR [three letter code amino acid variant] OR p.[three letter code amino acid variant] OR “[complementary DNA change]” OR “[dbSNP identity]”) with aliases for *KCNE1* in the first set of parentheses and variant-specific identifiers in the second set of parentheses, with bracketed terms adjusted per variant. Of the variants investigated in this study that do not possess a dbSNP ID (G25V, L51H, F54V, I82V, and V99L), search strings were entered as above without the dbSNP ID. An example of such a search string would be the following for *KCNE1* D76N: (KCNE1 OR ISK OR JLNS OR JLNS2 OR LQT2/5 OR LQT5 OR MinK OR “potassium voltage-gated channel subfamily E regulatory subunit 1”) AND (D76N OR Asp76Asn OR p.Asp76Asn OR “c.226G>A” OR “rs74315445”) Sources were included provided they list specific germline nucleotide changes and/or amino acid changes. Sources were excluded if they included identifiers for multiple missense variants without specifying nucleotide and/or protein changes (eg, a source includes rs199473348 without distinguishing between A8V or A8E). Sources that investigated cDNA changes not used in constructs in this study were excluded (eg, both c.273C>A and c.273C>G code for D91E but only the sources investigating c.273C>A were included). References with somatic mutations were excluded.

| Codon | Nucleotide Change | Protein Change | dbSNP ID | Studies with Genomic / Phylogenetic / Clinical / Pedigree Data | Studies with Functional Assay Data |
| --- | --- | --- | --- | --- | --- |
| 2 | c.4A>T | I2F | rs756727116 | <sup>1</sup> | This study |
| 5 | c.15C>A | N5K | rs762400333 | None found | This study |
| 7 | c.20C>T | T7I | rs28933384 | <sup>2-14</sup> | <sup>15-18</sup>, This study |
| 8 | c.23C>A | A8E | rs199473348 | None found | This study |
| 8 | c.23C>T | A8V | rs199473348 | <sup>1,7,12,14,19-32</sup> | <sup>32,33</sup>, This study |
| 10 | c.29C>T | T10M | rs144917638 | <sup>1,7,12,14,19,20,22,28,29,34-39</sup> | This study |
| 11 | c.31C>T | P11S | rs775325455 | None found | This study |
| 11 | c.32C>G | P11R | rs771981567 | None found | This study |
| 11 | c.32C>T | P11L | rs771981567 | None found | This study |
| 16 | c.46C>A | L16M | rs745951012 | None found | This study |
| 16 | c.47T>C | L16P | rs1981057774 | <sup>40</sup> | This study |
| 17 | c.51G>A | W17X | rs1244688796 | <sup>7,29,41</sup> | This study |
| 19 | c.55G>A | E19K | rs183412632 | <sup>42</sup> | <sup>43</sup>, This study |
| 20 | c.59C>T | T20I | rs199473349 | <sup>12,44-47</sup> | This study |
| 23 | c.68A>T | Q23L | rs778331220 | None found | This study |
| 24 | c.71G>A | G24D | rs756569224 | None found | This study |
| 25 | c.74G>T | G25V | Not found | 20,48-50 | <sup>50</sup>, This study |
| 27 | c.79A>G | M27V | rs755452452 | None found | This study |
| 27 | c.80T>C | M27T | rs749957298 | None found | This study |
| 28 | c.83C>T | S28L | rs199473350 | 1,7,12,29,38,47,51-61 | <sup>51</sup>, This study |
| 31 | c.91G>A | A31T | rs753462079 | <sup>1</sup> | This study |
| 31 | c.92C>T | A31V | rs532752739 | None found | This study |
| 32 | c.94C>T | R32C | rs200963514 | 1,37,42,59,62 | <sup>63,64</sup>, This study |
| 32 | c.95G>A | R32H | rs17857111 | 1,3,10,12,13,29,38,39,59,65-74 | <sup>15,74</sup>, This study |
| 33 | c.98G>T | R33M | rs1981037805 | <sup>7</sup> | This study |
| 33 | c.99G>T | R33S | rs771964597 | <sup>1</sup> | This study |
| 34 | c.100T>C | S34P | rs376479233 | <sup>42</sup> | This study |
| 34 | c.101C>T | S34F | rs774614463 | None found | This study |
| 35 | c.103C>T | P35S | rs771113232 | None found | This study |
| 36 | c.106C>T | R36C | rs372398235 | 22,42,54,75,76 | <sup>77</sup>, This study |
| 36 | c.107G>A | R36H | rs199473351 | 1,12,20,28,38,47,59,78-83 | <sup>79</sup>, This study |
| 37 | c.110G>A | S37N | rs1290531137 | None found | This study |
| 40 | c.118G>A | G40S | rs778439872 | <sup>1</sup> | This study |
| 41 | c.123G>C | K41N | rs763754260 | <sup>7</sup> | <sup>84</sup>, This study |
| 43 | c.127G>A | E43K | rs755781709 | 1,85 | This study |
| 44 | c.130G>A | A44T | rs767315445 | None found | This study |
| 46 | c.137A>G | Y46C | rs1402178514 | <sup>7</sup> | <sup>63,64,77,86,87</sup>, This study |
| 46 | c.138C>A | Y46X | rs758346045 | <sup>41</sup> | This study |
| 47 | c.139G>A | V47I | rs199473353 | 1,53,88-93 | <sup>94</sup>, This study |
| 47 | c.139G>T | V47F | rs199473353 | 3,6,7,12,95 | <sup>95</sup>, This study |
| 48 | c.142C>A | L48I | rs75610894 | None found | This study |
| 48 | c.142C>T | L48F | rs75610894 | 1,12,22,42,96-98 | This study |
| 49 | c.147G>A | M49I | rs773365460 | <sup>53</sup> | This study |
| 51 | c.152_153delTGinsAT | L51H | Not found | 3,6,7,12,95 | <sup>17,95,99-102</sup>, This study |
| 52 | c.154G>A | G52R | rs199473354 | 12,103,104 | <sup>103,105-110</sup>, This study |
| 53 | c.158T>C | F53S | rs199473355 | 12,47,80 | <sup>18</sup>, This study |
| 53 | c.158T>G | F53C | rs199473355 | <sup>7</sup> | <sup>63,87,111</sup>, This study |
| 53 | c.159C>A | F53L | rs1981011440 | None found | This study |
| 54 | c.160T>G | F54V | Not found | <sup>112,113</sup> | This study |
| 55 | c.163G>A | G55S | rs199473644 | <sup>1,7,12,14,23,28,29,38,47,69</sup> | This study |
| 58 | c.172A>C | T58P | rs147187721 | <sup>12,29,47</sup> | This study |
| 58 | c.173C>T | T58I | rs747321794 | <sup>41</sup> | <sup>114</sup>, This study |
| 58.5 | c.172_177delACCCTGinsCCCCCT | T58P-L59P | rs281865421 | <sup>2,7,115-118</sup> | <sup>106,119</sup>, This study |
| 59 | c.176T>C | L59P | rs141813529 | <sup>12,29,47,115</sup> | This study |
| 60 | c.179G>A | G60D | rs758961135 | <sup>49,50,120-123</sup> | <sup>49,50</sup>, This study |
| 61 | c.181A>G | I61V | rs143661740 | <sup>7,22,42</sup> | This study |
| 67 | c.199C>A | R67S | rs199473645 | <sup>115,117</sup> | This study |
| 67 | c.199C>G | R67G | rs199473645 | <sup>1,124</sup> | This study |
| 67 | c.199C>T | R67C | rs199473645 | <sup>1,7,12,14,29,47,51,125-128</sup> | <sup>51,63,86,129</sup>, This study |
| 67 | c.200G>A | R67H | rs79654911 | <sup>1,7,12,14,19,22,29,47,51,53,69,115,128,130-139</sup> | <sup>51,129</sup>, This study |
| 67 | c.200G>T | R67L | rs79654911 | <sup>7,41,140,141</sup> | This study |
| 69 | c.206A>G | K69R | rs149338401 | <sup>1,12,14,22,37,75,104,142</sup> | This study |
| 70 | c.208A>C | K70Q | rs1568836235 | None found | <sup>143</sup>, This study |
| 70 | c.209A>T | K70M | rs199473646 | <sup>7,12,29,47,144</sup> | <sup>129</sup>, This study |
| 70 | c.210G>C | K70N | rs199473356 | <sup>12,145-147</sup> | <sup>129,148,149</sup>, This study |
| 74 | c.220T>C | S74P | rs199473357 | <sup>1</sup> | This study |
| 74 | c.221C>G | S74W | rs74315446 | None found | This study |
| 74 | c.221C>T | S74L | rs74315446 | <sup>1-3,6,7,12,14,31,51,74,104,115,150-154</sup> | <sup>51,106,143,149,154-159</sup>, This study |
| 76 | c.226G>A | D76N | rs74315445 | <sup>1-3,6-8,12,14,19,22,23,29,51,53,55,69,115-117,120,128,139,151-154,160-181</sup> | <sup>18,33,95,143,154-159,181-192</sup>, This study |
| 80 | c.238G>A | V80I | rs769368494 | <sup>1,7,193,194</sup> | This study |
| 80 | c.238G>C | V80L | rs769368494 | None found | This study |
| 81 | c.242A>G | Y81C | rs199473359 | <sup>12,47,145,147,195</sup> | <sup>63,196,197</sup>, This study |
| 82 | c.244A>G | I82V | Not found | <sup>7</sup> | This study |
| 82 | c.246C>G | I82M | rs555086287 | None found | This study |
| 83 | c.247G>A | E83K | rs199473360 | <sup>1,12,14,23,29,38,47,122</sup> | This study |
| 87 | c.259T>C | W87R | rs199473361 | <sup>3,6,12,95,193</sup> | <sup>18,95,189,191</sup>, This study |
| 88 | c.262C>A | Q88K | rs750349088 | <sup>1</sup> | This study |
| 91 | c.271G>A | D91N | rs753805986 | None found | This study |
| 91 | c.273C>A | D91E | rs146614850 | 22,42,198,199 | This study |
| 93 | c.277G>A | A93T | rs1310229049 | 120,122,200 | This study |
| 95 | c.283G>A | V95I | rs772405174 | <sup>1</sup> | This study |
| 96 | c.287A>G | Q96R | rs767972997 | 22,68 | This study |
| 98 | c.292C>G | R98G | rs199473362 | None found | This study |
| 98 | c.292C>T | R98W | rs199473362 | 1,3,6,7,11,12,14,27,32,37,51,72,104,115,116,135,138,151,160,161,195,201-205 | 32,106, This study |
| 98 | c.293G>A | R98Q | rs150454912 | 1,7,22,39,42,179,206 | <sup>129</sup>, This study |
| 99 | c.295G>C | V99L | Not found | <sup>7</sup> | This study |
| 105 | c.314C>T | S105L | rs780041404 | 54,76,122,140,207,208 | This study |
| 109 | c.325G>A | V109I | rs77442996 | 1,3,6,7,11,12,14,19,20,22,23,28,36,46,59,122,142,151,209-211 | 210,212, This study |
| 112 | c.335A>G | H112R | rs1352531511 | None found | This study |
| 114 | c.340G>A | A114T | rs754020878 | None found | This study |
| 116 | c.347A>C | E116A | rs1235182008 | None found | This study |
| 117 | c.349C>T | Q117X | rs764130990 | <sup>29</sup> | This study |
| 120 | c.359C>T | T120I | rs1568835425 | None found | This study |
| 125 | c.374C>T | T125M | rs142511345 | 1,7,12,14,19,20,22,28,29,38,47,59,213,214 | This study |
| 127 | c.379C>A | P127T | rs199473647 | 3,6,12,59,66,69,72,74,151 | 74,212,215, This study |

## References

1. George AL, Jr. Molecular and genetic basis of sudden cardiac death. J Clin Invest. 2013;123:75–83.

2. Ackerman M, Atkins DL and Triedman JK. Sudden cardiac death in the young. Circulation. 2016;133:1006–26.

3. Schwartz PJ, Ackerman MJ, Antzelevitch C, et al. Inherited cardiac arrhythmias. Nat Rev Dis Primers. 2020;6:58.

4. Schwartz PJ, Crotti L and Insolia R. Long-QT syndrome: from genetics to management. Circ Arrhythm Electrophysiol. 2012;5:868–877.

5. Sanguinetti MC, Curran ME, Zou A, et al. Coassembly of Kv LQT1 and minK (IsK) proteins to form cardiac I_Ks_ potassium channel. Nature. 1996;384:80–83.

6. Barhanin J, Lesage F, Guillemare E, et al. KvLQT1 and IsK (minK) proteins associate to form the I_Ks_ cardiac potassium current. Nature. 1996;384:78–80.

7. Adler A, Novelli V, Amin AS, et al. An international, multicentered, evidence-based reappraisal of genes reported to cause congenital long QT syndrome. Circulation. 2020;141:418–428.

8. Schulze-Bahr E, Wang Q, Wedekind H, et al. KCNE1 mutations cause Jervell and Lange-Nielsen syndrome. Nature Genet. 1997;17:267–268.

9. Duggal P, Vesely MR, Wattanasirichaigoon D, et al. Mutation of the gene for IsK associated with both Jervell and Lange-Nielsen and Functional Profiling of KCNE1 Variants Romano-Ward forms of Long-QT syndrome. Circulation. 1998;97:142–146.

10. Roberts JD, Asaki SY, Mazzanti A, et al. An international multicenter evaluation of type 5 long QT syndrome: A low penetrant primary arrhythmic condition. Circulation. 2020;141:429–439.

11. Karczewski KJ, Francioli LC, Tiao G, et al. The mutational constraint spectrum quantified from variation in 141,456 humans. Nature. 2020;581:434–443.

12. Garmany R, Giudicessi JR, Ye D, et al. Clinical and functional reappraisal of alleged type 5 long QT syndrome: Causative genetic variants in the KCNE1-encoded minK β-subunit. Heart Rhythm. 2020;17:937–944.

13. Vanoye CG, Desai RR, Fabre KL, et al. High-throughput functional evaluation of KCNQ1 decrypts variants of unknown significance. Circ Genom Precis Med. 2018;11:e002345.

14. Vanoye CG, Thompson CH, Desai RR, et al. Functional evaluation of human ion channel variants using automated electrophysiology. Methods Enzymol. 2021;654:383–405.

15. Glazer AM, Wada Y, Li B, et al. High-throughput reclassification of SCN5A variants. Am J Hum Genet. 2020;107:111–123.

16. Jiang C, Richardson E, Farr J, et al. A calibrated functional patch-clamp assay to enhance clinical variant interpretation in KCNH2-related long QT syndrome. Am J Hum Genet. 2022;109:1199–1207.

17. O’Neill MJ, Ng CA, Aizawa T, et al. Multiplexed assays of variant effect and automated patch clamping improve KCNH2-LQTS variant classification and cardiac event risk stratification. Circulation. 2024;150:1869–1881.

18. Huang H, Kuenze G, Smith JA, et al. Mechanisms of KCNQ1 channel dysfunction in long QT syndrome involving voltage sensor domain mutations. Sci Adv. 2018;4:eaar2631.

19. Coyote-Maestas W, Nedrud D, Suma A, et al. Probing ion channel functional architecture and domain recombination compatibility by massively parallel domain insertion profiling. Nature Commun. 2021;12:7114.

20. Glazer AM, Kroncke BM, Matreyek KA, et al. Deep mutational ccan of an SCN5A voltage sensor. Circ Genom Precis Med. 2020;13:e002786.

21. Muhammad A, Calandranis ME, Li B, et al. High-throughput functional mapping of variants in an arrhythmia gene, KCNE1, reveals novel biology. Genome Med. 2024;16:73.

22. Towart R, Linders JT, Hermans AN, et al. Blockade of the I_Ks_ potassium channel: An overlooked cardiovascular liability in drug safety screening? J Pharmacol Toxicol Methods. 2009.

23. Richards S, Aziz N, Bale S, et al. Standards and guidelines for the interpretation of sequence variants: a joint consensus recommendation of the American College of Medical Genetics and Genomics and the Association for Molecular Pathology. Genet Med. 2015;17:405–424.

24. Landrum MJ, Lee JM, Riley GR, et al. ClinVar: public archive of relationships among sequence variation and human phenotype. Nucleic Acids Res. 2014;42:D980–D985.

25. Li Q and Wang K. InterVar: Clinical interpretation of genetic variants by the 2015 ACMG-AMP guidelines. Am J Hum Genet. 2017;100:267–280.

26. Harrison PW, Amode MR, Austine-Orimoloye O, et al. Ensembl 2024. Nucleic Acids Res. 2024;52:D891–d899.

27. Abou Tayoun AN, Pesaran T, DiStefano MT, et al. Recommendations for interpreting the loss of function PVS1 ACMG/AMP variant criterion. Hum Mutat. 2018;39:1517–1524.

28. Jarvik GP and Browning BL. Consideration of cosegregation in the pathogenicity classification of genomic variants. Am J Hum Genet. 2016;98:1077–1081.

29. Ng PC and Henikoff S. SIFT: Predicting amino acid changes that affect protein function. Nucleic Acids Res. 2003;31:3812–3814.

30. Adzhubei I, Jordan DM and Sunyaev SR. Predicting functional effect of human missense mutations using PolyPhen-2. Curr Protoc Hum Genet. 2013;76:7.20.1-7.20.41.

31. Appadurai V, DeBarber A, Chiang PW, et al. Apparent underdiagnosis of Cerebrotendinous Xanthomatosis revealed by analysis of ∼60,000 human exomes. Mol Genet Metab. 2015;116:298–304.

32. Guo MH and Gregg AR. Estimating yields of prenatal carrier screening and implications for design of expanded carrier screening panels. Genet Med. 2019;21:1940–1947.

33. Stenson PD, Mort M, Ball EV, et al. The Human Gene Mutation Database: towards a comprehensive repository of inherited mutation data for medical research, genetic diagnosis and next-generation sequencing studies. Hum Genet. 2017;136:665–677.

34. Takumi T, Ohkubo H and Nakanishi S. Cloning of a membrane protein that induces a slow voltage-gated potassium current. Science. 1988;242:1042–1045.

35. Kaczmarek LK. Voltage-dependent potassium channels: minK and Shaker families. New Biol. 1991;3:315–23.

36. Wang Q, Curran ME, Splawski I, et al. Positional cloning of a novel potassium channel gene: KVLQT1 mutations cause cardiac arrhythmias. Nature Genet. 1996;12:17–23.

37. Kapplinger JD, Tester DJ, Salisbury BA, et al. Spectrum and prevalence of mutations from the first 2,500 consecutive unrelated patients referred for the FAMILION long QT syndrome genetic test. Heart Rhythm. 2009;6:1297–1303.

38. Kaab S, Crawford DC, Sinner MF, et al. A large candidate gene survey identifies the KCNE1 D85N polymorphism as a possible modulator of drug-induced torsades de pointes. Circ Cardiovasc Genet. 2011;5:91–99.

39. McDonald TV, Yu Z, Ming Z, et al. A minK-HERG complex regulates the cardiac potassium current I_Kr_. Nature. 1997;388:289–292.

## References for Supplemental Table S3

1. Glazer, A.M., Davogustto, G., Shaffer, C.M., Vanoye, C.G., Desai, R.R., Farber-Eger, E.H., Dikilitas, O., Shang, N., Pacheco, J.A., Yang, T., et al. (2022). Arrhythmia Variant Associations and Reclassifications in the eMERGE-III Sequencing Study. Circulation 145, 877–891. 10.1161/circulationaha.121.055562.

2. Gaunt, T.R., Shah, S., Nelson, C.P., Drenos, F., Braund, P.S., Adeniran, I., Folkersen, L., Lawlor, D.A., Casas, J.P., Amuzu, A., et al. (2012). Integration of genetics into a systems model of electrocardiographic traits using HumanCVD BeadChip. Circ Cardiovasc Genet 5, 630–638. 10.1161/circgenetics.112.962852.

3. Holehouse, A.S., and Naegle, K.M. (2015). Reproducible Analysis of Post-Translational Modifications in Proteomes--Application to Human Mutations. PLoS One 10, e0144692. 10.1371/journal.pone.0144692.

4. Lavrov, A.V., Varenikov, G.G., and Skoblov, M.Y. (2020). Genome scale analysis of pathogenic variants targetable for single base editing. BMC Med Genomics 13, 80. 10.1186/s12920-020-00735-8.

5. Mazumder, R., Morampudi, K.S., Motwani, M., Vasudevan, S., and Goldman, R. (2012). Proteome-wide analysis of single-nucleotide variations in the N-glycosylation sequon of human genes. PLoS One 7, e36212. 10.1371/journal.pone.0036212.

6. Molnár, J., Szakács, G., and Tusnády, G.E. (2016). Characterization of Disease-Associated Mutations in Human Transmembrane Proteins. PLoS One 11, e0151760. 10.1371/journal.pone.0151760.

7. Roberts, J.D., Asaki, S.Y., Mazzanti, A., Bos, J.M., Tuleta, I., Muir, A.R., Crotti, L., Krahn, A.D., Kutyifa, V., Shoemaker, M.B., et al. (2020). An International Multicenter Evaluation of Type 5 Long QT Syndrome: A Low Penetrant Primary Arrhythmic Condition. Circulation 141, 429–439. 10.1161/circulationaha.119.043114.

8. Schulze-Bahr, E., Wang, Q., Wedekind, H., Haverkamp, W., Chen, Q., Sun, Y., Rubie, C., Hördt, M., Towbin, J.A., Borggrefe, M., et al. (1997). KCNE1 mutations cause jervell and Lange-Nielsen syndrome. Nat Genet 17, 267–268. 10.1038/ng1197-267.

9. Simpson, C.M., Zhang, B., Hornbeck, P.V., and Gnad, F. (2019). Systematic analysis of the intersection of disease mutations with protein modifications. BMC Med Genomics 12, 109. 10.1186/s12920-019-0543-2.

10. Sudandiradoss, C., and Sethumadhavan, R. (2008). In silico investigations on functional and haplotype tag SNPs associated with congenital long QT syndromes (LQTSs). Genomic Med 2, 55–67. 10.1007/s11568-009-9027-3.

11. Sun, B., Zhang, M., Cui, P., Li, H., Jia, J., Li, Y., and Xie, L. (2015). Nonsynonymous Single-Nucleotide Variations on Some Posttranslational Modifications of Human Proteins and the Association with Diseases. Comput Math Methods Med 2015, 124630. 10.1155/2015/124630.

12. Ware, J.S., Walsh, R., Cunningham, F., Birney, E., and Cook, S.A. (2012). Paralogous annotation of disease-causing variants in long QT syndrome genes. Hum Mutat 33, 1188–1191. 10.1002/humu.22114.

13. Won, H.H., Kim, H.J., Lee, K.A., and Kim, J.W. (2008). Cataloging coding sequence variations in human genome databases. PLoS One 3, e3575. 10.1371/journal.pone.0003575.

14. Yang, S., Lincoln, S.E., Kobayashi, Y., Nykamp, K., Nussbaum, R.L., and Topper, S. (2017). Sources of discordance among germ-line variant classifications in ClinVar. Genet Med 19, 1118–1126. 10.1038/gim.2017.60.

15. Ávalos Prado, P., Häfner, S., Comoglio, Y., Wdziekonski, B., Duranton, C., Attali, B., Barhanin, J., and Sandoz, G. (2021). KCNE1 is an auxiliary subunit of two distinct ion channel superfamilies. Cell 184, 534-544.e511. 10.1016/j.cell.2020.11.047.

16. Bas, T., Gao, G.Y., Lvov, A., Chandrasekhar, K.D., Gilmore, R., and Kobertz, W.R. (2011). Post-translational N-glycosylation of type I transmembrane KCNE1 peptides: implications for membrane protein biogenesis and disease. J Biol Chem 286, 28150–28159. 10.1074/jbc.M111.235168.

17. David, J.P., Andersen, M.N., Olesen, S.P., Rasmussen, H.B., and Schmitt, N. (2013). Trafficking of the IKs -complex in MDCK cells: site of subunit assembly and determinants of polarized localization. Traffic 14, 399–411. 10.1111/tra.12042.

18. Sahni, N., Yi, S., Taipale, M., Fuxman Bass, J.I., Coulombe-Huntington, J., Yang, F., Peng, J., Weile, J., Karras, G.I., Wang, Y., et al. (2015). Widespread macromolecular interaction perturbations in human genetic disorders. Cell 161, 647–660. 10.1016/j.cell.2015.04.013.

19. Amendola, L.M., Dorschner, M.O., Robertson, P.D., Salama, J.S., Hart, R., Shirts, B.H., Murray, M.L., Tokita, M.J., Gallego, C.J., Kim, D.S., et al. (2015). Actionable exomic incidental findings in 6503 participants: challenges of variant classification. Genome Res 25, 305–315. 10.1101/gr.183483.114.

20. Azevedo, L., Mort, M., Costa, A.C., Silva, R.M., Quelhas, D., Amorim, A., and Cooper, D.N. (2016). Improving the in silico assessment of pathogenicity for compensated variants. Eur J Hum Genet 25, 2–7. 10.1038/ejhg.2016.129.

21. Bennett, J.S., Bernhardt, M., McBride, K.L., Reshmi, S.C., Zmuda, E., Kertesz, N.J., Garg, V., Fitzgerald-Butt, S., and Kamp, A.N. (2019). Reclassification of Variants of Uncertain Significance in Children with Inherited Arrhythmia Syndromes is Predicted by Clinical Factors. Pediatr Cardiol 40, 1679–1687. 10.1007/s00246-019-02203-2.

22. Darbro, B.W., Singh, R., Zimmerman, M.B., Mahajan, V.B., and Bassuk, A.G. (2016). Autism Linked to Increased Oncogene Mutations but Decreased Cancer Rate. PLoS One 11, e0149041. 10.1371/journal.pone.0149041.

23. Famiglietti, M.L., Estreicher, A., Breuza, L., Poux, S., Redaschi, N., Xenarios, I., and Bridge, A. (2019). An enhanced workflow for variant interpretation in UniProtKB/Swiss-Prot improves consistency and reuse in ClinVar. Database (Oxford) 2019. 10.1093/database/baz040.

24. Fedida, J., Fressart, V., Charron, P., Surget, E., Hery, T., Richard, P., Donal, E., Keren, B., Duthoit, G., Hidden-Lucet, F., et al. (2017). Contribution of exome sequencing for genetic diagnostic in arrhythmogenic right ventricular cardiomyopathy/dysplasia. PLoS One 12, e0181840. 10.1371/journal.pone.0181840.

25. Gao, Y., Wang, H.Y., Guan, J., Lan, L., Zhao, C., Xie, L.Y., Wang, D.Y., and Wang, Q.J. (2021). Genetic Susceptibility Study of Chinese Sudden Sensorineural Hearing Loss Patients with Vertigo. Curr Med Sci 41, 673–679. 10.1007/s11596-021-2422-2.

26. Itoh, H., Crotti, L., Aiba, T., Spazzolini, C., Denjoy, I., Fressart, V., Hayashi, K., Nakajima, T., Ohno, S., Makiyama, T., et al. (2016). The genetics underlying acquired long QT syndrome: impact for genetic screening. Eur Heart J 37, 1456–1464. 10.1093/eurheartj/ehv695.

27. Itoh, H., Shimizu, W., Hayashi, K., Yamagata, K., Sakaguchi, T., Ohno, S., Makiyama, T., Akao, M., Ai, T., Noda, T., et al. (2010). Long QT syndrome with compound mutations is associated with a more severe phenotype: a Japanese multicenter study. Heart Rhythm 7, 1411–1418. 10.1016/j.hrthm.2010.06.013.

28. Kaltman, J.R., Evans, F., and Fu, Y.P. (2018). Re-evaluating pathogenicity of variants associated with the long QT syndrome. J Cardiovasc Electrophysiol 29, 98–104. 10.1111/jce.13355.

29. Kapplinger, J.D., Tester, D.J., Salisbury, B.A., Carr, J.L., Harris-Kerr, C., Pollevick, G.D., Wilde, A.A., and Ackerman, M.J. (2009). Spectrum and prevalence of mutations from the first 2,500 consecutive unrelated patients referred for the FAMILION long QT syndrome genetic test. Heart Rhythm 6, 1297–1303. 10.1016/j.hrthm.2009.05.021.

30. Le Scouarnec, S., Karakachoff, M., Gourraud, J.B., Lindenbaum, P., Bonnaud, S., Portero, V., Duboscq-Bidot, L., Daumy, X., Simonet, F., Teusan, R., et al. (2015). Testing the burden of rare variation in arrhythmia-susceptibility genes provides new insights into molecular diagnosis for Brugada syndrome. Hum Mol Genet 24, 2757–2763. 10.1093/hmg/ddv036.

31. Li, Z., Chen, P., Li, C., Tan, L., Xu, J., Wang, H., Sun, Y., Wang, Y., Zhao, C., Link, M.S., et al. (2020). Genetic arrhythmias complicating patients with dilated cardiomyopathy. Heart Rhythm 17, 305–312. 10.1016/j.hrthm.2019.09.012.

32. Ohno, S., Zankov, D.P., Yoshida, H., Tsuji, K., Makiyama, T., Itoh, H., Akao, M., Hancox, J.C., Kita, T., and Horie, M. (2007). N- and C-terminal KCNE1 mutations cause distinct phenotypes of long QT syndrome. Heart Rhythm 4, 332–340. 10.1016/j.hrthm.2006.11.004.

33. Du, C., El Harchi, A., Zhang, H., and Hancox, J.C. (2013). Modification by KCNE1 variants of the hERG potassium channel response to premature stimulation and to pharmacological inhibition. Physiol Rep 1, e00175. 10.1002/phy2.175.

34. Di Resta, C., Pietrelli, A., Sala, S., Della Bella, P., De Bellis, G., Ferrari, M., Bordoni, R., and Benedetti, S. (2015). High-throughput genetic characterization of a cohort of Brugada syndrome patients. Hum Mol Genet 24, 5828–5835. 10.1093/hmg/ddv302.

35. Kauferstein, S., Herz, N., Scheiper, S., Biel, S., Jenewein, T., Kunis, M., Erkapic, D., Beckmann, B.M., and Neumann, T. (2017). Relevance of molecular testing in patients with a family history of sudden death. Forensic Sci Int 276, 18–23. 10.1016/j.forsciint.2017.04.001.

36. Mahdieh, N., Khorgami, M., Soveizi, M., Seyed Aliakbar, S., Dalili, M., and Rabbani, B. (2020). Genetic homozygosity in a diverse population: An experience of long QT syndrome. Int J Cardiol 316, 117–124. 10.1016/j.ijcard.2020.05.056.

37. Ng, D., Johnston, J.J., Teer, J.K., Singh, L.N., Peller, L.C., Wynter, J.S., Lewis, K.L., Cooper, D.N., Stenson, P.D., Mullikin, J.C., and Biesecker, L.G. (2013). Interpreting secondary cardiac disease variants in an exome cohort. Circ Cardiovasc Genet 6, 337–346. 10.1161/circgenetics.113.000039.

38. Shakeel, M., Irfan, M., and Khan, I.A. (2018). Estimating the mutational load for cardiovascular diseases in Pakistani population. PLoS One 13, e0192446. 10.1371/journal.pone.0192446.

39. Stattin, E.L., Boström, I.M., Winbo, A., Cederquist, K., Jonasson, J., Jonsson, B.A., Diamant, U.B., Jensen, S.M., Rydberg, A., and Norberg, A. (2012). Founder mutations characterise the mutation panorama in 200 Swedish index cases referred for Long QT syndrome genetic testing. BMC Cardiovasc Disord 12, 95. 10.1186/1471-2261-12-95.

40. Yoshinaga, M., Kucho, Y., Sarantuya, J., Ninomiya, Y., Horigome, H., Ushinohama, H., Shimizu, W., and Horie, M. (2014). Genetic characteristics of children and adolescents with long-QT syndrome diagnosed by school-based electrocardiographic screening programs. Circ Arrhythm Electrophysiol 7, 107–112. 10.1161/CIRCEP.113.000426.

41. Faridi, R., Tona, R., Brofferio, A., Hoa, M., Olszewski, R., Schrauwen, I., Assir, M.Z.K., Bandesha, A.A., Khan, A.A., Rehman, A.U., et al. (2019). Mutational and phenotypic spectra of KCNE1 deficiency in Jervell and Lange-Nielsen Syndrome and Romano-Ward Syndrome. Hum Mutat 40, 162–176. 10.1002/humu.23689.

42. Huang, J., Wang, X., Hao, B., Chen, Y., Liu, H., Quan, L., Tang, D., Sheng, L., Li, M., Huang, E., et al. (2015). Genetic variants in KCNE1, KCNQ1, and NOS1AP in sudden unexplained death during daily activities in Chinese Han population. J Forensic Sci 60, 351–356. 10.1111/1556-4029.12687.

43. Sun, P., Wu, F., Wen, M., Yang, X., Wang, C., Li, Y., He, S., Zhang, L., Zhang, Y., and Tian, C. (2015). A distinct three-helix centipede toxin SSD609 inhibits I(ks) channels by interacting with the KCNE1 auxiliary subunit. Sci Rep 5, 13399. 10.1038/srep13399.

44. Huang, H., Millat, G., Rodriguez-Lafrasse, C., Rousson, R., Kugener, B., Chevalier, P., and Chahine, M. (2009). Biophysical characterization of a new SCN5A mutation S1333Y in a SIDS infant linked to long QT syndrome. FEBS Lett 583, 890–896. 10.1016/j.febslet.2009.02.007.

45. Millat, G., Kugener, B., Chevalier, P., Chahine, M., Huang, H., Malicier, D., Rodriguez-Lafrasse, C., and Rousson, R. (2009). Contribution of long-QT syndrome genetic variants in sudden infant death syndrome. Pediatr Cardiol 30, 502–509. 10.1007/s00246-009-9417-2.

46. Paludan-Müller, C., Ghouse, J., Vad, O.B., Herfelt, C.B., Lundegaard, P., Ahlberg, G., Schmitt, N., Svendsen, J.H., Haunsø, S., Bundgaard, H., et al. (2019). Reappraisal of variants previously linked with sudden infant death syndrome: results from three population-based cohorts. Eur J Hum Genet 27, 1427–1435. 10.1038/s41431-019-0416-3.

47. Takeda, J.I., Nanatsue, K., Yamagishi, R., Ito, M., Haga, N., Hirata, H., Ogi, T., and Ohno, K. (2020). InMeRF: prediction of pathogenicity of missense variants by individual modeling for each amino acid substitution. NAR Genom Bioinform 2, lqaa038. 10.1093/nargab/lqaa038.

48. Andreasen, L., Ahlberg, G., Tang, C., Andreasen, C., Hartmann, J.P., Tfelt-Hansen, J., Behr, E.R., Pehrson, S., Haunsø, S., LuCamp, et al. (2018). Next-generation sequencing of AV nodal reentrant tachycardia patients identifies broad spectrum of variants in ion channel genes. Eur J Hum Genet 26, 660–668. 10.1038/s41431-017-0092-0.

49. Olesen, M.S., Andreasen, L., Jabbari, J., Refsgaard, L., Haunsø, S., Olesen, S.P., Nielsen, J.B., Schmitt, N., and Svendsen, J.H. (2014). Very early-onset lone atrial fibrillation patients have a high prevalence of rare variants in genes previously associated with atrial fibrillation. Heart Rhythm 11, 246–251. 10.1016/j.hrthm.2013.10.034.

50. Olesen, M.S., Bentzen, B.H., Nielsen, J.B., Steffensen, A.B., David, J.P., Jabbari, J., Jensen, H.K., Haunsø, S., Svendsen, J.H., and Schmitt, N. (2012). Mutations in the potassium channel subunit KCNE1 are associated with early-onset familial atrial fibrillation. BMC Med Genet 13, 24. 10.1186/1471-2350-13-24.

51. Garmany, R., Giudicessi, J.R., Ye, D., Zhou, W., Tester, D.J., and Ackerman, M.J. (2020). Clinical and functional reappraisal of alleged type 5 long QT syndrome: Causative genetic variants in the KCNE1-encoded minK β-subunit. Heart Rhythm 17, 937–944. 10.1016/j.hrthm.2020.02.003.

52. Ichikawa, M., Aiba, T., Ohno, S., Shigemizu, D., Ozawa, J., Sonoda, K., Fukuyama, M., Itoh, H., Miyamoto, Y., Tsunoda, T., et al. (2016). Phenotypic Variability of ANK2 Mutations in Patients With Inherited Primary Arrhythmia Syndromes. Circ J 80, 2435–2442. 10.1253/circj.CJ-16-0486.

53. Lieve, K.V., Williams, L., Daly, A., Richard, G., Bale, S., Macaya, D., and Chung, W.K. (2013). Results of genetic testing in 855 consecutive unrelated patients referred for long QT syndrome in a clinical laboratory. Genet Test Mol Biomarkers 17, 553–561. 10.1089/gtmb.2012.0118.

54. Lopes, L.R., Syrris, P., Guttmann, O.P., O’Mahony, C., Tang, H.C., Dalageorgou, C., Jenkins, S., Hubank, M., Monserrat, L., McKenna, W.J., et al. (2015). Novel genotype-phenotype associations demonstrated by high-throughput sequencing in patients with hypertrophic cardiomyopathy. Heart 101, 294–301. 10.1136/heartjnl-2014-306387.

55. Mitchell, J.L., Cuneo, B.F., Etheridge, S.P., Horigome, H., Weng, H.Y., and Benson, D.W. (2012). Fetal heart rate predictors of long QT syndrome. Circulation 126, 2688–2695. 10.1161/circulationaha.112.114132.

56. Obeyesekere, M.N., Sy, R.W., Klein, G.J., Gula, L.J., Modi, S., Conacher, S., Leong-Sit, P., Skanes, A.C., Yee, R., and Krahn, A.D. (2012). End-recovery QTc: a useful metric for assessing genetic variants of unknown significance in long-QT syndrome. J Cardiovasc Electrophysiol 23, 637–642. 10.1111/j.1540-8167.2011.02265.x.

57. Robinson, J.A., Bos, J.M., Etheridge, S.P., and Ackerman, M.J. (2015). Breath Holding Spells in Children with Long QT Syndrome. Congenit Heart Dis 10, 354–361. 10.1111/chd.12262.

58. Rohatgi, R.K., Bos, J.M., and Ackerman, M.J. (2015). Stimulant therapy in children with attention-deficit/hyperactivity disorder and concomitant long QT syndrome: A safe combination? Heart Rhythm 12, 1807–1812. 10.1016/j.hrthm.2015.04.043.

59. Schneider, V.A., Graves-Lindsay, T., Howe, K., Bouk, N., Chen, H.C., Kitts, P.A., Murphy, T.D., Pruitt, K.D., Thibaud-Nissen, F., Albracht, D., et al. (2017). Evaluation of GRCh38 and de novo haploid genome assemblies demonstrates the enduring quality of the reference assembly. Genome Res 27, 849–864. 10.1101/gr.213611.116.

60. Schwartz, P.J., Stramba-Badiale, M., Crotti, L., Pedrazzini, M., Besana, A., Bosi, G., Gabbarini, F., Goulene, K., Insolia, R., Mannarino, S., et al. (2009). Prevalence of the congenital long-QT syndrome. Circulation 120, 1761–1767. 10.1161/circulationaha.109.863209.

61. Shim, S.H., Ito, M., Maher, T., and Milunsky, A. (2005). Gene sequencing in neonates and infants with the long QT syndrome. Genet Test 9, 281–284. 10.1089/gte.2005.9.281.

62. McGorrian, C., Constant, O., Harper, N., O’Donnell, C., Codd, M., Keelan, E., Green, A., O’Neill, J., Galvin, J., and Mahon, N.G. (2013). Family-based cardiac screening in relatives of victims of sudden arrhythmic death syndrome. Europace 15, 1050–1058. 10.1093/europace/eus408.

63. Sahu, I.D., Craig, A.F., Dunagan, M.M., Troxel, K.R., Zhang, R., Meiberg, A.G., Harmon, C.N., McCarrick, R.M., Kroncke, B.M., Sanders, C.R., and Lorigan, G.A. (2015). Probing Structural Dynamics and Topology of the KCNE1 Membrane Protein in Lipid Bilayers via Site-Directed Spin Labeling and Electron Paramagnetic Resonance Spectroscopy. Biochemistry 54, 6402–6412. 10.1021/acs.biochem.5b00505.

64. Xu, Y., Wang, Y., Meng, X.Y., Zhang, M., Jiang, M., Cui, M., and Tseng, G.N. (2013). Building KCNQ1/KCNE1 channel models and probing their interactions by molecular-dynamics simulations. Biophys J 105, 2461–2473. 10.1016/j.bpj.2013.09.058.

65. Berge, K.E., Haugaa, K.H., Früh, A., Anfinsen, O.G., Gjesdal, K., Siem, G., Oyen, N., Greve, G., Carlsson, A., Rognum, T.O., et al. (2008). Molecular genetic analysis of long QT syndrome in Norway indicating a high prevalence of heterozygous mutation carriers. Scand J Clin Lab Invest 68, 362–368. 10.1080/00365510701765643.

66. Boudellioua, I., Kulmanov, M., Schofield, P.N., Gkoutos, G.V., and Hoehndorf, R. (2018). OligoPVP: Phenotype-driven analysis of individual genomic information to prioritize oligogenic disease variants. Sci Rep 8, 14681. 10.1038/s41598-018-32876-3.

67. Hasdemir, C., Payzin, S., Kocabas, U., Sahin, H., Yildirim, N., Alp, A., Aydin, M., Pfeiffer, R., Burashnikov, E., Wu, Y., and Antzelevitch, C. (2015). High prevalence of concealed Brugada syndrome in patients with atrioventricular nodal reentrant tachycardia. Heart Rhythm 12, 1584–1594. 10.1016/j.hrthm.2015.03.015.

68. Leinonen, J.T., Crotti, L., Djupsjöbacka, A., Castelletti, S., Junna, N., Ghidoni, A., Tuiskula, A.M., Spazzolini, C., Dagradi, F., Viitasalo, M., et al. (2018). The genetics underlying idiopathic ventricular fibrillation: A special role for catecholaminergic polymorphic ventricular tachycardia? Int J Cardiol 250, 139–145. 10.1016/j.ijcard.2017.10.016.

69. Mullally, J., Goldenberg, I., Moss, A.J., Lopes, C.M., Ackerman, M.J., Zareba, W., McNitt, S., Robinson, J.L., Benhorin, J., Kaufman, E.S., et al. (2013). Risk of life-threatening cardiac events among patients with long QT syndrome and multiple mutations. Heart Rhythm 10, 378–382. 10.1016/j.hrthm.2012.11.006.

70. Retterer, K., Juusola, J., Cho, M.T., Vitazka, P., Millan, F., Gibellini, F., Vertino-Bell, A., Smaoui, N., Neidich, J., Monaghan, K.G., et al. (2016). Clinical application of whole-exome sequencing across clinical indications. Genet Med 18, 696–704. 10.1038/gim.2015.148.

71. Seethala, S., Singh, P., Shusterman, V., Ribe, M., Haugaa, K.H., and Němec, J. (2015). QT Adaptation and Intrinsic QT Variability in Congenital Long QT Syndrome. J Am Heart Assoc 4. 10.1161/jaha.115.002395.

72. Splawski, I., Shen, J., Timothy, K.W., Lehmann, M.H., Priori, S., Robinson, J.L., Moss, A.J., Schwartz, P.J., Towbin, J.A., Vincent, G.M., and Keating, M.T. (2000). Spectrum of mutations in long-QT syndrome genes. KVLQT1, HERG, SCN5A, KCNE1, and KCNE2. Circulation 102, 1178–1185. 10.1161/01.cir.102.10.1178.

73. Wang, M., and Wei, L. (2016). iFish: predicting the pathogenicity of human nonsynonymous variants using gene-specific/family-specific attributes and classifiers. Sci Rep 6, 31321. 10.1038/srep31321.

74. Westenskow, P., Splawski, I., Timothy, K.W., Keating, M.T., and Sanguinetti, M.C. (2004). Compound mutations: a common cause of severe long-QT syndrome. Circulation 109, 1834–1841. 10.1161/01.Cir.0000125524.34234.13.

75. Florentine, M.M., Rouse, S.L., Stephans, J., Conrad, D., Czechowicz, J., Matthews, I.R., Meyer, A.K., Nadaraja, G.S., Parikh, R., Virbalas, J., et al. (2022). Racial and ethnic disparities in diagnostic efficacy of comprehensive genetic testing for sensorineural hearing loss. Hum Genet 141, 495–504. 10.1007/s00439-021-02338-4.

76. Lopes, L.R., Zekavati, A., Syrris, P., Hubank, M., Giambartolomei, C., Dalageorgou, C., Jenkins, S., McKenna, W., Plagnol, V., and Elliott, P.M. (2013). Genetic complexity in hypertrophic cardiomyopathy revealed by high-throughput sequencing. J Med Genet 50, 228–239. 10.1136/jmedgenet-2012-101270.

77. Wang, Y.H., Jiang, M., Xu, X.L., Hsu, K.L., Zhang, M., and Tseng, G.N. (2011). Gating-related molecular motions in the extracellular domain of the IKs channel: implications for IKs channelopathy. J Membr Biol 239, 137–156. 10.1007/s00232-010-9333-7.

78. Chen, P., He, L., Pang, X., Wang, X., Yang, T., and Wu, H. (2016). NLRP3 Is Expressed in the Spiral Ganglion Neurons and Associated with Both Syndromic and Nonsyndromic Sensorineural Deafness. Neural Plast 2016, 3018132. 10.1155/2016/3018132.

79. Lane, C.M., Giudicessi, J.R., Ye, D., Tester, D.J., Rohatgi, R.K., Bos, J.M., and Ackerman, M.J. (2018). Long QT syndrome type 5-Lite: Defining the clinical phenotype associated with the potentially proarrhythmic p.Asp85Asn-KCNE1 common genetic variant. Heart Rhythm 15, 1223–1230. 10.1016/j.hrthm.2018.03.038.

80. Napolitano, C., Priori, S.G., Schwartz, P.J., Bloise, R., Ronchetti, E., Nastoli, J., Bottelli, G., Cerrone, M., and Leonardi, S. (2005). Genetic testing in the long QT syndrome: development and validation of an efficient approach to genotyping in clinical practice. Jama 294, 2975–2980. 10.1001/jama.294.23.2975.

81. Rajan, S., Chockalingam, P., Koneti, N.R., Geetha, T.S., Mishra, S., and Narasimhan, C. (2021). Initial experience and results of a cardiogenetic clinic in a tertiary cardiac care center in India. Ann Pediatr Cardiol 14, 443–448. 10.4103/apc.apc_123_21.

82. Rashid, M., van der Horst, M., Mentzel, T., Butera, F., Ferreira, I., Pance, A., Rütten, A., Luzar, B., Marusic, Z., de Saint Aubain, N., et al. (2019). ALPK1 hotspot mutation as a driver of human spiradenoma and spiradenocarcinoma. Nat Commun 10, 2213. 10.1038/s41467-019-09979-0.

83. Salazar-Mendiguchía, J., Ochoa, J.P., Palomino-Doza, J., Domínguez, F., Díez-López, C., Akhtar, M., Ramiro-León, S., Clemente, M.M., Pérez-Cejas, A., Robledo, M., et al. (2020). Mutations in TRIM63 cause an autosomal-recessive form of hypertrophic cardiomyopathy. Heart 106, 1342–1348. 10.1136/heartjnl-2020-316913.

84. Barro-Soria, R., Ramentol, R., Liin, S.I., Perez, M.E., Kass, R.S., and Larsson, H.P. (2017). KCNE1 and KCNE3 modulate KCNQ1 channels by affecting different gating transitions. Proc Natl Acad Sci U S A 114, E7367–e7376. 10.1073/pnas.1710335114.

85. Sarquella-Brugada, G., García-Algar, O., Zambrano, M.D., Fernández-Falgueres, A., Sailer, S., Cesar, S., Sebastiani, G., Martí-Almor, J., Aurensanz, E., Cruzalegui, J.C., et al. (2021). Early Identification of Prolonged QT Interval for Prevention of Sudden Infant Death. Front Pediatr 9, 704580. 10.3389/fped.2021.704580.

86. Sahu, I.D., Kroncke, B.M., Zhang, R., Dunagan, M.M., Smith, H.J., Craig, A., McCarrick, R.M., Sanders, C.R., and Lorigan, G.A. (2014). Structural investigation of the transmembrane domain of KCNE1 in proteoliposomes. Biochemistry 53, 6392–6401. 10.1021/bi500943p.

87. Wang, Y., Zhang, M., Xu, Y., Jiang, M., Zankov, D.P., Cui, M., and Tseng, G.N. (2012). Probing the structural basis for differential KCNQ1 modulation by KCNE1 and KCNE2. J Gen Physiol 140, 653–669. 10.1085/jgp.201210847.

88. Azaiez, H., Booth, K.T., Ephraim, S.S., Crone, B., Black-Ziegelbein, E.A., Marini, R.J., Shearer, A.E., Sloan-Heggen, C.M., Kolbe, D., Casavant, T., et al. (2018). Genomic Landscape and Mutational Signatures of Deafness-Associated Genes. Am J Hum Genet 103, 484–497. 10.1016/j.ajhg.2018.08.006.

89. Beaudoin, M., Lo, K.S., N’Diaye, A., Rivas, M.A., Dubé, M.P., Laplante, N., Phillips, M.S., Rioux, J.D., Tardif, J.C., and Lettre, G. (2012). Pooled DNA resequencing of 68 myocardial infarction candidate genes in French canadians. Circ Cardiovasc Genet 5, 547–554. 10.1161/circgenetics.112.963165.

90. Dewar, L.J., Alcaide, M., Fornika, D., D’Amato, L., Shafaatalab, S., Stevens, C.M., Balachandra, T., Phillips, S.M., Sanatani, S., Morin, R.D., and Tibbits, G.F. (2017). Investigating the Genetic Causes of Sudden Unexpected Death in Children Through Targeted Next-Generation Sequencing Analysis. Circ Cardiovasc Genet 10. 10.1161/circgenetics.116.001738.

91. Eisenberger, T., Di Donato, N., Decker, C., Delle Vedove, A., Neuhaus, C., Nürnberg, G., Toliat, M., Nürnberg, P., Mürbe, D., and Bolz, H.J. (2018). A C-terminal nonsense mutation links PTPRQ with autosomal-dominant hearing loss, DFNA73. Genet Med 20, 614–621. 10.1038/gim.2017.155.

92. Ryan, J.J., Kalscheur, M., Dellefave, L., McNally, E., and Archer, S.L. (2012). A KCNE1 missense variant (V47I) causing exercise-induced long QT syndrome (Romano Ward). Int J Cardiol 156, e33–35. 10.1016/j.ijcard.2011.08.022.

93. Sand, P.G., Luettich, A., Kleinjung, T., Hajak, G., and Langguth, B. (2010). An Examination of KCNE1 Mutations and Common Variants in Chronic Tinnitus. Genes (Basel) 1, 23–37. 10.3390/genes1010023.

94. Yang, Y., and Sigworth, F.J. (1998). Single-channel properties of IKs potassium channels. J Gen Physiol 112, 665–678. 10.1085/jgp.112.6.665.

95. Bianchi, L., Shen, Z., Dennis, A.T., Priori, S.G., Napolitano, C., Ronchetti, E., Bryskin, R., Schwartz, P.J., and Brown, A.M. (1999). Cellular dysfunction of LQT5-minK mutants: abnormalities of IKs, IKr and trafficking in long QT syndrome. Hum Mol Genet 8, 1499–1507. 10.1093/hmg/8.8.1499.

96. Kanchi, K.L., Johnson, K.J., Lu, C., McLellan, M.D., Leiserson, M.D., Wendl, M.C., Zhang, Q., Koboldt, D.C., Xie, M., Kandoth, C., et al. (2014). Integrated analysis of germline and somatic variants in ovarian cancer. Nat Commun 5, 3156. 10.1038/ncomms4156.

97. Leonenko, G., Richards, A.L., Walters, J.T., Pocklington, A., Chambert, K., Al Eissa, M.M., Sharp, S.I., O’Brien, N.L., Curtis, D., Bass, N.J., et al. (2017). Mutation intolerant genes and targets of FMRP are enriched for nonsynonymous alleles in schizophrenia. Am J Med Genet B Neuropsychiatr Genet 174, 724–731. 10.1002/ajmg.b.32560.

98. Raju, H., Ware, J.S., Skinner, J.R., Hedley, P.L., Arno, G., Love, D.R., van der Werf, C., Tfelt-Hansen, J., Winkel, B.G., Cohen, M.C., et al. (2019). Next-generation sequencing using microfluidic PCR enrichment for molecular autopsy. BMC Cardiovasc Disord 19, 174. 10.1186/s12872-019-1154-8.

99. Krumerman, A., Gao, X., Bian, J.S., Melman, Y.F., Kagan, A., and McDonald, T.V. (2004). An LQT mutant minK alters KvLQT1 trafficking. Am J Physiol Cell Physiol 286, C1453–1463. 10.1152/ajpcell.00275.2003.

100. Silva, J.R., Pan, H., Wu, D., Nekouzadeh, A., Decker, K.F., Cui, J., Baker, N.A., Sept, D., and Rudy, Y. (2009). A multiscale model linking ion-channel molecular dynamics and electrostatics to the cardiac action potential. Proc Natl Acad Sci U S A 106, 11102–11106. 10.1073/pnas.0904505106.

101. Um, S.Y., and McDonald, T.V. (2007). Differential association between HERG and KCNE1 or KCNE2. PLoS One 2, e933. 10.1371/journal.pone.0000933.

102. Vanoye, C.G., Welch, R.C., Tian, C., Sanders, C.R., and George, A.L., Jr. (2010). KCNQ1/KCNE1 assembly, co-translation not required. Channels (Austin) 4, 108–114. 10.4161/chan.4.2.11141.

103. Ma, L., Lin, C., Teng, S., Chai, Y., Bähring, R., Vardanyan, V., Li, L., Pongs, O., and Hui, R. (2003). Characterization of a novel Long QT syndrome mutation G52R-KCNE1 in a Chinese family. Cardiovasc Res 59, 612–619. 10.1016/s0008-6363(03)00510-8.

104. Ruklisa, D., Ware, J.S., Walsh, R., Balding, D.J., and Cook, S.A. (2015). Bayesian models for syndrome- and gene-specific probabilities of novel variant pathogenicity. Genome Med 7, 5. 10.1186/s13073-014-0120-4.

105. Castiglione, A., Hornyik, T., Wülfers, E.M., Giammarino, L., Edler, I., Jowais, J.J., Rieder, M., Perez-Feliz, S., Koren, G., Bősze, Z., et al. (2022). Docosahexaenoic acid normalizes QT interval in long QT type 2 transgenic rabbit models in a genotype-specific fashion. Europace 24, 511–522. 10.1093/europace/euab228.

106. Harmer, S.C., Wilson, A.J., Aldridge, R., and Tinker, A. (2010). Mechanisms of disease pathogenesis in long QT syndrome type 5. Am J Physiol Cell Physiol 298, C263–273. 10.1152/ajpcell.00308.2009.

107. Hoffmann, O.I., Kerekes, A., Lipták, N., Hiripi, L., Bodo, S., Szaloki, G., Klein, S., Ivics, Z., Kues, W.A., and Bosze, Z. (2016). Transposon-Based Reporter Marking Provides Functional Evidence for Intercellular Bridges in the Male Germline of Rabbits. PLoS One 11, e0154489. 10.1371/journal.pone.0154489.

108. Hornyik, T., Castiglione, A., Franke, G., Perez-Feliz, S., Major, P., Hiripi, L., Koren, G., Bősze, Z., Varró, A., Zehender, M., et al. (2020). Transgenic LQT2, LQT5, and LQT2-5 rabbit models with decreased repolarisation reserve for prediction of drug-induced ventricular arrhythmias. Br J Pharmacol 177, 3744–3759. 10.1111/bph.15098.

109. Major, P., Baczkó, I., Hiripi, L., Odening, K.E., Juhász, V., Kohajda, Z., Horváth, A., Seprényi, G., Kovács, M., Virág, L., et al. (2016). A novel transgenic rabbit model with reduced repolarization reserve: long QT syndrome caused by a dominant-negative mutation of the KCNE1 gene. Br J Pharmacol 173, 2046–2061. 10.1111/bph.13500.

110. Morokuma, J., Blackiston, D., and Levin, M. (2008). KCNQ1 and KCNE1 K+ channel components are involved in early left-right patterning in Xenopus laevis embryos. Cell Physiol Biochem 21, 357–372. 10.1159/000129628.

111. Wrobel, E., Rothenberg, I., Krisp, C., Hundt, F., Fraenzel, B., Eckey, K., Linders, J.T., Gallacher, D.J., Towart, R., Pott, L., et al. (2016). KCNE1 induces fenestration in the Kv7.1/KCNE1 channel complex that allows for highly specific pharmacological targeting. Nat Commun 7, 12795. 10.1038/ncomms12795.

112. Cheng, J., Kyle, J.W., Lang, D., Wiedmeyer, B., Guo, J., Yin, K., Huang, L., Vaidyanathan, R., Su, T., and Makielski, J.C. (2017). An East Asian Common Variant Vinculin P.Asp841His Was Associated With Sudden Unexplained Nocturnal Death Syndrome in the Chinese Han Population. J Am Heart Assoc 6. 10.1161/jaha.116.005330.

113. Liu, C., Zhao, Q., Su, T., Tang, S., Lv, G., Liu, H., Quan, L., and Cheng, J. (2013). Postmortem molecular analysis of KCNQ1, KCNH2, KCNE1 and KCNE2 genes in sudden unexplained nocturnal death syndrome in the Chinese Han population. Forensic Sci Int 231, 82–87. 10.1016/j.forsciint.2013.04.020.

114. Melman, Y.F., Krumerman, A., and McDonald, T.V. (2002). A single transmembrane site in the KCNE-encoded proteins controls the specificity of KvLQT1 channel gating. J Biol Chem 277, 25187–25194. 10.1074/jbc.M200564200.

115. Choi, S.H., Jurgens, S.J., Haggerty, C.M., Hall, A.W., Halford, J.L., Morrill, V.N., Weng, L.C., Lagerman, B., Mirshahi, T., Pettinger, M., et al. (2021). Rare Coding Variants Associated With Electrocardiographic Intervals Identify Monogenic Arrhythmia Susceptibility Genes: A Multi-Ancestry Analysis. Circ Genom Precis Med 14, e003300. 10.1161/circgen.120.003300.

116. Haverfield, E.V., Esplin, E.D., Aguilar, S.J., Hatchell, K.E., Ormond, K.E., Hanson-Kahn, A., Atwal, P.S., Macklin-Mantia, S., Hines, S., Sak, C.W., et al. (2021). Physician-directed genetic screening to evaluate personal risk for medically actionable disorders: a large multi-center cohort study. BMC Med 19, 199. 10.1186/s12916-021-01999-2.

117. Knight, L.M., Miller, E., Kovach, J., Arscott, P., von Alvensleben, J.C., Bradley, D., Valdes, S.O., Ware, S.M., Meyers, L., Travers, C.D., et al. (2020). Genetic testing and cascade screening in pediatric long QT syndrome and hypertrophic cardiomyopathy. Heart Rhythm 17, 106–112. 10.1016/j.hrthm.2019.06.015.

118. Tyson, J., Tranebjaerg, L., Bellman, S., Wren, C., Taylor, J.F., Bathen, J., Aslaksen, B., Sørland, S.J., Lund, O., Malcolm, S., et al. (1997). IsK and KvLQT1: mutation in either of the two subunits of the slow component of the delayed rectifier potassium channel can cause Jervell and Lange-Nielsen syndrome. Hum Mol Genet 6, 2179–2185. 10.1093/hmg/6.12.2179.

119. Huang, L., Bitner-Glindzicz, M., Tranebjaerg, L., and Tinker, A. (2001). A spectrum of functional effects for disease causing mutations in the Jervell and Lange-Nielsen syndrome. Cardiovasc Res 51, 670–680. 10.1016/s0008-6363(01)00350-9.

120. Christiansen, M., Hedley, P.L., Theilade, J., Stoevring, B., Leren, T.P., Eschen, O., Sørensen, K.M., Tybjærg-Hansen, A., Ousager, L.B., Pedersen, L.N., et al. (2014). Mutations in Danish patients with long QT syndrome and the identification of a large founder family with p.F29L in KCNH2. BMC Med Genet 15, 31. 10.1186/1471-2350-15-31.

121. Genovese, G., Fromer, M., Stahl, E.A., Ruderfer, D.M., Chambert, K., Landén, M., Moran, J.L., Purcell, S.M., Sklar, P., Sullivan, P.F., et al. (2016). Increased burden of ultra-rare protein-altering variants among 4,877 individuals with schizophrenia. Nat Neurosci 19, 1433–1441. 10.1038/nn.4402.

122. Ghouse, J., Have, C.T., Weeke, P., Bille Nielsen, J., Ahlberg, G., Balslev-Harder, M., Appel, E.V., Skaaby, T., Olesen, S.P., Grarup, N., et al. (2015). Rare genetic variants previously associated with congenital forms of long QT syndrome have little or no effect on the QT interval. Eur Heart J 36, 2523–2529. 10.1093/eurheartj/ehv297.

123. Zhang, J., Juhl, C.R., Hylten-Cavallius, L., Salling-Olsen, M., Linneberg, A., Holst, J.J., Hansen, T., Kanters, J.K., and Torekov, S.S. (2020). Gain-of-function mutation in the voltage-gated potassium channel gene KCNQ1 and glucose-stimulated hypoinsulinemia -case report. BMC Endocr Disord 20, 38. 10.1186/s12902-020-0513-x.

124. Rodriguez-Flores, J.L., Fakhro, K., Hackett, N.R., Salit, J., Fuller, J., Agosto-Perez, F., Gharbiah, M., Malek, J.A., Zirie, M., Jayyousi, A., et al. (2014). Exome sequencing identifies potential risk variants for Mendelian disorders at high prevalence in Qatar. Hum Mutat 35, 105–116. 10.1002/humu.22460.

125. Coll, M., Ferrer-Costa, C., Pich, S., Allegue, C., Rodrigo, E., Fernández-Fresnedo, G., Barreda, P., Mates, J., Martinez de Francisco, A.L., Ortega, I., et al. (2018). Role of genetic and electrolyte abnormalities in prolonged QTc interval and sudden cardiac death in end-stage renal disease patients. PLoS One 13, e0200756. 10.1371/journal.pone.0200756.

126. Lin, Y., Williams, N., Wang, D., Coetzee, W., Zhou, B., Eng, L.S., Um, S.Y., Bao, R., Devinsky, O., McDonald, T.V., et al. (2017). Applying High-Resolution Variant Classification to Cardiac Arrhythmogenic Gene Testing in a Demographically Diverse Cohort of Sudden Unexplained Deaths. Circ Cardiovasc Genet 10. 10.1161/circgenetics.117.001839.

127. Wang, D., Shah, K.R., Um, S.Y., Eng, L.S., Zhou, B., Lin, Y., Mitchell, A.A., Nicaj, L., Prinz, M., McDonald, T.V., et al. (2014). Cardiac channelopathy testing in 274 ethnically diverse sudden unexplained deaths. Forensic Sci Int 237, 90–99. 10.1016/j.forsciint.2014.01.014.

128. Wright, C.F., West, B., Tuke, M., Jones, S.E., Patel, K., Laver, T.W., Beaumont, R.N., Tyrrell, J., Wood, A.R., Frayling, T.M., et al. (2019). Assessing the Pathogenicity, Penetrance, and Expressivity of Putative Disease-Causing Variants in a Population Setting. Am J Hum Genet 104, 275–286. 10.1016/j.ajhg.2018.12.015.

129. Li, Y., Zaydman, M.A., Wu, D., Shi, J., Guan, M., Virgin-Downey, B., and Cui, J. (2011). KCNE1 enhances phosphatidylinositol 4,5-bisphosphate (PIP2) sensitivity of IKs to modulate channel activity. Proc Natl Acad Sci U S A 108, 9095–9100. 10.1073/pnas.1100872108.

130. Akhavanfard, S., Padmanabhan, R., Yehia, L., Cheng, F., and Eng, C. (2020). Comprehensive germline genomic profiles of children, adolescents and young adults with solid tumors. Nat Commun 11, 2206. 10.1038/s41467-020-16067-1.

131. Blom, L.J., Visser, M., Christiaans, I., Scholten, M.F., Bootsma, M., van den Berg, M.P., Yap, S.C., van der Heijden, J.F., Doevendans, P.A., Loh, P., et al. (2019). Incidence and predictors of implantable cardioverter-defibrillator therapy and its complications in idiopathic ventricular fibrillation patients. Europace 21, 1519–1526. 10.1093/europace/euz151.

132. Li, S., Zhang, C., Liu, N., Bai, H., Hou, C., and Pu, J. (2019). Clinical implications of sarcomere and nonsarcomere gene variants in patients with left ventricular noncompaction cardiomyopathy. Mol Genet Genomic Med 7, e874. 10.1002/mgg3.874.

133. Li, S., Zhang, C., Liu, N., Bai, H., Hou, C., Wang, J., Song, L., and Pu, J. (2018). Genotype-Positive Status Is Associated With Poor Prognoses in Patients With Left Ventricular Noncompaction Cardiomyopathy. J Am Heart Assoc 7, e009910. 10.1161/jaha.118.009910.

134. Liebrechts-Akkerman, G., Liu, F., van Marion, R., Dinjens, W.N.M., and Kayser, M. (2020). Explaining sudden infant death with cardiac arrhythmias: Complete exon sequencing of nine cardiac arrhythmia genes in Dutch SIDS cases highlights new and known DNA variants. Forensic Sci Int Genet 46, 102266. 10.1016/j.fsigen.2020.102266.

135. Marcondes, L., Crawford, J., Earle, N., Smith, W., Hayes, I., Morrow, P., Donoghue, T., Graham, A., Love, D., and Skinner, J.R. (2018). Long QT molecular autopsy in sudden unexplained death in the young (1-40 years old): Lessons learnt from an eight year experience in New Zealand. PLoS One 13, e0196078. 10.1371/journal.pone.0196078.

136. Middha, S., Lindor, N.M., McDonnell, S.K., Olson, J.E., Johnson, K.J., Wieben, E.D., Farrugia, G., Cerhan, J.R., and Thibodeau, S.N. (2015). How well do whole exome sequencing results correlate with medical findings? A study of 89 Mayo Clinic Biobank samples. Front Genet 6, 244. 10.3389/fgene.2015.00244.

137. Muin, D.A., Kollmann, M., Blatterer, J., Hoermann, G., Husslein, P.W., Lafer, I., Petek, E., and Schwarzbraun, T. (2021). Cardio-pathogenic variants in unexplained intrauterine fetal death: a retrospective pilot study. Sci Rep 11, 6737. 10.1038/s41598-021-85893-0.

138. Skinner, J.R., Crawford, J., Smith, W., Aitken, A., Heaven, D., Evans, C.A., Hayes, I., Neas, K.R., Stables, S., Koelmeyer, T., et al. (2011). Prospective, population-based long QT molecular autopsy study of postmortem negative sudden death in 1 to 40 year olds. Heart Rhythm 8, 412–419. 10.1016/j.hrthm.2010.11.016.

139. van Lint, F.H.M., Mook, O.R.F., Alders, M., Bikker, H., Lekanne Dit Deprez, R.H., and Christiaans, I. (2019). Large next-generation sequencing gene panels in genetic heart disease: yield of pathogenic variants and variants of unknown significance. Neth Heart J 27, 304–309. 10.1007/s12471-019-1250-5.

140. Mellor, G., Laksman, Z.W.M., Tadros, R., Roberts, J.D., Gerull, B., Simpson, C.S., Klein, G.J., Champagne, J., Talajic, M., Gardner, M., et al. (2017). Genetic Testing in the Evaluation of Unexplained Cardiac Arrest: From the CASPER (Cardiac Arrest Survivors With Preserved Ejection Fraction Registry). Circ Cardiovasc Genet 10. 10.1161/circgenetics.116.001686.

141. Peeters, U., Scornik, F., Riuró, H., Pérez, G., Komurcu-Bayrak, E., Van Malderen, S., Pappaert, G., Tarradas, A., Pagans, S., Daneels, D., et al. (2015). Contribution of Cardiac Sodium Channel β-Subunit Variants to Brugada Syndrome. Circ J 79, 2118–2129. 10.1253/circj.CJ-15-0164.

142. Ackerman, M.J., Tester, D.J., Jones, G.S., Will, M.L., Burrow, C.R., and Curran, M.E. (2003). Ethnic differences in cardiac potassium channel variants: implications for genetic susceptibility to sudden cardiac death and genetic testing for congenital long QT syndrome. Mayo Clin Proc 78, 1479–1487. 10.4065/78.12.1479.

143. Rocheleau, J.M., Gage, S.D., and Kobertz, W.R. (2006). Secondary structure of a KCNE cytoplasmic domain. J Gen Physiol 128, 721–729. 10.1085/jgp.200609657.

144. Steffensen, A.B., Refaat, M.M., David, J.P., Mujezinovic, A., Calloe, K., Wojciak, J., Nussbaum, R.L., Scheinman, M.M., and Schmitt, N. (2015). High incidence of functional ion-channel abnormalities in a consecutive Long QT cohort with novel missense genetic variants of unknown significance. Sci Rep 5, 10009. 10.1038/srep10009.

145. Jimmy, J.J., Chen, C.Y., Yeh, H.M., Chiu, W.Y., Yu, C.C., Liu, Y.B., Tsai, C.T., Lo, L.W., Yeh, S.F., and Lai, L.P. (2014). Clinical characteristics of patients with congenital long QT syndrome and bigenic mutations. Chin Med J (Engl) 127, 1482–1486.

146. Kothiyal, P., Cox, S., Ebert, J., Husami, A., Kenna, M.A., Greinwald, J.H., Aronow, B.J., and Rehm, H.L. (2010). High-throughput detection of mutations responsible for childhood hearing loss using resequencing microarrays. BMC Biotechnol 10, 10. 10.1186/1472-6750-10-10.

147. Lai, L.P., Su, Y.N., Chiang, F.T., Juang, J.M., Liu, Y.B., Ho, Y.L., Chen, W.J., Yeh, S.J., Wang, C.C., Ko, Y.L., et al. (2005). Denaturing high-performance liquid chromatography screening of the long QT syndrome-related cardiac sodium and potassium channel genes and identification of novel mutations and single nucleotide polymorphisms. J Hum Genet 50, 490–496. 10.1007/s10038-005-0283-3.

148. Barro-Soria, R., Rebolledo, S., Liin, S.I., Perez, M.E., Sampson, K.J., Kass, R.S., and Larsson, H.P. (2014). KCNE1 divides the voltage sensor movement in KCNQ1/KCNE1 channels into two steps. Nat Commun 5, 3750. 10.1038/ncomms4750.

149. Liin, S.I., Larsson, J.E., Barro-Soria, R., Bentzen, B.H., and Larsson, H.P. (2016). Fatty acid analogue N-arachidonoyl taurine restores function of I(Ks) channels with diverse long QT mutations. Elife 5. 10.7554/eLife.20272.

150. Cereda, M., Gambardella, G., Benedetti, L., Iannelli, F., Patel, D., Basso, G., Guerra, R.F., Mourikis, T.P., Puccio, I., Sinha, S., et al. (2016). Patients with genetically heterogeneous synchronous colorectal cancer carry rare damaging germline mutations in immune-related genes. Nat Commun 7, 12072. 10.1038/ncomms12072.

151. Kenna, K.P., McLaughlin, R.L., Hardiman, O., and Bradley, D.G. (2013). Using reference databases of genetic variation to evaluate the potential pathogenicity of candidate disease variants. Hum Mutat 34, 836–841. 10.1002/humu.22303.

152. Li, M.H., Abrudan, J.L., Dulik, M.C., Sasson, A., Brunton, J., Jayaraman, V., Dugan, N., Haley, D., Rajagopalan, R., Biswas, S., et al. (2015). Utility and limitations of exome sequencing as a genetic diagnostic tool for conditions associated with pediatric sudden cardiac arrest/sudden cardiac death. Hum Genomics 9, 15. 10.1186/s40246-015-0038-y.

153. Marschall, C., Moscu-Gregor, A., and Klein, H.G. (2019). Variant panorama in 1,385 index patients and sensitivity of expanded next-generation sequencing panels in arrhythmogenic disorders. Cardiovasc Diagn Ther 9, S292–s298. 10.21037/cdt.2019.06.06.

154. Splawski, I., Tristani-Firouzi, M., Lehmann, M.H., Sanguinetti, M.C., and Keating, M.T. (1997). Mutations in the hminK gene cause long QT syndrome and suppress IKs function. Nat Genet 17, 338–340. 10.1038/ng1197-338.

155. Chen, J., Liu, Z., Creagh, J., Zheng, R., and McDonald, T.V. (2020). Physical and functional interaction sites in cytoplasmic domains of KCNQ1 and KCNE1 channel subunits. Am J Physiol Heart Circ Physiol 318, H212–h222. 10.1152/ajpheart.00459.2019.

156. McCrossan, Z.A., Roepke, T.K., Lewis, A., Panaghie, G., and Abbott, G.W. (2009). Regulation of the Kv2.1 potassium channel by MinK and MiRP1. J Membr Biol 228, 1–14. 10.1007/s00232-009-9154-8.

157. Seebohm, G., Strutz-Seebohm, N., Ureche, O.N., Henrion, U., Baltaev, R., Mack, A.F., Korniychuk, G., Steinke, K., Tapken, D., Pfeufer, A., et al. (2008). Long QT syndrome-associated mutations in KCNQ1 and KCNE1 subunits disrupt normal endosomal recycling of IKs channels. Circ Res 103, 1451–1457. 10.1161/circresaha.108.177360.

158. Sesti, F., and Goldstein, S.A. (1998). Single-channel characteristics of wild-type IKs channels and channels formed with two minK mutants that cause long QT syndrome. J Gen Physiol 112, 651–663. 10.1085/jgp.112.6.651.

159. Strutz-Seebohm, N., Seebohm, G., Fedorenko, O., Baltaev, R., Engel, J., Knirsch, M., and Lang, F. (2006). Functional coassembly of KCNQ4 with KCNE-beta-subunits in Xenopus oocytes. Cell Physiol Biochem 18, 57–66. 10.1159/000095158.

160. Harmonizing Clinical Sequencing and Interpretation for the eMERGE III Network. (2019). Am J Hum Genet 105, 588–605. 10.1016/j.ajhg.2019.07.018.

161. Frequency of genomic secondary findings among 21,915 eMERGE network participants. (2020). Genet Med 22, 1470–1477. 10.1038/s41436-020-0810-9.

162. Behr, E.R., Ritchie, M.D., Tanaka, T., Kääb, S., Crawford, D.C., Nicoletti, P., Floratos, A., Sinner, M.F., Kannankeril, P.J., Wilde, A.A., et al. (2013). Genome wide analysis of drug-induced torsades de pointes: lack of common variants with large effect sizes. PLoS One 8, e78511. 10.1371/journal.pone.0078511.

163. Bowling, K.M., Thompson, M.L., Gray, D.E., Lawlor, J.M.J., Williams, K., East, K.M., Kelley, W.V., Moss, I.P., Absher, D.M., Partridge, E.C., et al. (2021). Identifying rare, medically relevant variation via population-based genomic screening in Alabama: opportunities and pitfalls. Genet Med 23, 280–288. 10.1038/s41436-020-00976-z.

164. Burashnikov, E., Pfeiffer, R., Barajas-Martinez, H., Delpón, E., Hu, D., Desai, M., Borggrefe, M., Häissaguerre, M., Kanter, R., Pollevick, G.D., et al. (2010). Mutations in the cardiac L-type calcium channel associated with inherited J-wave syndromes and sudden cardiac death. Heart Rhythm 7, 1872–1882. 10.1016/j.hrthm.2010.08.026.

165. Cuneo, B.F., Etheridge, S.P., Horigome, H., Sallee, D., Moon-Grady, A., Weng, H.Y., Ackerman, M.J., and Benson, D.W. (2013). Arrhythmia phenotype during fetal life suggests long-QT syndrome genotype: risk stratification of perinatal long-QT syndrome. Circ Arrhythm Electrophysiol 6, 946–951. 10.1161/circep.113.000618.

166. Diñeiro, M., Cifuentes, G.A., Capín, R., Santiago, A., Otero, A., Castillo, D., Pruneda, P.C., Ordóñez, G.R., Cabanillas, R., and Cadiñanos, J. (2020). Sequencing results from multiple individuals of different ethnicities strongly question the existence of the KCNE1B pseudogene. Eur J Hum Genet 28, 401–402. 10.1038/s41431-019-0502-6.

167. Downie, L., Halliday, J., Burt, R., Lunke, S., Lynch, E., Martyn, M., Poulakis, Z., Gaff, C., Sung, V., Wake, M., et al. (2020). Exome sequencing in infants with congenital hearing impairment: a population-based cohort study. Eur J Hum Genet 28, 587–596. 10.1038/s41431-019-0553-8.

168. Duggal, P., Vesely, M.R., Wattanasirichaigoon, D., Villafane, J., Kaushik, V., and Beggs, A.H. (1998). Mutation of the gene for IsK associated with both Jervell and Lange-Nielsen and Romano-Ward forms of Long-QT syndrome. Circulation 97, 142–146. 10.1161/01.cir.97.2.142.

169. Kanter, R.J., Pfeiffer, R., Hu, D., Barajas-Martinez, H., Carboni, M.P., and Antzelevitch, C. (2012). Brugada-like syndrome in infancy presenting with rapid ventricular tachycardia and intraventricular conduction delay. Circulation 125, 14–22. 10.1161/circulationaha.111.054007.

170. Murdock, D.R., Venner, E., Muzny, D.M., Metcalf, G.A., Murugan, M., Hadley, T.D., Chander, V., de Vries, P.S., Jia, X., Hussain, A., et al. (2021). Genetic testing in ambulatory cardiology clinics reveals high rate of findings with clinical management implications. Genet Med 23, 2404–2414. 10.1038/s41436-021-01294-8.

171. Noman, M., Ishaq, R., Bukhari, S.A., Ahmed, Z.M., and Riazuddin, S. (2019). Delineation of Homozygous Variants Associated with Prelingual Sensorineural Hearing Loss in Pakistani Families. Genes (Basel) 10. 10.3390/genes10121031.

172. Petri, H., Witting, N., Ersbøll, M.K., Sajadieh, A., Dunø, M., Helweg-Larsen, S., Vissing, J., Køber, L., and Bundgaard, H. (2014). High prevalence of cardiac involvement in patients with myotonic dystrophy type 1: a cross-sectional study. Int J Cardiol 174, 31–36. 10.1016/j.ijcard.2014.03.088.

173. Shashi, V., Schoch, K., Spillmann, R., Cope, H., Tan, Q.K., Walley, N., Pena, L., McConkie-Rosell, A., Jiang, Y.H., Stong, N., et al. (2019). A comprehensive iterative approach is highly effective in diagnosing individuals who are exome negative. Genet Med 21, 161–172. 10.1038/s41436-018-0044-2.

174. Tang, D., Fakiola, M., Syn, G., Anderson, D., Cordell, H.J., Scaman, E.S.H., Davis, E., Miles, S.J., McLeay, T., Jamieson, S.E., et al. (2018). Arylsulphatase A Pseudodeficiency (ARSA-PD), hypertension and chronic renal disease in Aboriginal Australians. Sci Rep 8, 10912. 10.1038/s41598-018-29279-9.

175. Tester, D.J., Arya, P., Will, M., Haglund, C.M., Farley, A.L., Makielski, J.C., and Ackerman, M.J. (2006). Genotypic heterogeneity and phenotypic mimicry among unrelated patients referred for catecholaminergic polymorphic ventricular tachycardia genetic testing. Heart Rhythm 3, 800–805. 10.1016/j.hrthm.2006.03.025.

176. Tester, D.J., Will, M.L., Haglund, C.M., and Ackerman, M.J. (2005). Compendium of cardiac channel mutations in 541 consecutive unrelated patients referred for long QT syndrome genetic testing. Heart Rhythm 2, 507–517. 10.1016/j.hrthm.2005.01.020.

177. Vergult, S., Dauber, A., Delle Chiaie, B., Van Oudenhove, E., Simon, M., Rihani, A., Loeys, B., Hirschhorn, J., Pfotenhauer, J., Phillips, J.A., 3rd, et al. (2012). 17q24.2 microdeletions: a new syndromal entity with intellectual disability, truncal obesity, mood swings and hallucinations. Eur J Hum Genet 20, 534–539. 10.1038/ejhg.2011.239.

178. Vijayakumar, R., Silva, J.N.A., Desouza, K.A., Abraham, R.L., Strom, M., Sacher, F., Van Hare, G.F., Haïssaguerre, M., Roden, D.M., and Rudy, Y. (2014). Electrophysiologic substrate in congenital Long QT syndrome: noninvasive mapping with electrocardiographic imaging (ECGI). Circulation 130, 1936–1943. 10.1161/circulationaha.114.011359.

179. Weeke, P., Mosley, J.D., Hanna, D., Delaney, J.T., Shaffer, C., Wells, Q.S., Van Driest, S., Karnes, J.H., Ingram, C., Guo, Y., et al. (2014). Exome sequencing implicates an increased burden of rare potassium channel variants in the risk of drug-induced long QT interval syndrome. J Am Coll Cardiol 63, 1430–1437. 10.1016/j.jacc.2014.01.031.

180. Whiffin, N., Roberts, A.M., Minikel, E., Zappala, Z., Walsh, R., O’Donnell-Luria, A.H., Karczewski, K.J., Harrison, S.M., Thomson, K.L., Sage, H., et al. (2019). Using High-Resolution Variant Frequencies Empowers Clinical Genome Interpretation and Enables Investigation of Genetic Architecture. Am J Hum Genet 104, 187–190. 10.1016/j.ajhg.2018.11.012.

181. Zhang, Y., Li, H., Wang, J., Wang, G., Tan, X., and Lei, M. (2020). Generation of three iPSC lines (XACHi007-A, XACHi008-A, XACHi009-A) from a Chinese family with long QT syndrome type 5 with heterozygous c.226G>A (p.D76N) mutation in KCNE1gene. Stem Cell Res 45, 101798. 10.1016/j.scr.2020.101798.

182. Abbott, G.W., and Goldstein, S.A. (2002). Disease-associated mutations in KCNE potassium channel subunits (MiRPs) reveal promiscuous disruption of multiple currents and conservation of mechanism. Faseb j 16, 390–400. 10.1096/fj.01-0520hyp.

183. Abitbol, I., Peretz, A., Lerche, C., Busch, A.E., and Attali, B. (1999). Stilbenes and fenamates rescue the loss of I(KS) channel function induced by an LQT5 mutation and other IsK mutants. Embo j 18, 4137–4148. 10.1093/emboj/18.15.4137.

184. Chen, J., Zheng, R., Melman, Y.F., and McDonald, T.V. (2009). Functional interactions between KCNE1 C-terminus and the KCNQ1 channel. PLoS One 4, e5143. 10.1371/journal.pone.0005143.

185. Chouabe, C., Neyroud, N., Richard, P., Denjoy, I., Hainque, B., Romey, G., Drici, M.D., Guicheney, P., and Barhanin, J. (2000). Novel mutations in KvLQT1 that affect Iks activation through interactions with Isk. Cardiovasc Res 45, 971–980. 10.1016/s0008-6363(99)00411-3.

186. Gage, S.D., and Kobertz, W.R. (2004). KCNE3 truncation mutants reveal a bipartite modulation of KCNQ1 K+ channels. J Gen Physiol 124, 759–771. 10.1085/jgp.200409114.

187. Haitin, Y., Wiener, R., Shaham, D., Peretz, A., Cohen, E.B., Shamgar, L., Pongs, O., Hirsch, J.A., and Attali, B. (2009). Intracellular domains interactions and gated motions of I(KS) potassium channel subunits. Embo j 28, 1994–2005. 10.1038/emboj.2009.157.

188. Hoppe, U.C., Marbán, E., and Johns, D.C. (2001). Distinct gene-specific mechanisms of arrhythmia revealed by cardiac gene transfer of two long QT disease genes, HERG and KCNE1. Proc Natl Acad Sci U S A 98, 5335–5340. 10.1073/pnas.091239098.

189. Kurokawa, J., Chen, L., and Kass, R.S. (2003). Requirement of subunit expression for cAMP-mediated regulation of a heart potassium channel. Proc Natl Acad Sci U S A 100, 2122–2127. 10.1073/pnas.0434935100.

190. Nakamura, H., Kurokawa, J., Bai, C.X., Asada, K., Xu, J., Oren, R.V., Zhu, Z.I., Clancy, C.E., Isobe, M., and Furukawa, T. (2007). Progesterone regulates cardiac repolarization through a nongenomic pathway: an in vitro patch-clamp and computational modeling study. Circulation 116, 2913–2922. 10.1161/circulationaha.107.702407.

191. Terrenoire, C., Clancy, C.E., Cormier, J.W., Sampson, K.J., and Kass, R.S. (2005). Autonomic control of cardiac action potentials: role of potassium channel kinetics in response to sympathetic stimulation. Circ Res 96, e25–34. 10.1161/01.RES.0000160555.58046.9a.

192. Zheng, R., Thompson, K., Obeng-Gyimah, E., Alessi, D., Chen, J., Cheng, H., and McDonald, T.V. (2010). Analysis of the interactions between the C-terminal cytoplasmic domains of KCNQ1 and KCNE1 channel subunits. Biochem J 428, 75–84. 10.1042/bj20090977.

193. Hietikko, E., Kotimäki, J., Okuloff, A., Sorri, M., and Männikkö, M. (2012). A replication study on proposed candidate genes in Ménière’s disease, and a review of the current status of genetic studies. Int J Audiol 51, 841–845. 10.3109/14992027.2012.705900.

194. Riuró, H., Campuzano, O., Berne, P., Arbelo, E., Iglesias, A., Pérez-Serra, A., Coll-Vidal, M., Partemi, S., Mademont-Soler, I., Picó, F., et al. (2015). Genetic analysis, in silico prediction, and family segregation in long QT syndrome. Eur J Hum Genet 23, 79–85. 10.1038/ejhg.2014.54.

195. Kwok, S.Y., Liu, A.P., Chan, C.Y., Lun, K.S., Fung, J.L., Mak, C.C., Chung, B.H., and Yung, T.C. (2018). Clinical and genetic profile of congenital long QT syndrome in Hong Kong: a 20-year experience in paediatrics. Hong Kong Med J 24, 561–570. 10.12809/hkmj187487.

196. Lvov, A., Gage, S.D., Berrios, V.M., and Kobertz, W.R. (2010). Identification of a protein-protein interaction between KCNE1 and the activation gate machinery of KCNQ1. J Gen Physiol 135, 607–618. 10.1085/jgp.200910386.

197. Wu, D.M., Lai, L.P., Zhang, M., Wang, H.L., Jiang, M., Liu, X.S., and Tseng, G.N. (2006). Characterization of an LQT5-related mutation in KCNE1, Y81C: implications for a role of KCNE1 cytoplasmic domain in IKs channel function. Heart Rhythm 3, 1031–1040. 10.1016/j.hrthm.2006.05.022.

198. de Villiers, C.P., van der Merwe, L., Crotti, L., Goosen, A., George, A.L., Jr., Schwartz, P.J., Brink, P.A., Moolman-Smook, J.C., and Corfield, V.A. (2014). AKAP9 is a genetic modifier of congenital long-QT syndrome type 1. Circ Cardiovasc Genet 7, 599–606. 10.1161/circgenetics.113.000580.

199. Hedley, P.L., Durrheim, G.A., Hendricks, F., Goosen, A., Jespersgaard, C., Støvring, B., Pham, T.T., Christiansen, M., Brink, P.A., and Corfield, V.A. (2013). Long QT syndrome in South Africa: the results of comprehensive genetic screening. Cardiovasc J Afr 24, 231–237. 10.5830/cvja-2013-032.

200. Mates, J., Mademont-Soler, I., Del Olmo, B., Ferrer-Costa, C., Coll, M., Pérez-Serra, A., Picó, F., Allegue, C., Fernandez-Falgueras, A., Álvarez, P., et al. (2018). Role of copy number variants in sudden cardiac death and related diseases: genetic analysis and translation into clinical practice. Eur J Hum Genet 26, 1014–1025. 10.1038/s41431-018-0119-1.

201. Coll, M., Striano, P., Ferrer-Costa, C., Campuzano, O., Matés, J., Del Olmo, B., Iglesias, A., Pérez-Serra, A., Mademont, I., Picó, F., et al. (2017). Targeted next-generation sequencing provides novel clues for associated epilepsy and cardiac conduction disorder/SUDEP. PLoS One 12, e0189618. 10.1371/journal.pone.0189618.

202. Guelly, C., Abilova, Z., Nuralinov, O., Panzitt, K., Akhmetova, A., Rakhimova, S., Kozhamkulov, U., Kairov, U., Molkenov, A., Seisenova, A., et al. (2021). Patients with coronary heart disease, dilated cardiomyopathy and idiopathic ventricular tachycardia share overlapping patterns of pathogenic variation in cardiac risk genes. PeerJ 9, e10711. 10.7717/peerj.10711.

203. Hayashi, K., Konno, T., Fujino, N., Itoh, H., Fujii, Y., Imi-Hashida, Y., Tada, H., Tsuda, T., Tanaka, Y., Saito, T., et al. (2016). Impact of Updated Diagnostic Criteria for Long QT Syndrome on Clinical Detection of Diseased Patients: Results From a Study of Patients Carrying Gene Mutations. JACC Clin Electrophysiol 2, 279–287. 10.1016/j.jacep.2016.01.003.

204. Jin, S.C., Dong, W., Kundishora, A.J., Panchagnula, S., Moreno-De-Luca, A., Furey, C.G., Allocco, A.A., Walker, R.L., Nelson-Williams, C., Smith, H., et al. (2020). Exome sequencing implicates genetic disruption of prenatal neuro-gliogenesis in sporadic congenital hydrocephalus. Nat Med 26, 1754–1765. 10.1038/s41591-020-1090-2.

205. Millat, G., Chevalier, P., Restier-Miron, L., Da Costa, A., Bouvagnet, P., Kugener, B., Fayol, L., Gonzàlez Armengod, C., Oddou, B., Chanavat, V., et al. (2006). Spectrum of pathogenic mutations and associated polymorphisms in a cohort of 44 unrelated patients with long QT syndrome. Clin Genet 70, 214–227. 10.1111/j.1399-0004.2006.00671.x.

206. Li, Y., Zhu, B., Su, J., Yin, Y., and Yu, F. (2016). Identification of SLC26A4 mutations p.L582LfsX4, p.I188T and p.E704K in a Chinese family with large vestibular aqueduct syndrome (LVAS). Int J Pediatr Otorhinolaryngol 91, 1–5. 10.1016/j.ijporl.2016.08.026.

207. Asatryan, B., Schaller, A., Seiler, J., Servatius, H., Noti, F., Baldinger, S.H., Tanner, H., Roten, L., Dillier, R., Lam, A., et al. (2019). Usefulness of Genetic Testing in Sudden Cardiac Arrest Survivors With or Without Previous Clinical Evidence of Heart Disease. Am J Cardiol 123, 2031–2038. 10.1016/j.amjcard.2019.02.061.

208. Boles, U., Simpson, C., Gul, E.E., Kiss, C., Enriquez, A., Jia, Z., Baranchuk, A., and Walia, J.S. (2017). Clinical evaluation of R860Q semi-conservative amino acid substitution in CACNA1C gene in association with long QT syndrome. Int J Cardiol Heart Vasc 15, 21–23. 10.1016/j.ijcha.2017.03.006.

209. Refsgaard, L., Holst, A.G., Sadjadieh, G., Haunsø, S., Nielsen, J.B., and Olesen, M.S. (2012). High prevalence of genetic variants previously associated with LQT syndrome in new exome data. Eur J Hum Genet 20, 905–908. 10.1038/ejhg.2012.23.

210. Schulze-Bahr, E., Schwarz, M., Hauenschild, S., Wedekind, H., Funke, H., Haverkamp, W., Breithardt, G., Pongs, O., and Isbrandt, D. (2001). A novel long-QT 5 gene mutation in the C-terminus (V109I) is associated with a mild phenotype. J Mol Med (Berl) 79, 504–509. 10.1007/s001090100249.

211. Waterston, R.H., Lindblad-Toh, K., Birney, E., Rogers, J., Abril, J.F., Agarwal, P., Agarwala, R., Ainscough, R., Alexandersson, M., An, P., et al. (2002). Initial sequencing and comparative analysis of the mouse genome. Nature 420, 520–562. 10.1038/nature01262.

212. Dvir, M., Strulovich, R., Sachyani, D., Ben-Tal Cohen, I., Haitin, Y., Dessauer, C., Pongs, O., Kass, R., Hirsch, J.A., and Attali, B. (2014). Long QT mutations at the interface between KCNQ1 helix C and KCNE1 disrupt I(KS) regulation by PKA and PIP_2_. J Cell Sci 127, 3943–3955. 10.1242/jcs.147033.

213. Borges, M.G., Rocha, C.S., Carvalho, B.S., and Lopes-Cendes, I. (2020). Methodological differences can affect sequencing depth with a possible impact on the accuracy of genetic diagnosis. Genet Mol Biol 43, e20190270. 10.1590/1678-4685-gmb-2019-0270.

214. Cuenca, S., Ruiz-Cano, M.J., Gimeno-Blanes, J.R., Jurado, A., Salas, C., Gomez-Diaz, I., Padron-Barthe, L., Grillo, J.J., Vilches, C., Segovia, J., et al. (2016). Genetic basis of familial dilated cardiomyopathy patients undergoing heart transplantation. J Heart Lung Transplant 35, 625–635. 10.1016/j.healun.2015.12.014.

215. Harmer, S.C., Mohal, J.S., Royal, A.A., McKenna, W.J., Lambiase, P.D., and Tinker, A. (2014). Cellular mechanisms underlying the increased disease severity seen for patients with long QT syndrome caused by compound mutations in KCNQ1. Biochem J 462, 133–142. 10.1042/bj20140425.

